# NERINE reveals rare variant associations in gene networks across phenotypes and implicates an *SNCA-PRL-LRRK2* subnetwork in Parkinson’s disease

**DOI:** 10.1101/2025.01.07.631688

**Authors:** Sumaiya Nazeen, Xinyuan Wang, Autumn R. Morrow, Ronya Strom, Elizabeth Ethier, Dylan Ritter, Alexander B. H. Henderson, Jalwa Afroz, Christopher S. Cassa, Nathan O. Stitziel, Rajat M. Gupta, Kelvin C. Luk, Lorenz Studer, Vikram Khurana, Shamil R. Sunyaev

**Affiliations:** Department of Biomedical Informatics, Harvard Medical School, Boston, MA, USA; Division of Genetics, Brigham and Women’s Hospital and Harvard Medical School, Boston, MA, USA; American Parkinson’s Disease Association Center for Advanced Research, Harvard Biomarkers Study 2.0 and MyTrial Programs, Division of Movement Disorders, Department of Neurology, Brigham and Women’s Hospital and Harvard Medical School, Boston, MA, USA; Broad Institute of MIT and Harvard, Cambridge, MA, USA; The Center for Stem Cell Biology, Sloan-Kettering Institute for Cancer Research, New York, NY, USA; Department of Neurology, Sean M. Healey & AMG Center for ALS, Massachusetts General Hospital and Harvard Medical School, Boston, MA, USA; Cardiovascular Division, John T. Milliken Department of Medicine, Washington University School of Medicine, St. Louis, MO, USA; Department of Genetics, Washington University School of Medicine, St. Louis, MO, USA; Division of Cardiovascular Medicine, Brigham and Women’s Hospital and Harvard Medical School, Boston, MA, USA; Department of Pathology and Laboratory Medicine, Perelman School of Medicine at the University of Pennsylvania, PA, USA; Aligning Science Across Parkinson’s (ASAP) Collaborative Research Network, Chevy Chase, MD, 20815; Harvard Stem Cell Institute, Cambridge, MA, USA

## Abstract

There are two primary approaches to study the genetic basis of human phenotypes. Experiments in model systems generate interpretable gene networks but, in isolation, do not establish relevance to the human condition. Statistical genetics identifies relevant association signals at the variant or gene level but lacks tools to test specific mechanistic models, as existing methods do not incorporate the topology of gene-gene interactions. We bridge these two strategies by introducing a method that competitively tests network hypotheses with rare variant associations. A hierarchical model-based association test NERINE for the first time incorporates gene network topology while remaining resilient to network inaccuracies. We demonstrate NERINE’s ability to test network hypotheses derived from both canonical pathway databases and model system screens. Comprehensive database-wide search of pathway networks with NERINE uncovers compelling associations for breast cancer, cardiovascular diseases, and type II diabetes, which are undetected by single-gene tests. Testing bespoke networks from experimental screens targeting key PD pathologies: dopaminergic neuron survival and *α-synuclein* pathobiology, NERINE highlights rare variant burden in gene modules related to autophagy, vesicle trafficking, and protein homeostasis. Genome-scale CRISPRi-screening of *α-synuclein* toxicity modifiers in human neurons and NERINE converge on *PRL*, revealing an intraneuronal *α-synuclein*/*prolactin* stress response that may impact resilience to PD pathologies.

## Introduction

Advancements in high-throughput screens, such as perturbation assays in model systems, have enabled the generation of vast amounts of data capturing crucial biological knowledge, often represented as gene and protein interaction networks^1–7^. Some networks represent well-characterized metabolic or signaling pathways, while others capture regulatory interactions, physical protein-protein interactions, or protein-DNA associations. More abstractly, genetic interactions, such as co-essentiality, reflect dependencies in gene contributions to cellular fitness and phenotypes. As a result, gene networks serve as fundamental frameworks for understanding polygenic trait variation, including complex disease mechanisms, enabling researchers to formulate testable biological hypotheses about human phenotypes^8–10^. However, establishing causal links between gene networks and human phenotypes requires direct evidence from human genetics.

Rare protein-coding variants offer a direct means of establishing such connections, as these mutations occur within gene bodies and can directly impact gene function^11,12^. When tested at the level of pathways and networks, they can reveal new insights about causal disease mechanisms^13–16^. Despite their potential, no existing rare variant association test leverages gene networks with defined topologies to assess their association with human phenotypes^14–25^. They treat pathways as either a single entity (“mega-gene”) or a simple collection of genes (“bag of genes”), not accounting for the complex interactions between genes and the fact that not all genes in a pathway contribute equally to a phenotype.

To address this gap, we present NERINE (NEtwork-based Rare varIaNt Enrichment), a new statistical framework for rare variant association testing that integrates information on genes and their interactions into a hierarchical model, ensuring robustness against network inaccuracies. NERINE offers several key advantages. First, it allows for the competitive evaluation of network hypotheses, prioritizing network topologies and the experimental assays that define them by their relevance to human phenotypes. Second, it enhances statistical power by aggregating rare variants across functionally connected genes, leading to improved mechanistic insights into the directional effects of genes on a phenotype. Third, it moves beyond broad and overlapping gene classifications, such as gene ontology, by utilizing experimentally derived relationships with defined edge geometries and signed weights, thereby increasing the biological specificity of the burden test. Importantly, fourth, it allows for inaccuracies (e.g., false positive “nodes”) in that network. This framework paves the way for integrative research strategies whereby genetic screens identify network-based hypotheses to test for association with human phenotypes. Networks demonstrating significant associations can subsequently be refined and validated through targeted experimental approaches, bridging the gap between genetic discovery and functional understanding.

We demonstrate two ways of applying NERINE: first, using pre-existing pathway modules to analyze prevalent diseases—breast cancer (BRCA), type II diabetes (T2D), coronary artery disease (CAD), and myocardial infarction (MI) in UK and MGB biobanks; second, leveraging experimentally derived gene networks from model system screens to uncover rare variant associations in a complex neurodegenerative disease like Parkinson’s disease (PD), where existing gene-level rare variant association tests have been thoroughly underpowered with existing cohorts^26,27^.

Comprehensive testing of database pathway modules in large-scale biobanks with NERINE has enabled us to identify several novel associations, including in the estrogen receptor pathway in BRCA, an adipogenesis-related network in T2D, and non-lipid gene networks (e.g., complement system) in CAD and MI. In PD, NERINE tests network hypotheses generated from experimental screens targeting two core pathologies of PD: i) alpha-synuclein (ɑS) proteotoxicity and ii) human dopaminergic (DA) neuron survival. We uncover associations in gene modules linked to autophagy regulation (*HMGB1-OPTN-USP10* module) and vesicle trafficking and protein homeostasis (*LRRK2-PRL-SNCA* module)—findings that all have eluded single-gene tests.

One novel finding by NERINE—an association of rare damaging missense variants at the *PRL* locus encoding *prolactin* (*Prl*) with PD risk—converges with an independent genome-scale functional screen in a CRISPRi-induced synucleinopathy cortical neuron (CiS-CN) model designed for studying physical and genetic modifiers of *α-synuclein* (αS) toxicity in cortical neurons^28^. Subsequent functional validation using the CiS-CN model and a well-established mouse PFF model^29^ uncovers a hitherto unknown intraneuronal role of *PRL* in *α-synuclein* stress response in PD. This underscores the transformative potential of experimental screens and human statistical genetics as mutually reinforcing methodologies to reveal novel disease mechanisms.

## Results

### Modeling rare variant burden in gene networks incorporating edge geometry

We present NERINE, a new statistical framework for assessing the cumulative effect of rare variants in a gene network on a dichotomous phenotype (**Figure 1**). NERINE uniquely incorporates information on network vertices (genes) and network edges (interactions) into a parametric model and is robust with respect to unimportant genes in the network (see **Supplementary Table T1** for distinction from other methods). Many data types can represent interactions between genes and gene products (i.e., transcripts and proteins). Interactions may differ in terms of relationship types (from protein complexes to sets of co-expressed genes to protein-DNA or protein-RNA interactions) and scale (from extensive protein interaction networks to pathways of just a dozen genes). Regardless of the data source, we encode gene-gene relationships using a positive semidefinite matrix, Σ.

**Figure 1.**
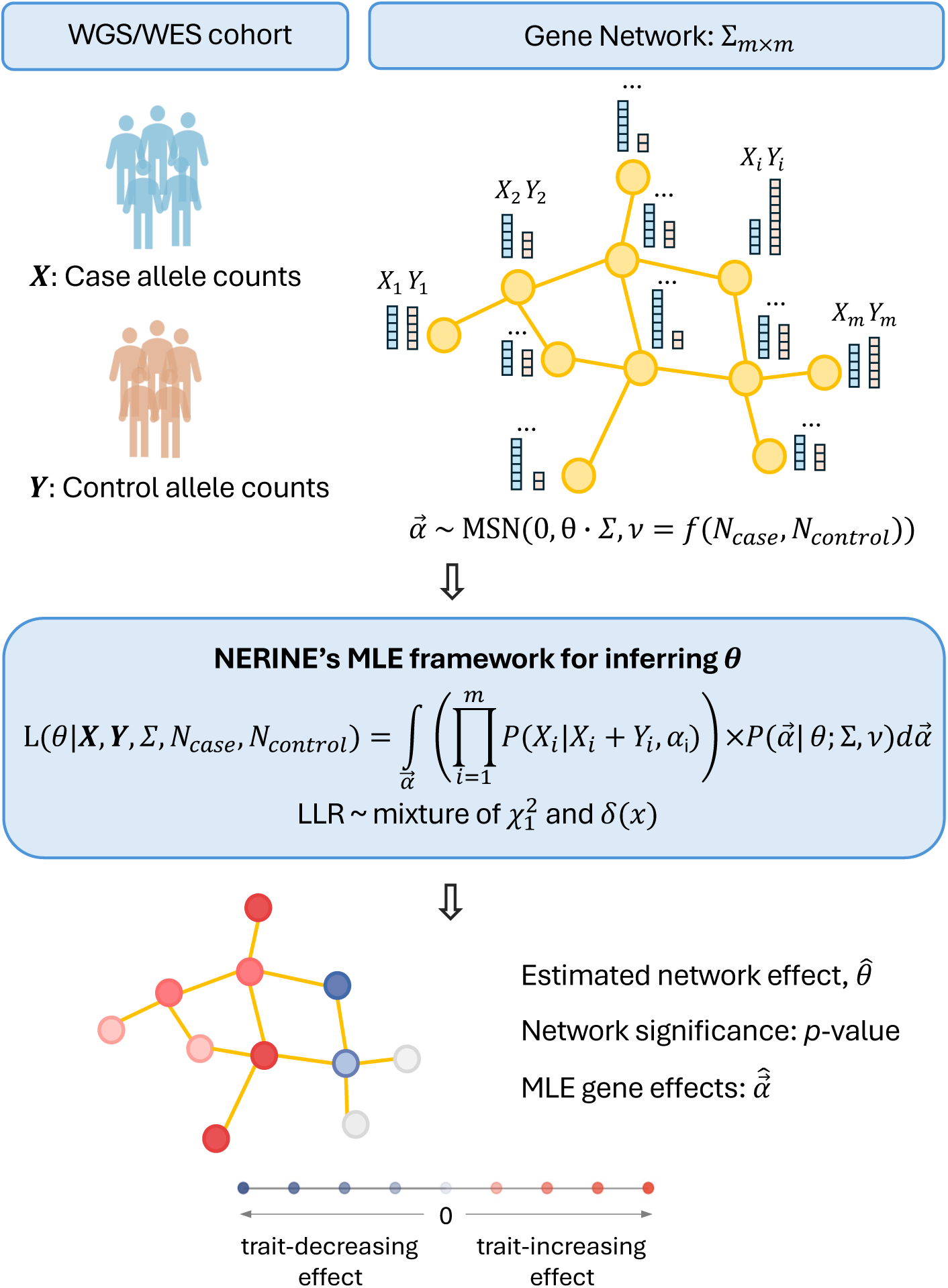
Overview of NERINE: a rare variant association test leveraging gene network topology. NERINE’s hierarchical framework for testing rare variant burden aggregated across a gene network for binary traits. Inputs include allele counts in cases (***X***) and controls (***Y***) from WGS or WES datasets and a gene network encoded by a symmetric positive semidefinite matrix, Σ_D×D_. Σ can represent diverse biological relationships (e.g., physical or genetic interactions, co-expression, co-essentiality, and pathway membership). Gene-effects 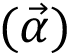 are modeled with a multivariate skew-normal distribution parameterized by Σ, network effect *θ*, and case-control skew *υ*. This parameterization allows network genes to have either zero effect or varying degrees of trait-increasing and decreasing effects. *θ* is inferred using a maximum likelihood estimation (MLE) framework where the likelihood, *L*, is an integral over the product of two terms—(i) the product of the per-gene conditional probability of observing allele count in cases (*X_i_*) given the allele count in the overall cohort (*X_i_* + *Y_i_*) in the network, and (ii) the probability of observing a specific combination of gene effects determined by *θ* and Σ. NERINE performs nested hypothesis testing, with log-likelihood ratio (LLR) being the test statistic, and provides an asymptotic *p*-value from the mixture of the delta function, *δ*(*x*), and a chi-square distribution with one degree of freedom *X_1_*^2^. It also estimates the most likely gene effects 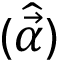 under the estimated 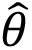 (**Methods**).

We model gene effect sizes, denoted by vector 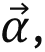 as draws from a multivariate skew-normal distribution 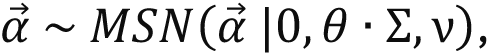 with univariate skew-normal marginals. This hierarchical model captures the biological expectation that functionally related genes exhibit correlated effect sizes (when non-zero) and correlated probabilities of having no phenotypic effect. The parameter *θ* reflects the importance of the gene group and is the object of inference, while the skewness parameter, ν = *f*(*N_case_*, *N_control_*) encodes the case–control imbalance (**Figure 1**, **Methods**). Here, *N_case_* and *N_control_* represent the number of cases and controls, respectively. When case-control groups are perfectly balanced, this model reduces to a multivariate normal, 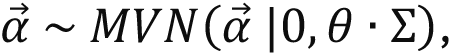 with scalar normal marginals. This parameterization allows gene effect sizes to vary in both magnitude and direction while permitting some to be zero (in case of noisy networks).

NERINE infers *θ* using the maximum likelihood estimation (MLE) framework, where the likelihood is given by,

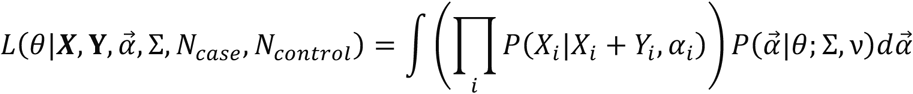

We approximate *L* as a weighted sum over multivariate quadrature points from the domain of integration achieved through a lookup table with pruning (**Methods, Supplementary Note**). To ensure computationally efficient evaluation of the quadrature points, we make several design choices. First, we model rare variant counts in member genes in cases (***X***) and controls (***Y***) as independent Poisson variables, with rates determined by population allele frequencies renormalized between cases and controls by 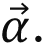 This implies that the conditional probability of observed rare variant count in each gene in cases, given the total rare variant count in the cohort, follows a Binomial distribution (**Supplementary Note**). Next, we apply a custom transformation to map the unbounded univariate skew-normal marginals of 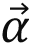 to Beta distributions on [0, 1] (**Supplementary Note**), so that the transformed *α*_i_s can be used as Binomial success probabilities. We denote the transformed gene-effects as 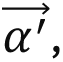 which incorporates skewness (*ν*) by controlling the shape parameters of the Beta distribution (**Supplementary Note**). When the network has no effect on the trait, the expected case–control allele ratio reflects sample size proportions in each gene. When the network influences the trait, allele count ratios deviate from this null expectation in relevant genes. The approximate likelihood is given by,

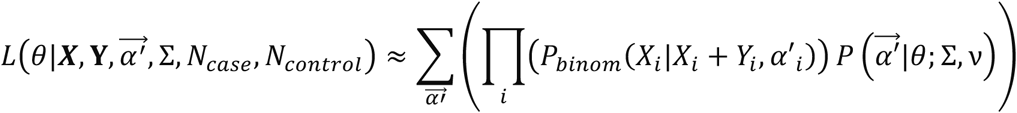

NERINE performs nested hypotheses testing, the test statistic being the log-likelihood ratio (LLR). We test the hypothesis *θ* = 0 vs. *θ* > 0 using the maximum likelihood estimate of *θ* (**Methods**). Since *θ* lies on the boundary of the parameter space, the test statistic, LLR, follows an asymptotic distribution arising from a weighted mixture of a point mass at zero and a chi-square distribution with one degree of freedom^30,31^. We confirmed this asymptotic behavior using extensive simulations with four well-studied canonical pathways (**Supplementary Figure S1** and **S2**), and artificially constructed topologies (**Supplementary Figure S3**) under the null model (*θ* = 0). NERINE provides asymptotic p-values for significant networks (*θ* > 0), and estimates the maximum likelihood effects of member genes under the inferred *θ*. However, gene-level p-values are not provided (**Methods**).

We benchmarked the performance of NERINE against existing rare variant tests in simulations under the alternative hypothesis (*θ* > 0) (**Figure 2**, **Supplementary Figures S4 and S5**, **Methods**). Compared with pathway-level adaptations of such tests, including CMC-Fisher, Fisher minimum p-value, Fisher combined test, SKAT-O, and pathway-based trend test, RVTT (only trait-increasing genes case), NERINE consistently showed higher empirical power, especially in noisy networks (**Figure 2, Supplementary Figures S4 and S5**). Positive control experiments using UK Biobank (UKBB) data on LDL and HDL cholesterol (LDL-C and HDL-C) confirmed NERINE’s ability to detect significant rare variant burdens in key lipid-related pathways in European ancestry-specific (**Supplementary Figures S6, S7, and S8**, **Methods**) as well as pan-ancestry (**Supplementary Figure S9**, **Supplementary Table T2**, **Methods**) analyses. Compared to SKAT-O, NERINE provided lower p-values, demonstrating its ability to leverage network connectivity for improved power (**Supplementary Figure S6C**). NERINE’s estimated effect sizes and directions for individual genes, correctly identified *PCSK9* and *APOB* as reducing LDL cholesterol and *LDLR* as increasing it, consistent with known biology^32^ (**Supplementary Figure S6D**). Similarly, for HDL cholesterol, it linked *ABCA1*, *LCAT*, and *APOA1* to low levels and *CETP*, *LIPC*, *LIPG*, and *SCARB1* to high levels. (**Supplementary Figure S6E**).

**Figure 2.**
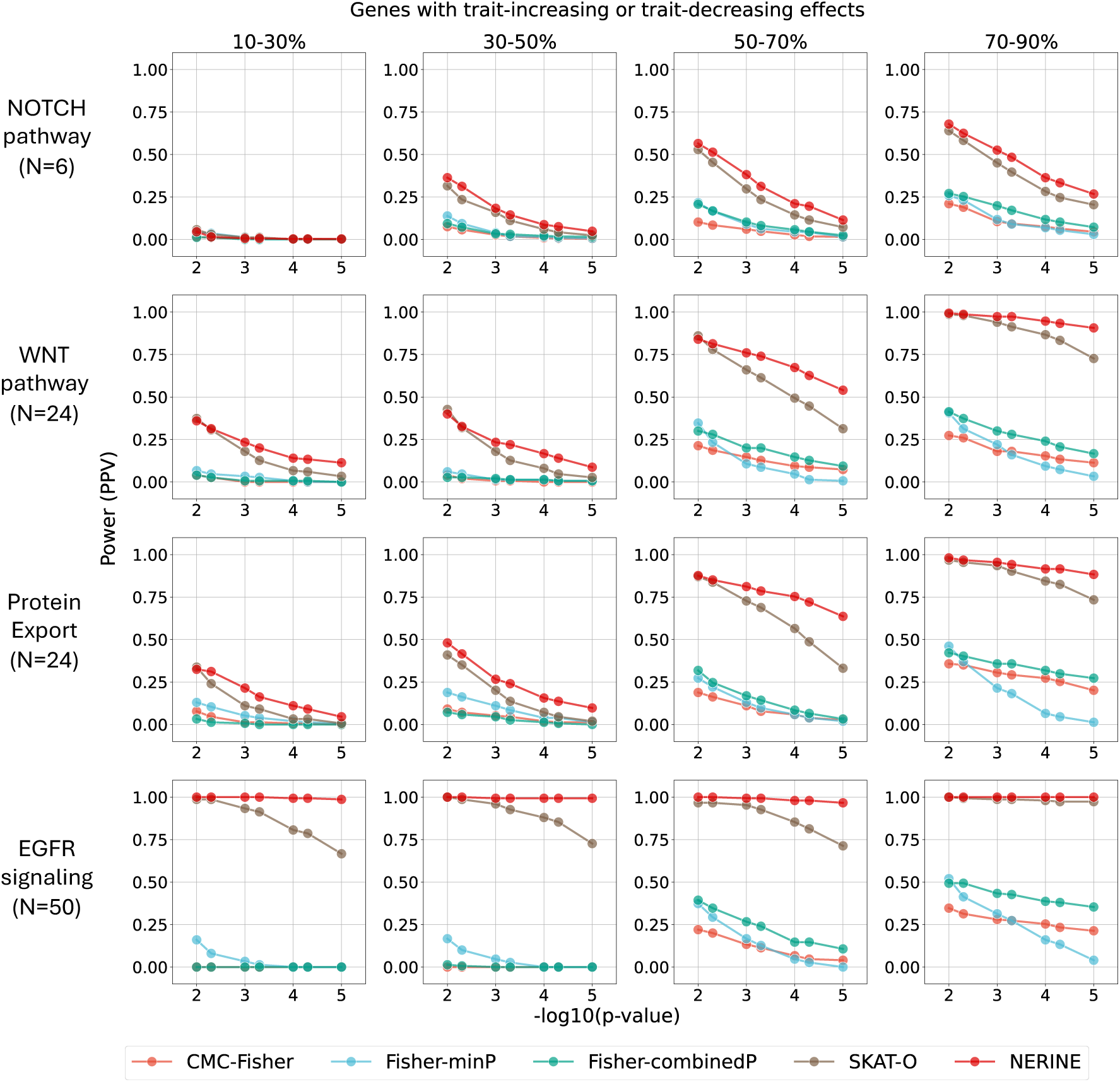
NERINE outperforms existing rare variant association tests in simulations. Power was evaluated using a simulated binary trait (2,000 cases and 2,000 controls) across four canonical pathways (NOTCH pathway, WNT pathway, protein export, and EGFR signaling) with non-zero network effect (*θ* = 0.2). Noise was varied by the proportion of genes with non-zero effects (10–90%), spanning highly noisy (only 10–30% genes with effect) to highly informative (70–90% genes with effect) networks. For each noise profile, 250 iterations were performed per network, and power was measured as the positive predictive value (PPV) at different significance cutoffs (1 × 10^-2^, 5 × 10^-3^, 1 × 10^-3^, 5 × 10^-4^, 1 × 10^-4^, 5 × 10^-5^, and 1 × 10^-5^) (**Methods**). NERINE consistently outperformed existing rare-variant association tests, with the largest gains in noisy settings. All tests were two-sided.

### Selecting the most informative network topology with NERINE

NERINE selects the most informative network topology when multiple alternatives exist for a given gene set, underscoring its ability to leverage the edge geometry of gene networks for more powerful inference. We demonstrated this feature in simulations with (i) four well-characterized pathways of varying sizes (NOTCH, WNT, protein export, and EGFR signaling) for a simulated binary phenotype, and (ii) significant lipid-related pathways on LDL-C and HDL-C phenotypes in UKBB. For the former, NERINE consistently achieved a higher PPV at different p-value cutoffs with “true” networks compared to randomized counterparts, especially in noisy scenarios (**Supplementary Figure S10, Methods**). Similarly, in the latter analysis, NERINE achieved higher log-likelihood ratios and lower p-values with real network topologies compared to random ones (**Supplementary Figure S11, Methods**).

When gene-gene relationships in a network are available from multiple data sources, NERINE selects the optimal topology for estimating the burden of rare variants in a network, as showcased using lipid-related pathway gene networks for LDL-C and HDL-C in UKBB (**Supplementary Table T3**; Methods). For lipid-related pathway gene sets, we applied NERINE on network topologies constructed from three data sources --- (i) high confidence experimentally derived physical and genetic interactions of proteins from canonical databases, (ii) gene-gene co-expression in liver tissue in GTEx (v8), and (iii) co-essentiality of genes in liver cell lines in DepMap3 (v2023Q2) (**Methods**). Our results revealed that the data source providing optimal topology was highly dependent on the specific gene set under investigation (**Supplementary Table T3**). **Figure 2A** shows an example scenario where one data source provided more relevant topological information than the others. When comparing individuals with low HDL-C to those with high HDL-C in UKBB, liver tissue co-expression emerged as the most informative source for the core module of HDL-cholesterol-related genes (Bonf. p-value = 1.34e-73). Example scenarios where data sources, other than co-expression, captured the edge relationships of specific gene sets more accurately are shown in **Supplementary Figure S12**. For instance, physical and genetic interactions best described the VLDLR internalization and degradation pathway module associated with the LDL-cholesterol phenotype in UKBB (Bonf. p-value = 2.50e-31). Meanwhile, co-essentiality in liver cell lines provided the most relevant edge geometry for the pathway of LDL, HDL, and triglyceride metabolism (Bonf. p-value = 1.58e-46). These findings underscore a crucial insight: different data sources encode distinct and complementary aspects of gene relationships. NERINE could effectively exploit this phenomenon to select the most informative data source.

Applying NERINE across tissue-specific co-expression networks spanning all 52 tissue types from GTEx (v8) further reinforced its ability to identify the most informative biological context. Specifically, liver tissue emerged as the most relevant context for core HDL-cholesterol-related genes, where NERINE a high estimated network effect of *θ* = 0.8 with the strongest statistical support (Bonf. p-value = 1.34e-73; **Figure 3B**, **Supplementary Table T4**). This result highlights a key advantage of NERINE—its capacity to leverage the variability observed in edge relationships and their strengths across different tissue contexts for the same set of input genes, thereby selecting the optimal tissue context with biological interpretability.

**Figure 3.**
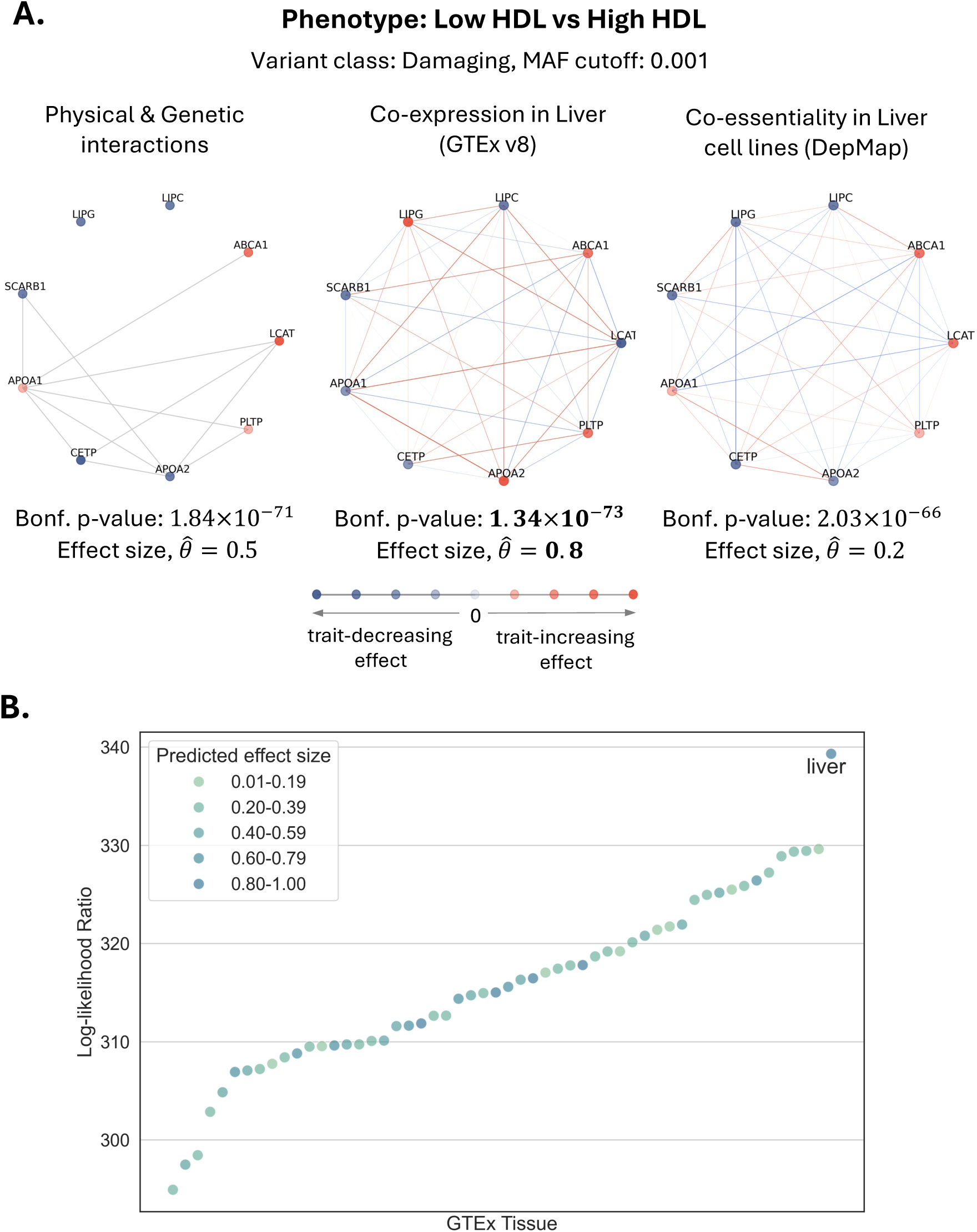
NERINE selects the most informative network topology. **A.** NERINE evaluates competing network topologies and selects the most informative one for a given gene set and phenotype. For core HDL-related genes, co-expression in liver (Bonf. *p* = 1.34 × 10^-73^; effect size, 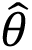 = 0.8) outperforms PPI and liver co-essentiality networks when testing for rare damaging variant burden in low vs. high HDL-C individuals (*N*_case_ = 26,800; *N_control_* = 27,178) in UKBB (**Methods**). Here, node colors denote the direction of gene effects (orange: trait-increasing, and purple: trait-decreasing), with intensity reflecting magnitude. Co-expression and co-essentiality network edges indicate correlation (red: positive, and blue: negative), with edge thickness reflecting the strength of the correlation. PPI network edges represent binary relationships and are colored gray. **B.** Across 52 GTEx (v8) tissues, NERINE identifies liver as the most informative context for core HDL-related genes when testing for rare damaging variant association with the binarized HDL-C phenotype in UKBB, as in **A** (**Methods**). NERINE yields 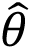 = 0.8, the highest LLR (339.32), and strongest significance (Bonf. *p* = 1.34 × 10^-73^) using the liver co-expression network.

### Identifying rare variant associations in common diseases across canonical pathway networks

NERINE enables comprehensive rare variant burden testing in gene networks for common diseases using biobank-scale data from the UK Biobank (UKBB) and the Mass General Brigham Biobank (MGBBB) (**Figure 4A**, **Methods**). Applied to breast cancer (BRCA), type II diabetes (T2D), coronary artery disease (CAD), and early-onset myocardial infarction (MI)—which impact 3-13% of the population—NERINE identified significant rare loss-of-function (LoF) burdens in seven pathways for BRCA, one for T2D, two for CAD, and fourteen for MI (**Figure 4B**, **Supplementary Table T5, Supplementary Figures S13–S15**). Other than LoF variants, significant burdens were also detected in the damaging and damaging missense categories (**Supplementary Table T5**), with no enrichment of neutral or synonymous variants. Notably, networks with significant burden in the BRCA phenotype overlapped with biologically plausible pathways, such as cancer susceptibility (i.e., DNA repair, cell cycle regulation, apoptosis) and hormone signaling, while cardiovascular disease networks highlighted lipid metabolism, immune response, blood coagulation, and apoptosis (**Supplementary Figures S13–S15, Supplementary Tables T6 and T7**), indicating true-positive findings. NERINE-predicted directional effects of genes in significant pathways per disease phenotype are shown in **Supplementary Tables T8-T11.**

**Figure 4.**
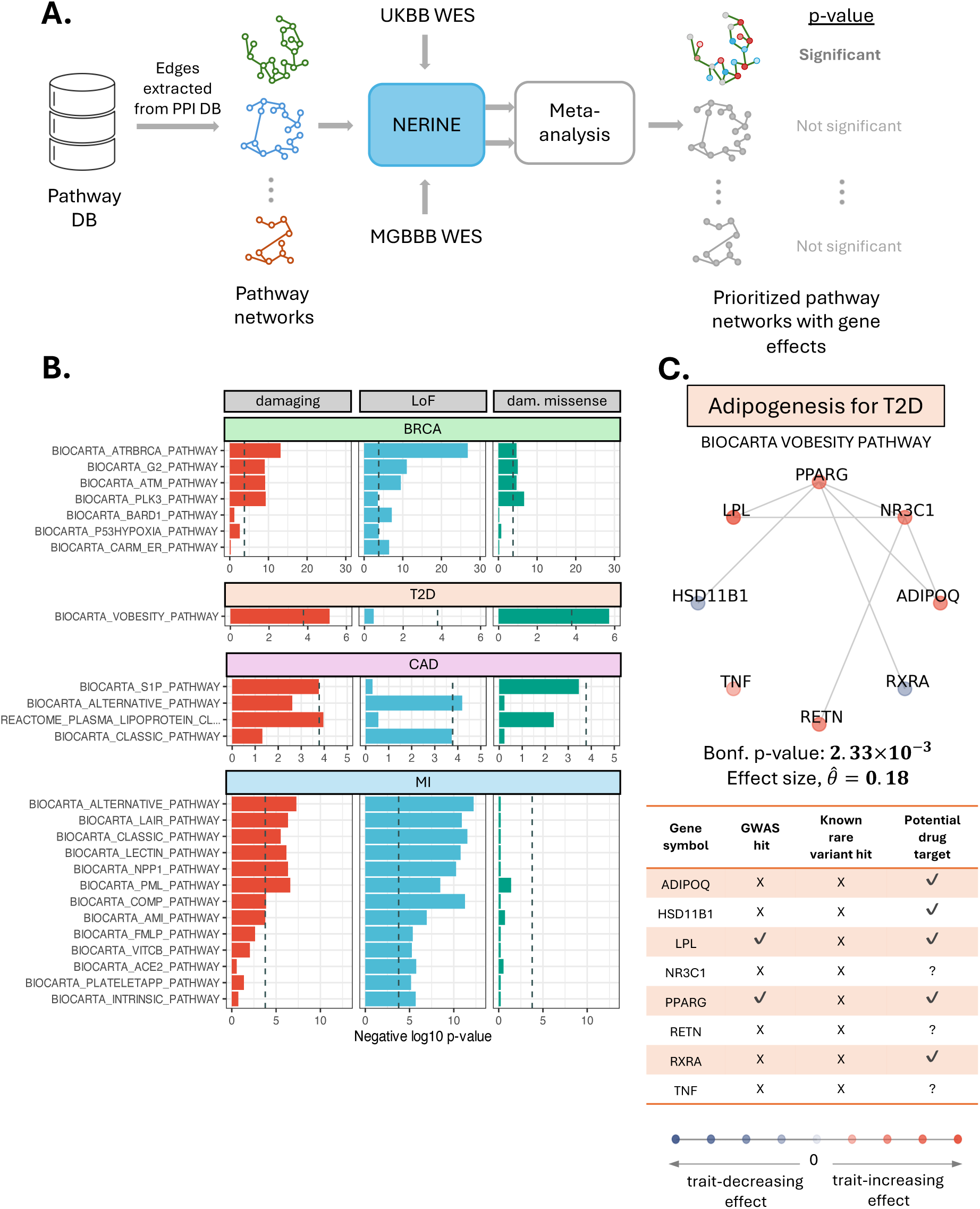
NERINE identifies rare variant burden in canonical pathway networks across common diseases in UKBB and MGBBB. **A.** NERINE’s application to case-control WES cohorts in UKBB and MGBBB for rare variant association across database pathway networks. NERINE tests 306 pathway networks (derived from curated physical/genetic interactions) and six variant classes (LoF, damaging missense, damaging, missense, synonymous, neutral missense) (**Methods**). Case-control cohort sizes (*N_case_*/*N_control_*): BRCA (UKBB: 10,648/91,886; MGBBB: 1,113/2,459), T2D (UKBB: 22,502/68,370; MGBBB: 747/2,188), CAD (UKBB: 4,561/12,321; MGBBB: 902/1,488), and MI (UKBB: 2,521/5,012; MGBBB: 326/2,068). Two-sided *p*-values were meta-analyzed via Fisher’s combined test and Bonferroni-corrected with Nyholt’s adjustment (**Methods**). **B.** Bonferroni-significant disease pathways with rare variant burden in LoF, damaging missense, and damaging categories are shown. Negative log-transformed Fisher’s combined *p*-values are reported; dashed gray line indicates the Bonferroni threshold of 0.05. **C.** Adipogenesis pathway (BIOCARTA VOBESITY PATHWAY) shows significant rare damaging variant-burden in T2D (avg. 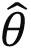 = 0.18, Bonf. *p* = 2.33 × 10^-3^) and serves as a representative example of NERINE’s output. Top: NERINE-predicted gene effects (averaged across cohorts) across the network nodes (orange: trait-increasing; blue: trait-decreasing; intensity reflects magnitude). Bottom: Information on gene-phenotype association in germline genetics from the GWAS catalog (https://www.ebi.ac.uk/gwas/), Genebass^33^, SAIGE-GENE+^19^, and disease-specific literature^34–36^. Network genes, *RXRA*, *PPARG*, *ADIPOQ*, *TNF*, *NR3C1*, *LPL*, *RETN*, and *HSD11B1* were not identified in prior rare variant studies; only *PPARG* and *LPL* in GWAS (bottom). Five of eight genes are under investigation as T2D drug targets^37–43^, highlighting NERINE’s ability to identify therapeutically relevant pathways.

Unlike single-gene tests, which had previously identified only a handful of rare variant associations in these diseases (Genebass^33^— BRCA: 6, T2D: 3, CAD: 1, and MI: 1), NERINE captured aggregated effects of rare variants across gene networks, estimating directional contributions of member genes to putative disease pathways. For instance, in the T2D phenotype, we detected a significant burden of rare damaging variants (i.e., LoF and predicted damaging missense) in the adipogenesis pathway (i.e., BIOCARTA VOBESITY PATHWAY, 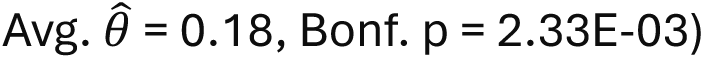 (**Figure 4C**, **Supplementary Tables T5 and T9**), highlighting the potential roles of *RXRA*, *PPARG*, *ADIPOQ*, *TNF*, *NR3C1*, *LPL*, *RETN*, and *HSD11B1*—genes largely unrecognized in previous rare variant studies^19,33^ (**Figure 4C**, **Supplementary Table T9**). Only *PPARG* and *LPL* were implicated in common variant GWAS studies^34–36^. Sensitivity analysis showed that while the inclusion of *LPL* was critical for the database-wide significant signal in this network, the inclusion of *PPARG* was not (**Supplementary Figure S16**). Five of the genes had been studied as drug targets before^37–43^ (**Figure 4C**). Notably, NERINE’s findings align with the fact that the deficiency of *adiponectin*—a protein encoded by *ADIPOQ*—increase insulin resistance^39^ and the risk of T2D, suggesting *ADIPOQ* as a potential candidate for therapeutic investigation. Additionally, NERINE’s prediction of trait-decreasing roles of damaging variants in *RXRA* and *HSD11B1* supports their inhibition as possible therapeutic strategies for T2D^40–43^, paralleling the example of *PCSK9* in cardiovascular disease where protective LoF and damaging missense variants guided effective therapy development^44,45^. However, further experiments are required to distinguish the roles of predicted damaging missenses from LoF variants contributing towards the observed trait-decreasing effect. While NERINE identified rare damaging variant burden in *TNF*, *NR3C1*, and *RETN* in the adipogenesis network, their mechanistic roles in T2D remain understudied, highlighting NERINE’s ability to suggest potential gene-disease associations for further investigation.

### Examples of new network-level associations in breast cancer and cardiovascular diseases

NERINE not only established rare variant connections between new gene networks and phenotypes but also highlighted how rare functional variants, aggregated at the gene level, could influence the phenotype within the network context. For instance, while the estrogen receptor pathway has been clinically studied for breast cancer, rare variant evidence for member genes beyond *BRCA1* has been limited^46,47^. NERINE identified a significant LoF burden at the network level, predicting how putative LoF variants in member genes might have exerted either trait-increasing or trait-decreasing effects on the phenotype. As expected, rare LoF variants in *BRCA1* were associated with an increased risk of BRCA (**Figure 5A: left**; 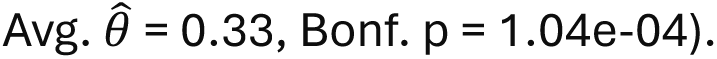 However, after removing the effect of *BRCA1* LoF variants, the network still showed a nominally significant association with BRCA in UKBB, suggesting albeit small but non-zero contributions of other member genes (**Supplementary Figure S17**).

**Figure 5.**
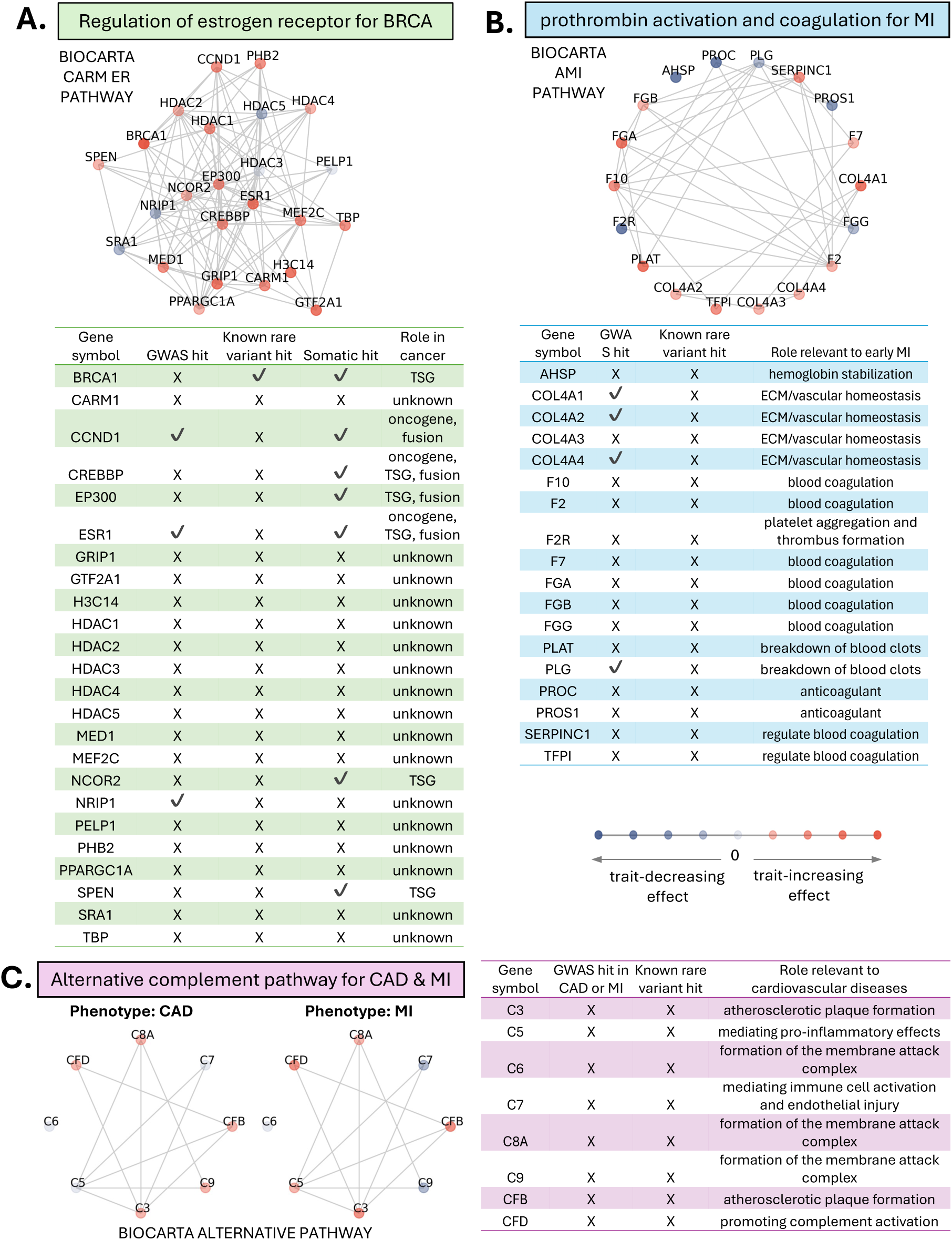
Examples of NERINE’s novel findings in breast cancer and cardiovascular diseases. **A.** Estrogen receptor regulation (BIOCARTA CARM ER PATHWAY) shows significant LoF burden in BRCA (avg. 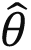 = 0.33, Bonf. *p* = 1.04 × 10^-4^). Top: NERINE-predicted gene effects (averaged across biobanks). Bottom: Information on gene-phenotype association from somatic and germline genetics with functional annotations. **B.** Intrinsic prothrombin activation and coagulation network (BIOCARTA AMI PATHWAY) shows significant LoF burden in MI (avg. 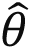 = 0.18, Bonf. *p* = 3.50 × 10^-5^). Top: NERINE-predicted gene effects (averaged across biobanks). Bottom: Information on gene-phenotype association from germline genetics with functional annotations. **C.** Alternative complement system network (BIOCARTA ALTERNATIVE PATHWAY) shows significant LoF burden in CAD (Left; avg. 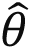 = 0.03, Bonf. *p* = 1.94 × 10^-2^), and MI (Middle; avg. 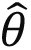 = 0.30, Bonf. *p* = 1.76 × 10^-10^). Right: Information on gene-phenotype association from germline genetics with functional annotations. Across panels, node color represents the direction of gene effect (orange: trait-increasing; purple: trait-decreasing; intensity reflects magnitude). Information sources: GWAS associations from GWAS catalog (https://www.ebi.ac.uk/gwas/); rare variant associations from Genebass^33^, SAIGE-GENE+^19^, and disease-specific studies^47,53,54^; functional annotations for estrogen receptor regulation pathway members from COSMIC (https://cancer.sanger.ac.uk/cosmic); functional annotations of network genes from SynGO (https://www.syngoportal.org/).

Among the estrogen receptor pathway members, 17 were classified as neither tumor suppressor genes (TSGs) nor oncogenes (**Figure 5A: right**). Within the context of this network, NERINE newly linked rare LoF variants in putative GWAS genes, *ESR1* and *CCND1*, to increased BRCA risk. Additionally, it shed light on the effects of loss-of-function of genes, which had unknown oncogenic roles and no genetic evidence. LoF variants in *TBP*, *GRIP1*, *GTF2A1*, *HDAC1*, *MED1*, *MEF2C*, and *PHB2* showed trait-increasing effects, while *HDAC5*, *NRIP1*, and *SRA1* loss-of-function had trait-decreasing effects (**Supplementary Table T8**). Indeed, *PHB2* (*prohibitin 2*) has recently emerged as a key biomarker and therapeutic target in BRCA because of its potential as a tumor suppressor through several possible mechanisms, including inhibition of estrogen receptor-alpha (*ERα*), interaction with *BIG3* (*brefeldin A-inhibited guanine nucleotide-exchange protein 3*), and promotion of *PIG3* (*p53-inducible gene 3*) transcription^48–50^. Furthermore, NERINE’s prediction of *HDAC5* loss-of-function variants decreasing the risk of BRCA aligns with the fact that the inhibition of *HDAC5* induces intrinsic apoptosis in human breast cancer cells, thereby exerting an anti-neoplastic effect^51,52^, providing rationale for follow-up investigations for therapeutic potential.

Rare variant studies in cardiovascular diseases had been underpowered to uncover novel mechanisms beyond lipid-related pathways despite sequencing of over 100,000 subjects in multiple biobanks^33,53^. Here, we identified a significant enrichment for rare variants in several non-lipid pathway gene networks (**Figure 4B**, **Supplementary Figures S14 and S15**) whose member genes had not been identified in single-gene burden analyses^33^. One such pathway of note in MI is the intrinsic prothrombin activation and coagulation cascade (i.e., BIOCARTA AMI PATHWAY), with a significant LoF variant burden 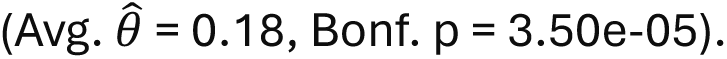 This module consists of several genes associated with collagen and vascular basement membrane integrity (**Figure 5B: left, Supplementary Table T11**). Among the members, extracellular matrix proteins (i.e. *COL4A1, COL4A2, COL4A4*), and clotting factor (*PLG*) had previously been linked to myocardial infarction by common variant GWAS (**Figure 5B: right**) and in vivo studies in knockout mice^5,54–57^. NERINE provided more direct rare variant evidence for this gene module in MI.

Interestingly, NERINE identified an enrichment of rare LoF variants in the alternative complement pathway for both CAD 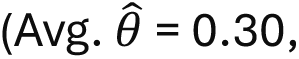 Bonf. p = 1.76e-10), a connection not previously established through human genetics (**Figure 5C**). Functional studies in murine models linked complement components, *C3* and *C5*, to inflammation, endothelial dysfunction, and atherosclerosis^58–60^. In the network, we observed trait-increasing effects for *C3* LoF variants in both CAD and MI and *C5* LoFs in MI. Additionally, rare LoF variants in complement factors *C8A*, *CFB*, and *CFD* had trait-increasing effects on both conditions (**Supplementary Tables T10 and T11**). These directionalities can help reconcile the growing experimental evidence demonstrating that complement activation products in human atherosclerotic plaques can exhibit both protective and pro-atherogenic properties^60^. While the complement signal in cardiovascular diseases is intriguing, the observed differences between MGBBB and UKBB results might be because MGBBB participants had a higher frequency of cardiac events, with thrombosis and inflammation, compared to those in UKBB.

### Bridging genetics and experimental biology with NERINE in Parkinson’s disease

NERINE provides a robust framework for integrating insights from human genetics, cell systems, and model organisms simultaneously to facilitate the study of complex diseases. We illustrate this with Parkinson’s disease (PD), a complex age-related neurodegenerative disorder for which there are no disease-modifying therapies, but for which there is a growing number of genomic and biologic datasets^2,16,28,61–63^ ideally suited for testing with NERINE. While GWAS have identified hundreds of PD-associated loci^64,65^ and linkage analyses^66–68^ have uncovered Mendelian rare variants in key genes—including *SNCA* (encoding αS itself), *GBA1*, and *LRRK2*—existing rare-variant association tests struggle to identify additional causal loci with replication across multiple independent cohorts^26,27^. For idiopathic PD, which accounts for ∼80-85% of cases^69^, the role of rare variants remained poorly understood. Traditional pathway-based analyses^70^ have implicated broad PD mechanisms, including vesicle trafficking, G-protein–coupled receptor (GPCR) signaling, and immune pathways. However, limited resolution and statistical power, due to aggregation of signal across many genes, prevent researchers from determining individual gene contributions or how genes interact to influence disease processes. Thus, such aggregated pathway-or ontology-level signal are less informative for generating experimentally testable hypothesis in specific cellular contexts.

It is possible to assemble the same canonical gene sets into molecular networks with well-defined edge geometries and apply NERINE to pinpoint specific genes and modules for targeted experimentation. We demonstrate this by applying NERINE to gene ontology (GO) biological process modules significantly enriched in PD GWAS-associated genes^64,65,71^ (**Methods**, **Supplementary Figure S18A**, **Supplementary Table T12**). Using database edge-based topologies, NERINE identified a significant burden of rare LoF variants in the *peptidyl-threonine modification* co-essentiality module in two independent PD case-control cohorts from UKBB and AMP-PD 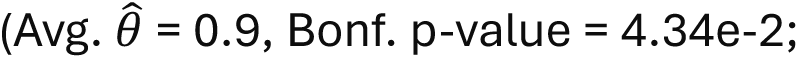 **Supplementary Figure 18B**, **Supplementary Table T13**). Within this module, NERINE suggested trait-increasing LoF burdens in *MCCC1* and *DYRK1A*, and trait-decreasing LoF effects in several other genes, including *FYN*, *USP8*, and *LRRK2* (**Supplementary Table T14**; see **Supplementary Note** for detailed discussion).

### Experimentally derived networks to capture core PD pathologies

As a complementary approach, we turned to biologically informed networks constructed from large-scale genetic data in cellular and model-organism studies. These networks represent experimentally validated assemblies of genes with defined functional relationships, offering a degree of biological specificity not captured by canonical pathways or ontologies. NERINE can pinpoint gene-and module-level effect sizes within these networks, enabling targeted follow-up experiments. Specifically, we analyzed networks comprising genes that modulate *α-synuclein* toxicity and mistrafficking or support dopaminergic (DA) neuron survival, representing the two pathological hallmarks of Parkinson’s disease (PD): *α-synuclein* aggregation in Lewy bodies and DA neuron loss^72,73^

(**Figure 6A**; **Methods**). We assessed these networks in independent PD case-control cohorts from UKBB and AMP-PD.

**Figure 6.**
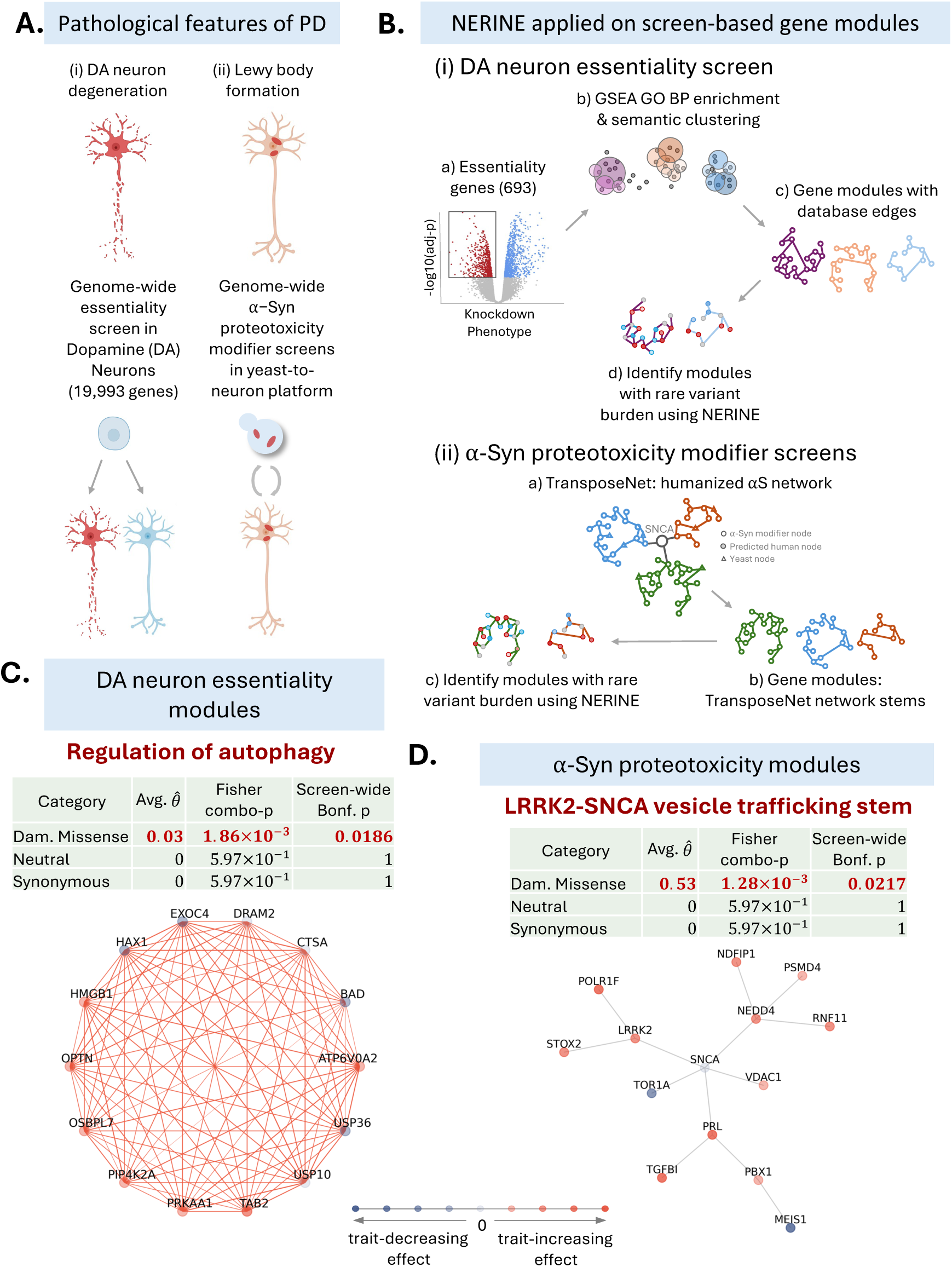
NERINE identifies rare-variant burden in bespoke PD networks from model-system screens. **A.** Key pathological features of PD—Lewy body formation and dopaminergic (DA) neuron loss—targeted via (i) DA neuron essentiality and (ii) *α*-*synuclein* (*αS*) proteotoxicity screens. **B.** Network hypotheses generation from model system screens (**Methods**). Network hypotheses were derived from (i) DA neuron essentiality genes (10 GO BP modules) and (ii) *αS*-toxicity modifier genes (17 TransposeNet stems). Topologies for DA essentiality modules were generated using PPI, substantia nigra co-expression (GTEx v8), and CNS co-essentiality (DepMap v2023Q2) data. For *αS* proteotoxicity network stems, TransposeNet topologies^2^ were used. **C.** NERINE identified screen-wide Bonferroni-significant burden of rare damaging missense variants across AMP-PD (2,117 cases and 1,095 controls) and UKBB-extreme (2,237 cases and 2,553 controls) cohorts in the *HMGB1-OPTN-*containing *autophagy regulation* module (avg. 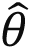 = 0.03, Bonf. *p* = 1.86 × 10^-3^), with substantia nigra co-expression as the optimal topology. **D.** The *LRRK2-SNCA-*containing *vesicle tra[icking and protein homeostasis* stem showed screen-wide Bonferroni-significant burden of rare damaging missense variants (avg. 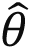 = 0.53, Bonf. *p* = 1.28 × 10^-3^) across AMP-PD and UKBB-sporadic cohorts (AMP-PD: 2,117/1,095; UKBB: 2,237/167,188). In **C** and **D**, node color represents the direction of gene effect (orange: trait-increasing; purple: trait-decreasing; intensity reflects magnitude). In **C**, edge color indicates the sign of the correlation (red: positive; blue: negative), while edge width reflects the correlation strength. In **D**, TransposeNet edges represent binary relationships and are colored in gray.

### Rare variant burden in DA neuron essentiality genes involved in autophagy regulation

Using a genome-scale CRISPR screen of 19,993 genes (**Figure 6B; top**), we identified 693 essential genes for DA neuron survival with significant enrichment in ten GO modules, covering a wide range of biological processes, including autophagy regulation, mRNA processing, apoptosis, and cellular response to DNA damage (**Supplementary Table T15**, **Methods**). We generated networks from these ontological modules by grouping semantically similar GO biological process terms enriched in essentiality genes and then extracting the edge relationships of genes within each group from PPI, co-expression, and co-essentiality databases (**Methods**). NERINE identified a significant rare damaging variant burden the gene module linked to the *regulation of autophagy* 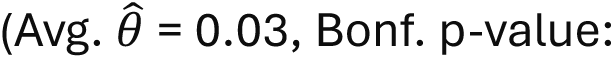 Bonf. p-value: 1.86e-2; **Figure 6C**, **Supplementary Figure S19**, **Supplementary Tables T16 and T17**) in sporadic PD cases compared to controls in the AMP-PD sporadic and UKBB extreme (median age of controls at recruitment >= median age of cases) cohorts (**Methods**). NERINE predicted trait-increasing effects for damaging missense and damaging variants in key genes such as *HMGB1* and *OPTN* in PD (**Figure 6C**, **Supplementary Figure S19**, **Supplementary Tables T17**). The impairment of *HMGB1* function may inhibit autophagy and promote αS accumulation^74^, while the disruption of *OPTN* function may result in the improper clearance of damaged mitochondria (or mitophagy), leading to neurodegeneration^75^. NERINE also identified a trait-increasing burden of rare damaging variants in *USP10* (**Supplementary Figure S19**, **Supplementary Table T17**) whose inactivation may disrupt αS-containing aggresome formation^76^, exacerbating toxicity. These findings readily suggest a small number of potential candidate genes and testable hypotheses supported by orthogonal human genetic evidence, justifying in-depth future experimentation.

### Rare variant burden in a LRRK2-and SNCA-containing αS proteotoxicity module related to vesicle tra[icking and protein homeostasis

Next, we applied NERINE on an αS proteotoxicity network assembled through “yeast-to-neuron” discovery screens^2^ targeting the “αS accumulation in Lewy bodies” feature of PD (**Figure 6B: bottom**). The αS network was generated using the TransposeNet methodology that connected hits from multiple genome-scale deletion and over-expression screens against ɑS proteotoxicity in yeast into an optimal topology by transposing yeast genes into the human interactome space and adding “predicted” genetic nodes as needed. TransposeNet introduced numerous human genes, without any known homologs in yeast, including *SNCA* (that encodes ɑS itself) and *LRRK2*, two genes that are among the most definitively connected directly to PD risk in GWAS and Mendelian linkage analyses^64–68^.

Importantly, the network, validated in human iPSC cortical and DA neurons^2,16^, converged with a proteome-scale proximity labeling screen for αS in neurons^63^. It spanned many relevant pathways, including vesicle trafficking, mRNA metabolism and translation, mitophagy, oxidative metabolism, calcium/NFAT signaling, Toll-like receptor signaling, and purine metabolism (**Supplementary Tables T18**).

Among the TransposeNet αS network branches (**Supplementary Figure S20**), we observed a screen-wide significant burden of rare damaging missense variants in the *LRRK2-* and *SNCA*-containing *vesicle tra[icking and protein homeostasis*-related gene module associated with PD risk in AMP-PD and UKBB 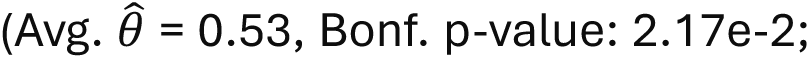 **Figure 6D**, **Supplementary Figure S21**, **Supplementary Tables T18 and T19**). None of the member genes, except *LRRK2* showed an exome-wide significant rare-variant burden in previous single-gene analysis^26,33^, and even this association could not be replicated in multiple cohorts. The network statistically supported by NERINE showed trait-increasing effects of rare damaging missenses in several *SNCA*-interactors in the stem, including *VDAC1*, *NEDD4*, and *TOR1A*, which were previously linked to PD pathology only in experimental models^61,62,77–81^. *PRL* also showed a strong trait-increasing effect, but it was not previously causally linked to PD. Beyond direct interactors of *SNCA* in the module, NERINE detected non-zero effects in genes such as *NDFIP1*, *PBX1*, and *RNF11* with PD risk, which had been linked to PD pathobiology only in functional studies^82–85^ but not through rare variant studies.

### Convergence of an unbiased functional genomics screen in CRISPRi-induced synucleinopathy model with NERINE on *PRL-SNCA* interaction

NERINE pinpointed genetic signals within the humanized TransposeNet αS proteinopathy network, which was originally derived from yeast-based screens^2^ and therefore lacked the specific human cellular context for key network interactions. Brain-related disease like PD involves a plethora of cell types and peripheral systems that can influence brain function, offering many possibilities for follow-up experimentation. To focus on neuronal mechanisms, we sought convergence between NERINE’s findings and an independent genome-scale functional screen in a CRISPR interference (CRISPRi) induced synucleinopathy cortical neuron (CiS-CN) model^28^ designed for studying genetic determinants of αS toxicity (**Methods**). Briefly, this model was established in the WTC11 iPSC genetic background^28^, and enabled—i) one-step trans-differentiation (AAVS1-Ngn2) into cortical glutamatergic neurons that are vulnerable in Lewy body diseases, including PD; ii) flexible gene knockdown via CRISPRi (dCas9-KRAB)^86^; iii) lentivirally-driven clonal αS expression at brain-like levels. The CiS-CN model, with two *SNCA*-overexpression (*SNCA*-OE) clones (*SNCA-*high and *SNCA-*intermediate) and a control clone (*SNCA-*endo1), thus provided a tractable but physiologically relevant system for studying αS aggregation and toxicity in cortical neurons^28,87^.

CRISPRi was utilized to target physical and genetic interactors of ɑS in the CiS-CN model, including all genes from the TransposeNet ɑS proteinopathy network, using a custom sgRNA library (**Methods**). The full screen will be described in a forthcoming publication. Briefly, we assessed sgRNA representation at start, DIV28, and DIV42, and looked for the dropout of sgRNAs in three comparisons: (1) *SNCA*-high vs. *SNCA*-endo1 at DIV42, (2) DIV42 vs. DIV28 in *SNCA*-endo1, and (3) DIV42 vs. DIV28 in *SNCA*-high (**Figures 7A,B**). Genes showing significant sgRNA dropout in MAGeCK-iNC analysis (FDR < 0.1) were classified as αS toxicity enhancers (**Methods**).

**Figure 7.**
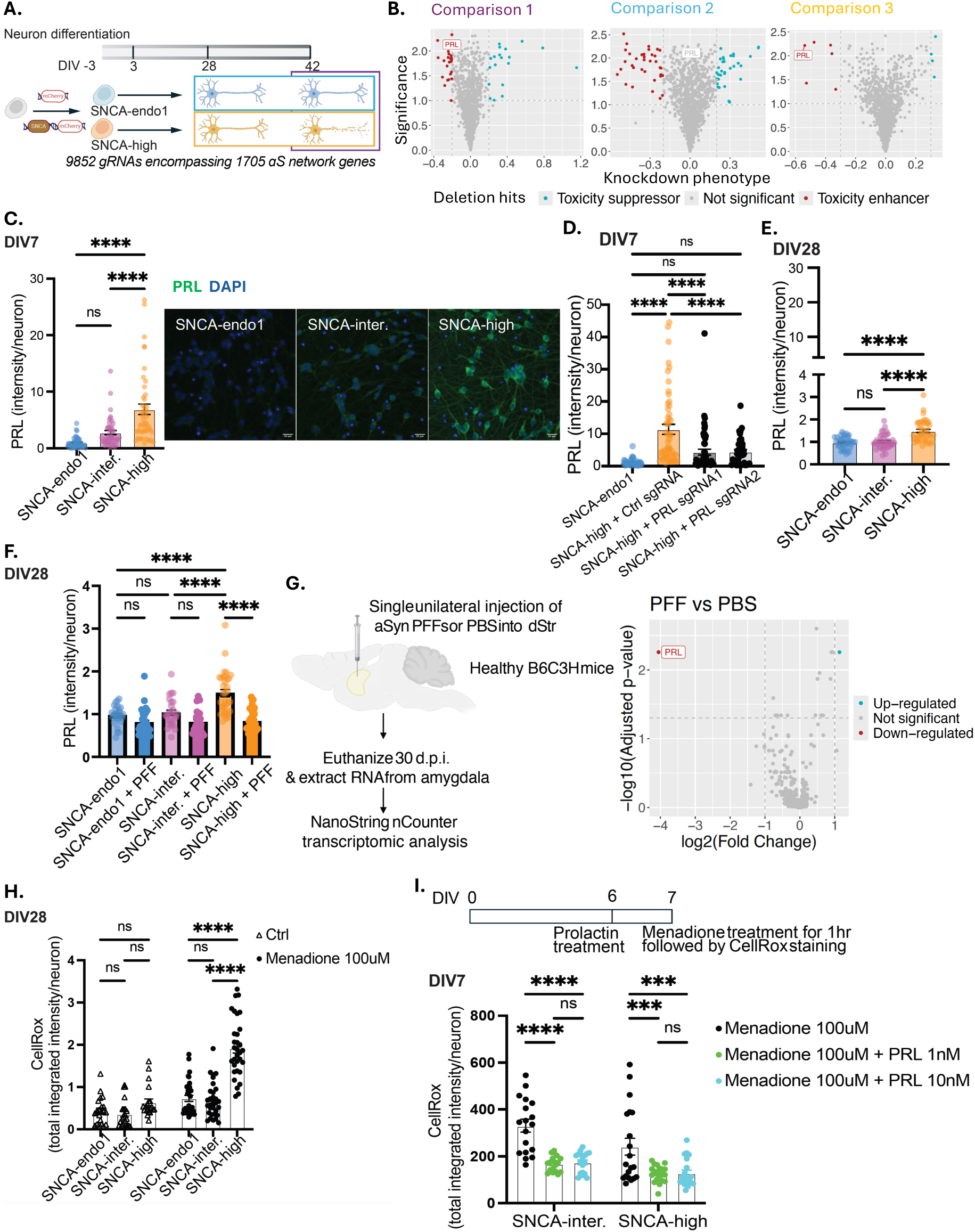
Unbiased functional genomics screens converge on an intraneuronal *SNCA-PRL* stress response. **A.** Top: Timeline of CiS-CN differentiation. Bottom: iPSCs were transduced to overexpress either mCherry (control) or SNCA-mCherry, generating the *SNCA-endo1* and *SNCA*-high lines, respectively. At DIV0, neurons were transduced with the sgRNA library. Neuronal samples were harvested and sequenced at DIV3, DIV28, and DIV42 using next-generation sequencing. *Comparison 1* shows the comparison of the sgRNA frequencies between DIV42 *SNCA*-high neurons and DIV42 *SNCA-*endo1 neurons. *Comparison 2* compares the sgRNA frequencies in *SNCA-*endo1 neurons between DIV42 and DIV28. *Comparison 3* compares the sgRNA frequencies in *SNCA*-high neurons between DIV42 and DIV28. **B.** Left: *PRL* sgRNA-containing neurons dropped out in Comparison 1, indicating PRL knockdown was toxic to *SNCA*-high neurons compared to *SNCA-*endo1 neurons. Middle: No significant drop-out in *PRL* sgRNA-containing neurons was observed in Comparison 2, indicating PRL knockdown was non-toxic to DIV42 *SNCA*-endo1 neurons compared to DIV28 *SNCA-*endo1 neurons. Right: *PRL* sgRNA-containing neurons dropped out in Comparison 3, indicating *PRL* knockdown was toxic to DIV42 *SNCA*-high neurons compared to DIV28 *SNCA*-high neurons. **C.** Left: Immunostaining data show prolactin was upregulated in DIV7 *SNCA*-high neurons (n=3, one-way ANOVA). Right: Representative images acquired with a Nikon Eclipse Ti microscope, with solar power set to 10% and an exposure time of 50ms for *prolactin*. Scale bar: 20um. **D.** *PRL* knockdown was validated with two different *PRL* sgRNAs from the screen library (n=3, one-way ANOVA). DIV0 *SNCA*-high neurons were transduced with either control sgRNA or PRL sgRNA at MOI=5. At DIV7, neurons were stained with *prolactin* antibody and Hoechst. Images were captured with a Nikon Eclipse Ti microscope. *Prolactin* intensity per neuron is reported here. **E.** Immunostaining data show *prolactin*-upregulation was diminished in DIV28 *SNCA-*high neurons (n=3, one-way ANOVA). Images were captured with a Nikon Eclipse Ti microscope, with solar power set to 30% and a 30ms exposure time for *prolactin*. **F.** Immunostaining data show *prolactin*-upregulation was diminished in DIV28 *SNCA*-high neurons treated with PFF (n=3, one-way ANOVA). Images were captured with a Nikon Eclipse Ti microscope, with solar power set to 30% and a 30ms exposure time for *prolactin*. **G.** Left: Illustration of the PFF-induced mouse model. Amygdala tissue, micro-dissected from mice and injected with *αS* PFFs, was subjected to transcriptomic analysis using the NanoString Neuropathology panel (**Methods**). Right: In comparison to PBS-injected control animals, which had no pS129 positive inclusions across all brain regions, PFF-injected mice showed a 16.6-fold downregulation of *Prl* mRNA expression in the amygdala region ipsilateral to the site of injection (n=5 per group; 3 males/2 females). **H.** CellRox assay shows that, with menadione treatment, oxidative stress in DIV28 *SNCA*-high neurons was significantly higher than in others of the same age (n=3, two-way ANOVA). **I.** Top: Timeline of exogenous *prolactin* assay. Bottom: CellRox assay shows that exogenous *prolactin* significantly decreased menadione-triggered oxidative stress in DIV7 *SNCA*-OE neurons (n=3, two-way ANOVA).

In these comparisons, *PRL* emerged as a particularly intriguing finding. NERINE had already pinpointed *PRL* within the LRRK2– and SNCA-containing branch of the same TransposeNet network (see above; **Figure 6D**), where damaging missense variants showed a strong trait-increasing effect. In the CiS-CN CRISPRi screen, sgRNAs targeting *PRL* significantly dropped out in *SNCA*-high versus *SNCA*-endo at DIV42 (Comparison 1) and from DIV28 to DIV42 in *SNCA*-high clones (Comparison 3), aligning with the NERINE result. Thus, in light of these convergent results, *PRL*, which previously lacked direct evidence of a causal role in PD, was selected as a key candidate for further experimental validation.

### A novel intraneuronal *SNCA-PRL* stress response identified in human neurons

While *Prl* is predominantly expressed in neuroendocrine tissue, its unexpected identification in our neuronal functional genomics screen suggests a highly unexpected cell autonomous role in neurodegeneration. Previous studies, though conflicting, have suggested *PRL* expression in rodent neurons outside the pituitary under stress conditions^88^, with speculation, albeit based on a few studies, that its expression may only be detected under conditions of neuronal stress. There is also evidence that its protein levels may be post-translationally regulated despite low mRNA expression^89^. Overall, the strong literature consensus is that any effects of *Prl* are likely to be through exogenous release of *Prl* from the pituitary gland.

To investigate the possibility that *Prl* may be part of an intraneuronal stress response directly related to αS, we performed immunostaining in neurons of different age. *Prl* expression was significantly increased in *SNCA-overexpression* neurons, with ∼7-fold-higher expression in *SNCA-high*, and 2.5-fold-higher in *SNCA-intermediate.* than in *SNCA-*endo1 neurons at DIV7 (**Figure 7C**). This elevation was abrogated, as expected, by our sgRNAs directed to *PRL* (**Figure 7D**). Importantly, *Prl* levels were reduced over time, such that by DIV28, *Prl* levels in the *SNCA-*high lines, were only ∼1.5-fold-higher than in *SNCA-*endo1 neurons (**Figure 7E**). These data collectively suggest a potentially protective role of αS-induced *Prl* that diminishes over time.

We next tested whether inducing αS aggregation, thereby reducing soluble αS levels^87^, in cellular and animal models via pre-formed αS fibrils (PFFs)^29,90^ led to reduced *Prl* levels. Our CiS-CN neurons challenged with seven (7) days of exposure to PFFs showed a reduction of *Prl* levels as indicated by immunofluorescence (**Figure 7F**). We further considered whether this was transcriptional or post-transcriptional. mRNA expression levels of *PRL* were low in our models, possibly related to the immaturity of the iPSC-derived neurons. We thus turned to a well-established mouse PFF model^29^. We unilaterally injected PBS versus 5 µg of PFFs (2.5 µL volume) into the dorsal striatum of mice (**Figure 7G: left**, **Methods**). Mice were aged for 30 days post-injection. By this stage, as has been previously described^91^, the αS aggregation pathology “spreads” distally and reaches the cortex and the amygdala, two brain regions highly susceptible to αS pathologies in later stage PD. We harvested mRNA from the amygdala and assessed gene expression with the Nanostring Neuropathology panel. In comparison to PBS-injected control animals which showed no pS129 positive inclusions across all brain regions (not shown here), PFF-injected mice showed a 16.6-fold downregulation of *Prl* mRNA expression in the amygdala region ipsilateral to the site of injection (n=5 per group; 3 males/2 females; **Figure 7G: right**), consistent and indeed stronger than our findings in the shorter-term human CiS neuronal model. At least in the intact rodent model, our data indicate that a component of this effect is transcriptional.

Our data indicate that in the context of aging and fibrillar αS pathologies, *Prl* levels drop intraneuronally, and potentially could sensitize the neurons to αS-induced cytotoxicity. To determine whether *Prl* directly protects against αS toxicity, we assessed its neuroprotective effects against oxidative stress in our CiS models. By DIV28, *SNCA-*high neurons exhibited heightened sensitivity to menadione-induced oxidative stress^92^, measured with the fluorogenic probe CellRox (Thermo) (**Figure 7H**). In *SNCA-*high and *SNCA-*endo1 models, we pre-conditioned DIV6 CiS neurons with *prolactin* for 24hrs before treatment with menadione (**Figure 7I: top**). Importantly, exogenous *prolactin* treatment significantly decreased oxidative stress in CiS neurons (**Figure 7I: bottom**), strongly suggesting a neuroprotective effect against αS-induced toxicity. Our data collectively indicate that aged neurons in which αS aggregates leads to reduced Prl levels and increased vulnerability to oxidative stress. Taken together, the convergence of our forward genetics iPSC screen with NERINE uncovered an unexpected αS-*Prl* intraneuronal stress response, suggesting that *Prl* may play a crucial role in neuronal resilience to αS pathology in PD. These data underscore the utility of NERINE in uncovering novel biological insights into complex disease processes.

## Discussion

NERINE is a powerful statistical test designed to bridge the gap between human genetics and experimental biology, adding a crucial mechanistic understanding to rare variant burden analysis. Unlike traditional rare variant tests, which focus on isolated genes or regions, NERINE contextualizes rare variant effects within biological networks, thereby enhancing our ability to interpret genetic associations. By incorporating network topologies, we provide a more structured framework for understanding how rare variants contribute to disease mechanisms.

One of NERINE’s key advantages is its ability to select between competing network topologies for the same gene set—an ability no other method currently offers. By utilizing edge geometry, NERINE gains a statistical power boost, a feature demonstrated through extensive simulations and real-world data analyses, even in cases where network structures are poorly defined. Despite its strengths, NERINE has some limitations, particularly in cases when the networks are too large, or sample sizes are exceedingly small (see **Methods**). However, its ability to integrate human genetic data with functional insights from model system experiments makes it a valuable tool for hypothesis testing in disease genetics.

To showcase NERINE’s utility, we applied it in two distinct contexts: first, to test network hypotheses from publicly available databases across four prevalent diseases—BRCA, T2D, CAD, and MI. NERINE provided novel rare variant signals for biologically plausible disease mechanisms, such as collagens and inflammatory response genes in cardiovascular diseases, adipogenesis pathway in T2D, and the network of estrogen receptor regulation in BRCA. The implication of collagens, key structural proteins in the extracellular matrix, in MI is consistent not only with their primary role in maintaining the stability of atherosclerotic plaques^93^ but also with their indirect role in coagulation via platelet effects and interactions with clotting factors^94^. Furthermore, the significant rare variant burden in networks linking inflammatory response genes to the risk of both CAD and MI supports the potential of anti-inflammatory therapies as an alternative to LDL-lowering drugs for treating cardiovascular diseases^95^. Second, we applied NERINE to bespoke gene networks derived from experimental screens in Parkinson’s disease (PD), a complex disorder where traditional rare-variant analyses have struggled due to limited sample sizes. Here, NERINE revealed a significant rare variant burden in gene modules related to autophagy regulation, vesicle trafficking, and protein homeostasis—signals largely undetected by single-gene tests.

Our findings further highlighted two major contributions: first, we provided rare variant evidence supporting hits from previous functional studies in model systems. For example, several member genes of the *LRRK2-and SNCA*-containing *vesicle tra[icking and protein homeostasis* subnetwork (e.g., *NEDD4*, *NDFIP1*, *VDAC1, PBX1, TOR1A*, and *RNF11*) which have been previously implicated in PD only in functional studies^61,62,77–85,96–98^, are now corroborated with human genetics evidence by NERINE. Second, NERINE helped resolve conflicting hypotheses by providing genetic support for specific disease mechanisms. For instance, there is biological data suggesting haploinsufficiency of *DYRK1A*—a member of the PD GWAS gene module involved in *peptidyl-threonine modification* identified by NERINE—reduces its kinase activity and exacerbates DA neuron degeneration in mice^99^. There is also data arguing the opposite through phosphorylation of ɑS^100^. By providing genetic evidence in favor of the latter direction, we offered some resolution of the matter.

Although NERINE does not provide gene-level p-values within a network, it yields informative effect sizes and directions for member genes that often align with known biology, as demonstrated in our positive control experiments with lipid phenotypes in the UK Biobank. Importantly, NERINE captures both trait-increasing and trait-decreasing rare variant burdens at the gene level, derived from its joint modeling of network interactions and observed allele counts at each locus. While the latter may reflect gain-of-function effects requiring experimental validation, they can be of considerable therapeutic significance. A well-established precedent is *PCSK9*, where rare LoF variants are associated with reduced LDL cholesterol levels; this discovery directly enabled the development of PCSK9 inhibitors as an effective therapeutic strategy for hypercholesterolemia with significantly improved cardiac outcomes^44,45^. In this context, the trait-decreasing effects predicted by NERINE within disease-relevant networks may highlight genes of potential therapeutic interest, supporting future experimentation.

NERINE’s integrative framework for unifying insights from model system screens with human genetics complements traditional pathway-based investigations in PD^70,101^. Traditional investigations rely on pathway-based polygenic risk scores from common variants or on rare variant aggregation tests, such as SKAT-O, applied at the pathway level^70,101^ to implicate broad mechanisms, including endolysosomal trafficking, GPCR signaling, neuronal transmission, and immune response^70,101^. While informative, such approaches generally lack statistical power, and resolution to pinpoint contributions of individual genes and their specific interactions. A network-based approach like NERINE, which incorporates experimentally defined gene relationships, provides additional biological specificity, improves power to detect associations, and helps guide mechanistic hypothesis generation for follow-up studies.

Among NERINE’s findings in PD, the most intriguing one is related to *PRL*, supported by converging evidence from NERINE and our functional CRISPRi cellular screens. *PRL* encodes the protein *Prl*, a hormone secreted by the pituitary gland with no known connection to ɑS. The original TransposeNet network^2^ unexpectedly introduced *Prl* to a small network of genes that also included key known PD genes SNCA (encoding ɑS itself) and *LRRK2* (encoding a protein with increased enzymatic kinase activity that has been implicated in PD). Mechanistically, a connection of *Prl* to ɑS had been elusive until this study. This is because, although not unprecedented, the presence of *Prl* in neurons outside the pituitary has been highly speculative^88,102^. While *Prl* showed some neuroprotective effects against oxidative stress^88,103–105^, this effect had been attributed exclusively to an exogenous endocrine effect through pituitary secretion. It was thus remarkable for *Prl* loss-of-function to simultaneously be implicated by both NERINE from human genetic data and in a genome-scale screen for enhancers of ɑS proteotoxicity: predicted deleterious missense mutations were tied to PD risk by NERINE, and *Prl* knockdown enhanced ɑS toxicity within neurons. Our data suggest that the effect results from an early, possibly post-translational, intraneuronal mechanism in which *Prl* is specifically elevated in response to ɑS overexpression. This response diminishes with time and with ɑS aggregation, the latter effect being at least partly transcriptional in more chronic treatment of mice with fibrillar ɑS. This decrease is, in turn, associated with the sensitization to *PRL* knockdown and exogenous stress, demonstrated in this study with the oxidative stressor menadione. While we could not directly assess pituitary function in individuals carrying *PRL* variants in our study cohorts, literature evidence provides additional context. A recent proteomic study in the Harvard Biomarkers Study cohort found that *Prl* levels in the cerebrospinal fluid—likely reflecting non-pituitary sources—was the top feature in a classifier model distinguishing PD patients from controls, even after accounting for levodopa effects^106^. Thus, these converging findings from NERINE, our CRISPRi screen, mouse models, and literature strengthen the plausibility of a role for an unexpected role of *Prl* in PD that originates not in the pituitary but within neurons subjected to ɑS stress. Further work will now be necessary to clarify its mechanisms and to evaluate potential pleiotropic effects.

In sum, NERINE bridges the gap between human genetics and experimental biology, providing a robust framework for attributing putative disease-causing factors to experimentally derived molecular networks. Such a method is especially significant for complex diseases like PD, where the limited availability of well-phenotyped genomic data hinders traditional approaches. As large-scale molecular interaction networks become increasingly common, we anticipate that research programs where human genetics and experimental biology mutually reinforce each other—such as those enabled by NERINE—will play a growing role in identifying disease-relevant signals, resolving conflicting evidence, and uncovering druggable targets.

## Methods

### Overview of the NERINE methodology

NERINE models the “variant-gene-network” hierarchy as follows: it encodes gene-gene relationships in a network of *m* genes by a positive semidefinite matrix, Σ. We assume that phenotypic effects of genes within a network, represented as vector 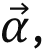 are drawn from a multivariate skew-normal distribution 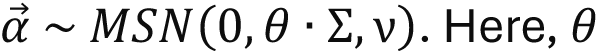 Here, *θ* is a parameter reflecting the cumulative effect of the gene network on a phenotype and is the object of inference (**Figure 1**). The skewness parameter *υ* = *f*(*N_case_*, *N_control_*) adjusts for case-control imbalance. Here, N_case_ and N_control_ represent the case-and control-group sizes. For balanced cohorts (i.e., *ν* = 0), the model reduces to 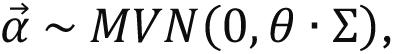 where marginals are normally distributed with zero mean. Under this model, an edge between two genes in the network implies that they have either correlated non-zero effect sizes or correlated chances of having no phenotypic effects.

The network-effect, *θ* on a dichotomous phenotype is estimated using the maximum likelihood estimation (MLE) framework, where the likelihood is modeled as an integral over two components: (i) the product of conditional probabilities of observed case mutation counts given total mutation counts in network-genes, and (ii) the probability of gene effects given the network parameters:

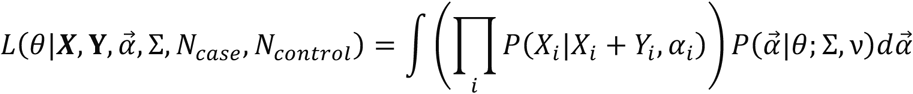

We approximate this integral as a weighted sum over multivariate quadrature points from the domain of integration. The weight of each multivariate quadrature point is given by the product of the corresponding univariate weights. To make the computation of the integrand tractable, NERINE makes several design choices and employs a lookup table approach with pruning (**Supplementary Note**).

NERINE models rare variant counts in genes for cases and controls as two independent Poisson random variables as follows,

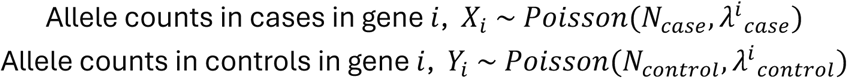

Here, *λ_case_^i^* and *λ_control_^i^* denote the rate parameters for case-and control-allele count distributions in gene *i*, and correspond to population allele frequencies renormalized between cases and controls by transformed network-gene effect, *α_i_*. This implies that the conditional probability of observed rare variant count in each gene in cases, given the total rare variant count in the cohort, follows a Binomial distribution (**Supplementary Note**).

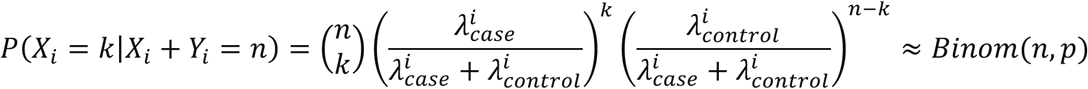

where *p* = *Ф*(*α_i_*) (**Supplementary Note**).

Since each *α*_i_represents unbounded univariate skew normal distributions, they cannot be directly used as Binomial success probabilities. Hence, we apply a custom variable transformation to map *α_i_*s to Beta distributions so that they fall on the interval [0, 1] (**Supplementary Note**). This transformation ensures that the mean gene effect is centered around *N_case_*/(*N_case_* + *N_control_*), and the shape of the transformed distribution adapts with *θ*: small *θ*s (close to 0) concentrate density near the mean, while large *θ*s shift density toward the extremes. Case-control skew is incorporated using the shape parameters of the Beta distribution. The transformed gene-effects within the network are denoted 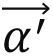 (**Supplementary Note**).

Thus, NERINE’s approximate likelihood takes the form:

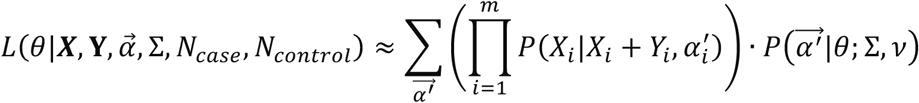

The conditional probability term is calculated using the probability density function of a standard Binomial distribution, which makes the likelihood computation fast and tractable. To calculate the probability of network-gene effects 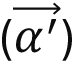 for a given network topology (Σ) and network effect (*θ*), we use a lookup table, where highly improbable 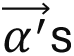 are pruned (**Supplementary Note**).

NERINE performs nested hypothesis testing; the null hypothesis being *H*_0_: *θ*=0 (i.e., the gene network does not affect the trait and expected ratio of variant counts between cases and controls is proportional to the ratio of sample sizes in each gene) and the alternative set of hypotheses being *H*_z_: *θ* = *θ*_z_>0 (i.e., the gene network has an overall effect on the trait and the ratio of variant counts between cases and controls may deviate from the null expectation in relevant member genes). The test statistic of NERINE is the log-likelihood ratio (LLR):

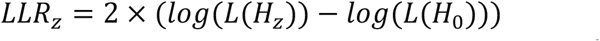

We denote the maximum-likelihood estimate of network effect with 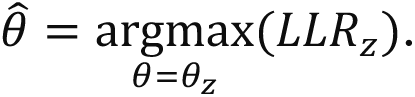 Since *θ* lies on the boundary of the parameter space, the test statistic asymptotically follows the distribution of a weighted mixture of a point mass at zero and a chi-square distribution with a degree of freedom of one (*dof* = 1)^30,31^. NERINE draws its asymptotic p-values from this distribution. For significant networks, NERINE calculates the maximum likelihood gene-specific effects 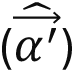 under the estimated 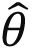 as follows:

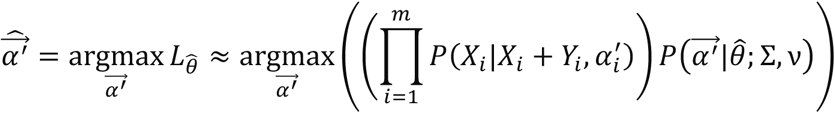

The search space for possible network-gene effects 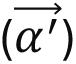 is determined by the estimated 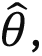 the network structure (Σ), and the lookup table entries (**Supplementary Note**).

NERINE has several limitations. Currently the test is designed to accommodate dichotomous traits only. Continuous traits need to be dichotomized before NERINE can be run on them. Covariates correction is not directly included in the test. For continuous traits, it can be done as a preprocessing step using a regression-based approach and then the trait can be dichotomized. For binary traits, NERINE currently performs ancestry-stratified analysis and combines the p-values using Fisher’s combined test post-hoc. For large networks with > 50 genes, NERINE’s test statistic falls outside the asymptotic regime, hence we recommend using NERINE with networks up to 50 genes.

### Simulations under the null model

To evaluate the performance of NERINE under the null model (*θ*=0), we performed extensive simulations with well-studied biological pathways, namely, NOTCH signaling (m=6), WNT signaling (m=24), protein export (m=24), and EGFR signaling (m=50), from the canonical pathways database (**Supplementary Figures S1 and S2**) and different simulated network architectures—(i) clique: complete graph with all nodes connected to each other; (ii) path: each node connected to two other nodes except the first and the last nodes; (iii) random: randomly generated scale-free graph of *m* nodes; and (iv) isolated genes: nodes not connected with each other (**Supplementary Figure S3**).

For the canonical pathways, gene lists were extracted from MSigDB (v7.3) and high confidence physical and genetic interactions from protein-protein interaction (PPI) databases were used as network edges between pathway genes. We simulated three different scenarios – (i) equal sized case/control groups (*N_case_* = 1,000; *N_control_* = 1,000) (ii) case group is larger (*N_case_* = 3,000; *N_control_* = 1,000), and (iii) control group is larger (*N_case_* = 1,000; *N_control_* = 3,000).

Under different network architectures and case-control skews, we simulated allele counts in cases and controls using independent Binomial distributions under the null model. For each gene, we assumed the presence of up to five qualifying loci each with minor allele frequency (MAF) of 0.001. The binomial probabilities for case-and control-groups are adjusted according to the group sizes and MAF of variants assuming no gene-specific effects under the null model (*θ* = 0). For each scenario, we performed 1,000 iterations to generate the QQ-plots.

We used the *pchibarsq* function from the *emdbook* (v1.3.13) in R (v4.3.2) to calculate the p-values from the mixture of chi-square distribution with one degree of freedom (*dof* = 1) and the delta function at zero (0). We calculated 95% bootstrap confidence intervals around NERINE’s test-statistic for visualization.

For evaluating the null behavior of NERINE’s test statistic in simulated networks of different sizes (*m* = 5, 10, and 25 genes) and different topological architectures (i.e., clique, path, random, and isolated nodes), we used equal sized case-and control-groups (**Supplementary Figure S3**). For each scenario, we performed 1,000 iterations to generate the QQ-plots. The allele counts in cases and controls were generated from independent Binomial distributions following the same procedure as above.

### Performance benchmarking with simulated data

We evaluated the performance of NERINE under the alternative hypothesis (*θ*>0) in two sets of simulations—(i) when genes have only trait-increasing effects, and (ii) when genes have both trait-increasing and trait-decreasing effects using the same database pathway network topologies used for the null simulations (**Figure 2**, **Supplementary Figure S4**). For each scenario, we simulated different noise profiles, i.e., varying proportions of genes within the network with effects on the trait given the network topology. This mimics situations from having a very noisy network (i.e., ∼10-30% genes with an effect on the trait) to a highly relevant network (i.e., ∼70-90% genes with an effect on the trait). We simulated allele counts from cases and controls using independent Binomial distributions under the alternative model with *θ* = 0.2. For each gene, we assumed the presence of up to five (5) qualifying loci per gene with minor allele frequency (MAF) of 0.001. The Binomial probabilities for case-and control-groups are adjusted according to the group sizes and MAF of variants assuming possible gene-specific effect configurations under the alternative model (i.e., *α⃗* given the network topology (Σ) and network effect, *θ* = 0.2). Using this setup, we simulated a cohort of 2,000 cases and 2,000 controls.

Currently, there are no existing rare variant association tests that take gene network topology into account Thus, we compared the performance of NERINE with existing gene-level rare variant association tests adapted to the pathway level, namely, CMC-Fisher test^107^, Fisher minimum p-value test^108^, Fisher combined test^108^, SKAT-O^18^, and pathway-based rare variant trend test (RVTT)^15,16^. For each noise profile, we performed 250 iterations, resulting in 1,000 iterations per network. Empirical power of each method was measured as the positive predictive value (PPV) across iterations using different p-value cutoffs (*c*): 1e-2, 5e-3, 1e-3, 5e-4, 1e-4, 5e-5, and 1e-5. Here,

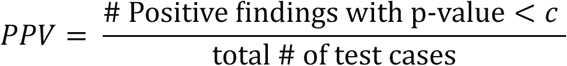

Additionally, we conducted performance benchmarking for NERINE on different simulated network architectures for 25 genes (**Supplementary Figure S5**). As before, we performed two sets of simulations—(i) when genes have only trait-increasing effects, and (ii) when genes have both trait-increasing and trait-decreasing effects for four network topologies—clique, path, random graph, and isolated nodes. We simulated allele counts in 1,000 cases and 1,000 controls using independent binomial distributions under the alternative model with a different network effect, *θ* = 0.5. We simulated various network-noise profiles ranging from 0-90% following the same procedure as above. Empirical power of each method was calculated as PPV across 250 iterations per noise profile per network topology, resulting in 1,250 iterations per network. Since RVTT, by design, assumes that all qualifying rare variants in a pathway have the same direction of effects, we compared RVTT’s performance with NERINE in simulations with genes having only trait-increasing effects. Moreover, RVTT computes a permutation-based p-value, and for 10,000 iterations its p-values cannot be less than 1e-4. Hence, we compared RVTT’s performance only at cutoff values ≥ 1e-4.

Furthermore, we evaluated NERINE’s performance with true versus randomized network architectures in simulations with the same four well-studied pathways (**Supplementary Figure S10**). We constructed the “true” network topologies for these pathways using the high-confidence protein–protein and genetic interactions from PPI databases as described above. For each gene set, we also generated 100 randomized networks by randomly introducing edges among the member genes. A binary trait was simulated in 2,000 cases and 2,000 controls with varying trait-increasing and trait-decreasing effects under the alternative model with a network effect, *θ* = 0.1. We simulated different noise profiles (i.e., 10-30%, 30-50%, 50-70%, and 70-90%) and performed 250 iterations per noise profile. We measured NERINE’s PPV at different p-value thresholds (i.e., 1e-2, 5e-3, 1e-3, 5e-4, 1e-4, 5e-5, 1e-5).

### Pathway database construction

We created a pathway database with all canonical pathways of five to fifty genes from the BIOCARTA database along with all lipid-, DNA replication-, DNA damage repair-, and cell cycle-related pathways from the REACTOME, KEGG, PID, and Wiki pathways databases. The lists of member genes for these pathways were extracted from the Molecular Signatures Database (MSigDB v7.3; https://www.gsea-msigdb.org/gsea/msigdb). The database contained 306 pathways with a median pathway length of 25 genes (**Supplementary Table T20**). Due to the pleiotropy of genes, many biological pathways tend to overlap significantly with each other. Thus, we determined the effective number of independent hypotheses (*t_eff_*) in our pathway database of *t* pathways by adapting Nyholt’s approach^109^.

First, we calculated the pairwise Jaccard similarity of pathways, and encode it in a matrix, *M*_t×t_, which is symmetric but not positive semi-definite. We then converted it to its nearest positive definite matrix using the *nearPD* function^110^ from the *Matrix* package (v1.6-5) in R (v4.3.2). Next, we determined the eigenvalues 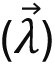 of this approximate covariance matrix. The effective number of hypotheses/pathways was computed using the following formula^109^:

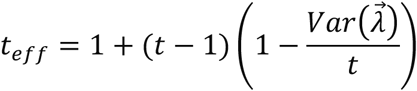

For our pathway database, we found the effective number of independent hypotheses, *t_eff_* to be 300, which was used for Bonferroni correction to determine database-wide significance.

### Gene-network topology extraction

NERINE treats the gene-gene network as an input and can flexibly handle any symmetric pairwise relationship matrix that is positive-semidefinite. For a screen that provided the gene network topology, we used that network as is. For example, for the *α*-synuclein proteotoxicity screen in PD, we directly applied NERINE on the published TransposeNet humanized *α*-synuclein-modifier network^2^ stems. In absence of the true network topology for a particular gene set, we adopted the following approach to construct one.

For canonical database pathway gene sets, we constructed network topologies by extracting physical and genetic interactions from several sources: (i) high confidence physical interactions (weight >= 0.7) from STRING v11.5^111^, (ii) InWeb inBio Map database^112^, (iii) HuRI^6^, (iv) genetic interactions from the Megchelenbrink *et al.* study^113^, and (v) humanized *α*-synuclein-, *β*-amyloid-, and TDP-43-modifier networks^2^. Using a heuristic approach, we first generated adjacency matrices for the genes, where edges were represented as binary values (1 = presence of an edge; 0 = absence of an edge), corresponding to the non-zero oi-diagonal entries of the network. The diagonal was set to two (2) to indicate an equal prior on each gene. Because these modified adjacency matrices were not always positive semidefinite, we applied the *nearPD* function^110^ from the **Matrix** package (v1.6-5) in R (v4.3.2) to compute the nearest positive-definite matrix. This matrix was input to NERINE as an approximate covariance matrix, Σ.

We also constructed co-expression networks in relevant tissue types for different phenotypes using the bulk tissue expression data from the Genotype Tissue Expression database^114^ (GTEx v8). We computed a real-valued gene-gene co-expression networks in a specific tissue where the edges between two genes represented the Pearson correlation of their expression profiles in that tissue types. For lipid-related phenotypes, we constructed co-expression networks using the bulk expression data from liver tissue. For Parkinson’s disease (PD), we used bulk expression data from the substantia nigra region of the mid-brain.

To construct co-essentiality networks in relevant cell lines for different phenotypes, we used the gene dependency data from CRISPR knockout screens from project Achilles, as well as genomic characterization data from the Cancer Cell Lines Encyclopedia (CCLE) project from the DepMap portal^115,116^ (release: 2023Q2). We computed a gene-gene co-essentiality networks in relevant cell lines where the edges between two genes represent the Pearson correlation of their dependency profiles in those cell lines. While for lipid-related phenotypes, we constructed co-essentiality networks using data from liver cell lines, for PD, we used data from cell lines pertaining to the central nervous system (CNS). The cell lines used in this study are listed in the **Supplementary Table T21**.

### Performance benchmark on lipid phenotypes in the UK biobank (UKBB)

Using jointly genotyped variant calls from the whole exomes of 469,589 individuals in the UK biobank (UKBB), we performed positive control experiments with NERINE in two lipid related phenotypes: direct LDL cholesterol (LDL-C; data field: 30780) and HDL cholesterol (HDL-C; data field: 30760) (**Supplementary Figures S6-S9**). We created two dichotomous phenotypes: (i) high LDL vs. low LDL and (ii) low HDL vs. high HDL, by selecting individuals belonging to the top and bottom quartiles of the distributions for LDL-C and HDL-C measurements. European ancestry groups had the largest sample sizes: LDL-C (30,007 cases and 28,673 controls), and HDL-C (26,800 cases and 27,178 controls). NERINE was competitively applied on these two phenotypes across networks from our canonical pathways database. We extracted the edge relationships of genes from high-confidence physical and genetic interactions from protein interaction databases for this exercise. This analysis served as an ideal positive control because the genetic determinants of these phenotypes are well-annotated.

Variants with a minor allele frequency (MAF) < 0.001 were considered rare for the test to not be influenced by artificial signal from common variant space propagated through linkage disequilibrium (LD). We stratified variants into six functional categories: LoF (i.e., frameshifts, insertions, deletions, and splice variants), damaging missense (i.e., missenses predicted to be damaging by in-silico tools), damaging (i.e., LoF and damaging missenses), missense, neutral (i.e., missenses predicted to be benign by in-silico tools), and synonymous. We used neutral and synonymous variants in a pathway as control to safeguard against technical biases and LD leakage. Bonferroni correction, accommodating the presence of correlated hypotheses in the database^109^, was used to control for Type I error.

To ensure our results for LDL-C phenotype were not driven by *LDLR* and *PCSK9*, which have large individual effect sizes on LDL-C levels, we performed sensitivity analysis by removing *LDLR* and *PCSK9* from the significant networks for the LDL phenotype. Even without *LDLR* and *PCSK9*, the module of core LDL-related genes, along with the LDL clearance and chylomicron clearance pathways remained significant for the LDL phenotype after Bonferroni correction (**Supplementary Figure S22**).

For many disease phenotypes, the large sample sizes available for the UKBB lipid phenotypes, are simply not attainable. Thus, we performed a down-sampling experiment to evaluate the consistency of NERINE’s performance across different sample sizes and the robustness of NERINE’s performance with small sample sizes. We downsampled the high LDL vs. low LDL cohort at different case-control ratios (1/3, 1/10, and 1/60) and competitively applied NERINE across the pathway database, demonstrating NERINE’s effectiveness even in a cohort with as few as 500 cases and 500 controls (**Supplementary Figure S23**).

Since rare variants are more susceptible to subtle effects of population stratification than common variants, we performed stratified analysis of the LDL-C phenotype in all five major ancestry groups (i.e., European, American, African American, South Asian, and East Asian) and meta-analyzed the results using Fisher’s combined test (**Supplementary Figure S9**, **Supplementary Table T2**). Bonferroni correction was applied on the meta p-values.

We further evaluated NERINE on real versus randomized network topologies for binarized LDL-C and HDL-C phenotypes (**Supplementary Figure S11**). For each phenotype, we selected database-wide significant pathways with rare damaging variant burden and used the most significant topologies for each pathway shown in **Supplementary Table T3** as “real” network topologies. We also generated 100 randomized networks per pathway by introducing random binary edges among member genes. We applied NERINE to both real and randomized networks and assessed its performance by comparing LLRs and p-values; higher LLRs and lower p-values indicated superior performance.

### Study cohorts for common disease phenotypes

We used whole exome sequencing (WES) data from two population-scale biobanks, namely, the UK Biobank (UKBB) and the Mass General Brigham Biobank (MGBBB) and whole genome sequencing (WGS) data from the AMP-PD consortium for investigating network-level rare variant associations in complex diseases.

Our analysis focused on four representative, high-prevalence complex diseases of public health relevance: breast cancer (BRCA), type II diabetes mellitus (T2D), coronary artery disease (CAD), and early-onset myocardial infarction (MI). These phenotypes were chosen from the UK Biobank because large sample sizes were available, and replication cohorts existed in the MGB Biobank. They served to showcase NERINE’s applicability in comprehensive database searches for rare variant network burden.

Cohorts for BRCA, T2D, CAD, and MI were selected primarily based on the summary diagnoses recorded in the data field 41270 as ICD-10 codes. For BRCA, the case group consisted of unrelated females of European ancestry with ICD-10 code C50 and the control group consisted of unrelated European females of age 60 or above with no history of neoplasms (ICD-10 codes: C00-C97 and D00-D48). The resulting cohort had 10,648 cases and 91,886 controls. For T2D, we included unrelated European individuals with ICD-10 code E11 in the case group and unrelated European individuals with no endocrine, nutritional and metabolic diseases (ICD10 codes: E00-E90) in the control group. The resulting cohort had 22,502 cases and 68,370 controls. We created an age-stratified case-control cohort for the CAD phenotype where cases consisted of unrelated European individuals of age <=65 with ICD-10 code I25 and controls consisted of unrelated European individuals of age > 65 with no diseases of the circulatory system (ICD-10 codes: I00-I99). This left us with 4,561 cases and 12,321 controls. Finally, for the MI phenotype, our cohort consisted of 2,521 cases and 5,012 controls. Cases included unrelated European individuals with ICD-10 code I 21. Only males with age <=55 and females with age <=65 were included in the case group. Whereas controls consisted of unrelated European individuals of age >= 69 who have no history of any disease of the circulatory system (ICD-10 codes: I00-I99).

Parkinson’s disease (PD) was chosen as a case study to demonstrate NERINE’s ability to bridge human genetics with model-system screens. We had access to two genome-scale screens, targeting the main pathological features of the disease and providing experimental gene networks for testing. We also had our in-house iPSC-derived cortical neuron model (CiS-CN) for functional validation. We focused on sporadic disease cases as they constitute∼85% of the PD population.

For sporadic PD phenotype, we created a discovery cohort from UKBB individuals by including unrelated European individuals with ICD-10 code G20 and no family history of PD as cases. The control group consisted of unrelated European individuals with no history of PD and other nervous system diseases (ICD-10 codes: G00-G99). In the original UKBB cohort, we observed notable differences in the incidence of type 2 diabetes (T2D) between PD cases (17.5%) and controls (7.1%), which could bias downstream analyses; therefore, we excluded all individuals with a history of T2D from both groups. We termed this cohort the “UKBB-Sporadic” cohort with 2,237 cases and 167,188 controls. Controls were on average younger than cases at recruitment in this cohort (**Supplementary Table T22**), which could introduce bias when investigating signatures of dopamine (DA) neuron degeneration. Thus, for the DA neuron essentiality screen analysis, we employed a super-control design by selecting older controls with a higher median age at recruitment than the cases (**Supplementary Table T22**), which are more likely to capture naturally occurring signatures of neurodegeneration and less likely to have sporadic PD later in life. We termed this cohort with super controls as UKBB-Extreme (Ncase = 2,237 and Ncontrol = 2,553).

For the database-wide investigations of BRCA, T2D, CAD, and MI, we created replication cohorts using the jointly genotyped WES data from 53,343 individuals in MGBBB, a biorepository of consented patient samples at Mass General Brigham (parent organization of Massachusetts General Hospital and Brigham and Women’s Hospital). Same inclusion/exclusion criteria were used for each phenotype to ensure consistency between the biobanks. The resulting cohort sizes are as follows: BRCA (Ncase = 1,113; Ncontrol = 2,459), T2D (Ncase = 747; Ncontrol = 2,188), CAD (Ncase = 902; Ncontrol =1,488), and MI (Ncase = 326; Ncontrol = 2,068). For the BRCA cohort, we used an MAF cutoff of 0.001 to select rare variants. For the other three phenotypes, variants with MAF <0.03 were considered to be “rare”. Since the cohort sizes in MGBBB were smaller compared to UKBB for T2D, CAD, and MI, we did this adjustment to the MAF cutoff and performed tests in the synonymous and neutral missense categories for each pathway to make sure that there was no LD leakage.

For PD, we created a replication cohort using the WGS data from 10,418 individuals from AMP-PD (Accelerating Medicines Partnership: Parkinson’s Disease) v3 release (2022), which encompasses study participants at Mass General Brigham (i.e., Harvard Biomarkers Study 2.0). Any individual belonging to the genetic registry and genetic cohort group, as well as subjects without evidence of dopamine deficit (SWEDD) and subjects belonging to the prodromal categories and the AMP-LBD cohort were excluded from the analysis. As the AMP-PD cohort predominantly consists of individuals of European ancestry, we retained only unrelated individuals of the same ancestry group. We called the resulting cohort “AMP-PD-sporadic,” which consisted of 2,117 sporadic PD cases and 1,095 neurotypical controls. Both case and control groups had similar median ages in this cohort (**Supplementary Table 22**).

We performed both variant-and sample-level quality control (QC) steps on each dataset to ensure the study cohorts are free from technical biases as much as possible (see **Supplementary Note** for detailed steps). We retained only high-quality biallelic variants passing GATK best practices filters and having maximum 10% missingness for our analysis. We annotated variants with their functional consequences and gnomAD allele frequencies with VEP (v109) and dbNSFP (v4.3a) database. We used six masks to group variants into functional categories: (i) *Damaging missense*: missense variants predicted to be either ‘‘P’’ or ‘‘D’’ by PolyPhen2 (v2.2.3) or ‘‘deleterious’’ by SIFT (v6.2.1), (ii) *LoF*: variants labelled as splice donors, splice acceptors, splice region variants, stop-gained, stop-lost, start-lost, frameshifts, in-frame insertions, and in-frame deletions; (iii) *Damaging*: LoFs and damaging missenses, (iv) *Missense*, (v) *Neutral*: missense variants predicted to be either ‘‘B’’ by PolyPhen2 or ‘‘tolerated’’ by SIFT, and (vi) *Synonymous*. These masks were used in our analysis of binarized cholesterol phenotypes as well as complex diseases.

Notably, NERINE is agnostic to the variant annotation tool and can test any user-defined categories of rare variants. To assess robustness, we reclassified variants using REVEL^117^ (version May 2021) and AlphaMissense^118^ scores (hg38 v1) within the VEP (v109) plugin and custom annotation tools in a positive control experiment with the LDL-C phenotype in UKBB. We defined three categories: (i) *pathogenic missense* (REVEL > 0.649 or AlphaMissense > 0.564), (ii) *pathogenic* (pathogenic missense plus LoF), and (iii) *benign missense* (REVEL < 0.29 and AlphaMissense < 0.34). Results were highly concordant with those obtained using PolyPhen2 and SIFT (**Supplementary Figure S24**).

We removed sample outliers based on Ts/Tv, Het/Hom ratios, and per-haploid SNV counts, where outliers were defined as samples that were three standard deviations away from the mean. We retained only unrelated European samples for our analyses because, at the time, individuals of European ancestry constituted the largest group in our study cohorts. Concentrated efforts in building large biobanks with diverse participants are already underway^119^ and will enable NERINE to overcome this limitation and provide more insight into the contribution of rare variants to common disease etiology across populations.

### Human iPSCs culture and DA neuron di_erentiation

Human H9 (MSKCC stem cell core facility; WA-09) ESCs were cultured in Essential 8 (E8) medium (Gibco/Thermo Fisher Scientific; Cat. No. A1517001) on 10cm plates coated with Vitronectin (Thermo Fisher Scientific; Cat. No. A14700) diluted 1:100 in Dulbecco’s PBS (DPBS; Thermo Fisher Scientific; Cat. No. 14190136) and passaged at ∼80% confluence using StemPro Accutase Cell Dissociation Reagent (Gibco/Thermo Fisher Scientific; Cat. No. A11105-01). E8 medium was replaced every day. ESCs harboring a doxycycline-inducible Cas9 cassette in the AAVS1 safe harbor locus were differentiated following a previously published protocol^120^.

Briefly, ESCs were dissociated and plated at high density (600,000 cells/cm²) in Neurobasal medium (Gibco/Thermo Fisher Scientific; Cat. No. 21103-049) supplemented with 1X N2 supplement (Life Technologies; Cat. No. 17502048), 1X B27 supplement (Life Technologies; Cat. No. 17504044), 2 mM L-glutamine (Gibco/Thermo Fisher Scientific; Cat. No. 25030-081), penicillin-streptomycin (Gibco/Thermo Fisher Scientific; Cat. No. 15140122), 250 nM LDN193189 (Reprocell; Cat. No. 04-0074), 10 μM SB431542 (R&D Systems; Cat. No. 1614), 1 μM CHIR99021 (R&D Systems; Cat. No. 4432), 500 ng/mL SHH (R&D Systems; Cat. No. 464-SH), and 10 nM Y-27632 ROCK inhibitor (Bio-Techne; Cat. No. HY-10583). Cells were counted and plated on DMEM/F12 (Gibco/Thermo Fisher Scientific; Cat. No. 11320033) and Geltrex-coated plates (Life Technologies; Cat. No. A1413201) at high density (600,000 cells/cm^2^), while being maintained in this medium from DIV 0 (day of plating) to DIV 3. Y-27632 ROCK inhibitor was added only at plating. Medium change was performed on DIV 4 and every three days till DIV 9. On DIV 4, the medium was supplemented with 6µM CHIR99021. On DIV 7, the medium was composition was changed to exclude SB431542 and SHH. From DIV 10, cells were transitioned to maturation medium with the following composition: Neurobasal Medium as the base, 1X B27 Supplement, 2mM L-glutamine, penicillin-streptomycin, 3µM CHIR99021, 0.2mM Ascorbic Acid (Sigma-Aldrich; Cat. No. A4034), 20ng/mL GDNF (Gibco/Thermo Fisher Scientific; Cat. No. 450-10), 1ng/mL TGFβ3 (Gibco/Thermo Fisher Scientific; Cat. No. 100-36E), 20ng/mL BDNF (R&D Systems; Cat. No. 248-BDB), and 0.2mM dbcAMP (Sigma-Aldrich; Cat. No. 4043). For DIVs 10 and 11, cells were maintained in this medium.

On DIV 11, partially-differentiated cells were released and centrifuged as above, and pelleted cells were resuspended in the same medium from DIVs 10-11 and plated at high density (800,000 cells/cm^2^) on 15µg/mL Poly-L-Ornithine (Sigma-Aldrich; Cat. No. P3655), 2µg/mL Fibronectin (Gibco/Thermo Fisher Scientific; Cat. No. 356008), and 1µg/mL Laminin (R&D Systems; Cat. No. 3400-010-03)-coated plates in DPBS. From DIVs 12 to 15, cell were placed into maturation medium with a composition similar to DIVs 10-11 medium supplemented with 1 μM IWP-2 (Tocris Bioscience; Cat. No. 3533) and 100 ng/mL FGF-18 (PeproTech; Cat. No. 100-28). Doxycycline hydrochloride (2 μg/mL; Sigma-Aldrich; Cat. No. D3447) was added on DIVs 14 and 15 to induce Cas9 expression. On DIV 16, partially-differentiated cells were released and centrifuged as above, and pelleted cells were replated at 1,200,000 cells/cm² and resuspended in maturation medium with a composition similar to DIVs 10-11 medium supplemented with 10 μM DAPT (R&D Systems; Cat. No. 2634) for ten days. On DIV 25, neurons were dissociated and replated at low density (200,000 cells/cm²) and maintained in maturation medium containing 15µg/mL Poly-L-Ornithine, 2µg/mL Fibronectin, and 1µg/mL Laminin-coated plates in DPBS.

### Genome-wide CRISPR/Cas9 screen in PSC-derived DA neurons

We performed genome-wide CRISPR/Cas9 screen in DA neurons differentiated from human WA-09 (H9) embryonic stem cells as described above. The Gattinara human CRISPR pooled knockout library^121^ (Addgene; pooled library #136986) was used for this screen; this library includes two gRNAs for 19,993 genes as well as 500 non-targeting controls and 500 one-intergenic site targeting controls. Guide RNAs (gRNAs) representing 19,993 genes were transduced into H9 hESC lines at an MOI of 0.3-0.5 and 1000x representation. Transduced stem cells were selected by puromycin and differentiated toward DA neurons until reaching the neural progenitor stage as described by Kim and colleagues^122^. Briefly, at DIVs 14-16 of the differentiation, we induced iCas9 expression by doxycycline addition using the AAVS1 safe-harbor locus as previously described^123^, while cells were neural progenitors. Then, we waited until DIV25 when they differentiated into DA neurons to take our initial sample to obtain gRNA representation through whole genome sequencing (DIV26). The remaining neurons were allowed to stay in the dish until our final collection time (DIV42) when neuronal cell death began. Samples were processed for library preparation and sequenced and sequencing reads were aligned to the screened library and analyzed using *MAGeCK-MLE* from the *MAGeCKFlute*^124^ (v2.6.0) package in R (v4.3.2). Essentiality genes were classified as having Wald test FDR-adjusted p-value < 0.05 and beta <-0.58. Broadly essential genes^124^ that are not specific to DA neurons, were removed from the list of essentiality genes. We identified 693 essentiality genes. The complete screen will be described in a forthcoming publication.

### Constructing ontology-based network topologies from gene sets

For PD GWAS genes as well essential genes for DA neuron survival, we first performed gene set enrichment analysis using the *enrichr* function from the *GSEApy* package (v 1.1.3) in python (v 3.12.4) and identified all GO biological processes with nominal significance (p-value < 0.05). We then grouped semantically similar GO terms using REVIGO^125^ (http://revigo.irb.hr/) to identify GO biological process (BP) modules with minimal overlap. We only kept modules of 10 or more genes for our analysis. For PD GWAS genes, we identified six such modules (**Supplementary Table T12**) and for DA neuron essentiality genes, we identified 10 such modules (**Supplementary Table T15**). To impose network topology on these gene sets, we extracted edge relationships of genes in each group from three different data sources as described above: (i) high-confidence physical and genetics interactions from protein interaction databases, (ii) co-expression in the substantia nigra region of the mid-brain from GTEx v8, and (iii) co-essentiality in CNS cell types in DepMap v2023Q2.

We also explored an alternative approach from a recent study^65^ for constructing gene modules from PD GWAS loci, which involved running MAGMA^126^ analysis on GWAS genes, followed by GO BP enrichment. This approach identified 21 significant conditionally independent GO terms^65^. We then imposed network topology on the overlapping genes with each GO term by extracting gene-gene relationships from the sources described above.

### Human iPSCs culture and induced neuron di_erentiation

Human GM29371 iPSCs (Coriell Institute; GM29371*C) were cultured in Stemflex (Gibco/Thermo Fisher Scientific; Cat. No. A33493) on 6-well plates coated with Matrigel Matrix (Corning; Cat. No. 356231) diluted 1:100 in Knockout DMEM (Gibco/Thermo Fisher Scientific; Cat. No. 10829-018). Essential 8 Medium (Gibco/Thermo Fisher Scientific; Cat. No. A1517001) was replaced every day. When 80% confluent, cells were passaged with StemPro Accutase Cell Dissociation Reagent (Gibco/Thermo Fisher Scientific; Cat. No. A11105-01). Human iPSCs engineered to express *NGN2* under a doxycycline-inducible system in the AAVS1 safe harbor locus were differentiated following previously published protocol^86^. Briefly, iPSCs were released as above, centrifuged, and resuspended in N2 Pre-Differentiation Medium containing the following: Knockout DMEM/F12 (Gibco/Thermo Fisher Scientific; Cat. No. 12660-012) as the base, 1X MEM Non-Essential Amino Acids (Gibco/Thermo Fisher Scientific; Cat. No. 11140-050), 1X N2 Supplement (Gibco/Thermo Fisher Scientific; Cat. No. 17502-048), 10ng/mL NT-3 (PeproTech; Cat. No. 450-03), 10ng/mL BDNF (PeproTech; Cat. No. 450-02), 1 μg/mL Mouse Laminin (Thermo Fisher Scientific; Cat. No. 23017-015), 10nM ROCK inhibitor (Peprotech; Cat. No. 1293823), and 2μg/mL doxycycline hydrochloride (Sigma-Aldrich; Cat. No. D3447-500MG) to induce expression of mNGN2. iPSCs were counted and plated on Matrigel-coated plates in N2 Pre-Differentiation Medium for three days. After three days, hereafter DIV 0, pre-differentiated cells were released and centrifuged as above, and pelleted cells were resuspended in Classic Neuronal Medium containing the following: half DMEM/F12 (Gibco/Thermo Fisher Scientific; Cat. No. 11320-033) and half Neurobasal-A (Gibco/Thermo Fisher Scientific; Cat. No. 10888-022) as the base, 1X MEM Non-Essential Amino Acids, 0.5X GlutaMAX Supplement (Gibco/Thermo Fisher Scientific; Cat. No. 35050-061), 0.5X N2 Supplment, 0.5X B27 Supplement (Gibco/Thermo Fisher Scientific; Cat. No. 17504-044), 10ng/mL NT-3, 10ng/mL BDNF, 1μg/mL Mouse Laminin, and 2μg/mL doxycycline hydrochloride. Pre-differentiated cells were subsequently counted and plated on BioCoat Poly-D-Lysine coated plates (Corning; Cat. No. 356470) in Classic Neuronal Medium. On DIV 7 and each week after, medium change was performed without doxycycline added. In the PFF exposure experiment, a complete medium change to the medium containing 10 µg/mL synthetic PFF was performed on DIV 21 neurons, and the neurons were fixed at DIV28 for immunostaining.

### CRISPRi screen and analysis in the CiS-CN model

A customized CRISPRi library comprising 9,852 sgRNAs targeting 1,705 physical and genetic interactors of α-synuclein (αS)—including genes from the TransposeNet αS proteinopathy network—along with negative controls, was constructed. The library was packaged into lentivirus by the Virus Core at Boston Children’s Hospital. Neurons were transduced with the viral library and cultured until the harvesting date (DIVs 3, 28, and 42). Genomic DNA was extracted from harvested neurons, and the sgRNA-encoding regions were amplified for sequencing based on previously described protocols^86,127,128^. Sequencing and annotation were conducted at Memorial Sloan Kettering Cancer Center. Data was analyzed using the *MAGeCK-iNC* pipeline^86^ from the *MAGeCK* package (v0.5.9.2) using python (v2.7). Hits were classified as having an FDR-adjusted p-value < 0.1, and the gene product cutoff was selected by the *MAGeCK-iNC* pipeline dynamically for each comparison. Hits with significant sgRNA dropout were termed “toxicity enhancers.” Details of the functional screen in its entirety will be described in a forthcoming publication from our group.

### Immunostaining and microscopy imaging

Neurons are fixed with 4% PFA (EM Sciences; Cat. No. 15710) for 15 minutes at room temperature, and then permeabilized with 0.5% Triton X-100 (Sigma-Aldrich; Cat. No. T8787) and blocked with 0.05% Triton X-100 and 5% BSA (Sigma-Aldrich; Cat. No. A7906) in DPBS (Thermo Fisher Scientific; Cat. No. 14040182) for 1hr at room temperature. Samples were incubated with primary antibodies (PRL: Thermo Fisher Scientific; Cat. No. MA1-10597; 1:200 dilution) at 4°C overnight, followed by incubation with secondary antibody and Hoechst 33342 (Thermo Fisher Scientific; Cat. No. H3570; 1:2000 dilution) for 1hr at room temperature. Images were captured with identical settings for parallel cultures using Nikon Eclipse Ti microscope or Nikon TiE/C2 confocal microscope. Image analysis was performed with ImageJ2^129^ Macro Software (**Supplementary Note**). *Prl* level was determined by D (D = total *prolactin* intensity / total DAPI number in a given image).

### Oxidative stress assay

DIV0 CiS neurons were seeded at a density of 40,000 cells/well of poly-L-ornithine-coated 96-well plate (solution: Sigma-Aldrich; P4957). At DIV6, the neuron media was fully changed for 100uL *Prolactin*-containing (Thermo Fisher Scientific; Cat. No. 100-07-10UG) media at a concentration of 0,1 or 10 nM. 24hrs post-treatment, the neuron media was fully changed for 100uL of Menadione-containing (Thermo Fisher Scientific; Cat. No. ICN10225925) media at a concentration of 0 or 100uM and incubated for an hour at 37°C. After an hour, CELLROX™ Green Reagent (Thermo Fisher Scientific; Cat. No. C10444) was added on top of the Menadione-treated media to a final concentration of 5 uM CELLROX™ for 30 minutes at 37°C. All media was then removed, and wells were washed 3 times with PBS. After the third wash, the PBS was replaced with neuron media. The plates were then taken to the Incucyte^®^ S3 live-cell analysis system (Sartorius) for imaging. Incucyte analysis was perform with S3 software, and CellRox = total integrated intensity/neuron was reported.

### Mouse model

Wildtype B6C3F1 mice (The Jackson Laboratories; Stock 100010) were used for the stereotactic injection studies described. Animals were maintained on a 12-hour light/dark schedule and provided with food ad libitum. All housing, breeding, and procedures were performed according to the NIH Guide for the Care and Use of Experimental Animals and approved by the University of Pennsylvania Institutional Animal Care and Use Committee.

### ɑS and PFF preparation

Full-length mouse ɑS was expressed in BL21 (DE3) RIL-competent E. coli cells (Agilent Technologies; Cat. No. 230245) transformed with pRK172/mSyn containing ɑS cDNA^29^. Protein purification was previously described^29,130^. Cultures expanded in Terrific Broth (Sigma-Aldrich; Cat. No. T9179, composition: 12 g/L of Bacto-tryptone, 24 g/L of yeast extract 4% (vol/vol) glycerol, 17 mM KH2PO4 and 72 mM K2HPO4) containing ampicillin (Fisher Scientific; Cat. No. BP1760) were harvested and sonicated in high salt buffer (750 mM NaCl in 10 mM Tris, pH 7.6; NaCl: Fisher Scientific, Cat. No. S271; Tris: Fisher Scientific, Cat. No. BP152). After boiling for 15 mins, the supernatant was dialyzed against 10 mM Tris, pH 7.6, 50 mM NaCl, 1 mM EDTA (Sigma-Aldrich; Cat. No. E5134) overnight at 4°C, filtered and concentrated using Amicon Ultra-15 centrifugal filter units (Merck Millipore; Cat. No. UFC901008). Gel filtration using a Superdex 200 column (Cytiva; Cat. No. 17517501) was performed and fractions containing ɑS pooled and dialyzed in 10 mM Tris, pH 7.6, 50 mM NaCl, 1 mM EDTA overnight. The product was polished using a HiTrapQ HP column (Cytiva; Cat. No. 645932) and eluted over an ionic gradient (25 to 1,000 mM NaCl). Fractions containing ɑS were combined and dialyzed into DPBS (Invitrogen; Cat. No. 14200075), sterile filtered and concentrated to 5 mg/mL and frozen at-80°C until used. PFFs were assembled by shaking monomer at 5 mg/mL using a Thermomixer C (Eppendorf) set at 1,000 rpm for 7 days at 37°C. Fibril content was validated by sedimentation at 100,000 x g for 30 minutes and Thioflavin T fluorimetry.

### Stereotaxic administration of PFFs

Prior to injection, PFFs were diluted to 2 mg/mL in DPBS and sonicated using a bath sonicator (Diagenode; Biorupter UCD-300) on high power for 10 cycles (30 sec on; 30 sec oi) at 10°C. Each mouse received a single unilateral injection of PFFs (5 µg of PFFs in 2.5 µL volume) into the dorsal striatum using a Hamilton syringe (33 gauge) using the following co-ordinates: AP +0.2 mm relative to Bregma; ML +2.0 mm; depth 2.6 mm beneath the dura. DPBS injected into the same region was used as a negative control. Mice were perfused transcardially with heparinized PBS at 30 d.p.i. and brains flash frozen at-80°C until use.

### RNA isolation and NanoString analysis

The amygdala region ipsilateral to PFF-or PBS injection was microdissected from each brain and homogenized in 1 ml of TRI Reagent (Sigma-Aldrich, Cat. No. T9424) using TissueRuptor II (Qiagen, Cat. No. 9002755) with a disposable probe (Qiagen, Cat. No. 990890). RNA was then isolated using Direct-zol RNA MiniPrep kit with in-column DNaseI treatment (Zymo; Cat. No. R2050). Samples were quantitated with a NanoDrop 1000 Spectrophotometer (Thermo Fisher Scientific) and assayed for RNA integrity on a 4200 TapeStation (Agilent; Cat. No. G2991AA). NanoString hybridization of the resultant RNA was carried out for a constant 18 hours at 65° C on the *Mus musculus* Neuropathology panel (v1.0). Post-hybridization processing in the nCounter Prep Station used the High Sensitivity settings. The cartridge scanning parameter was set at high (555 FOV). RNA isolation and NanoString studies were performed at the Wistar Genomics core facility. Genes with hybridization counts >20 were normalized to the geometric mean of panel positive controls as recommended by the manufacturer. Differential expression analysis was performed using a generalized linear model in the nCounter module in Rosalind (Rosalind.bio). Adjusted p-values were calculated using the Benjamini-Hochberg method using treatment (i.e. PFF vs PBS).

## Data and code availability

Canonical pathway gene set were obtained from MSigDB^131^ (v7.3; https://www.gsea-msigdb.org/gsea/msigdb/human/collections.jsp). Human physical protein–protein interactions (PPI) were downloaded from STRING^132^ (v11.5; https://string-db.org/cgi/download), HuRI (http://www.interactome-atlas.org/download: last accessed on Jan 2022), and inBio Map^112^ (https://www.intomics.com/inbio/map: last accessed on Jan 2022) databases. Additionally, genetic interactions data was obtained from the supplement of the Megchelenbrink *et al.* study^113^. TransposeNet’s humanized *α*-synuclein-, *β*-amyloid-, and TDP-43-modifier networks were obtained from our previously published study^2^. Bulk expression data in TPM format from different human tissue types were downloaded from the GTEx^114^ (v8; https://www.gtexportal.org/), and gene dependency data in different cell lines from DepMap^116^ (release: 2023Q2; https://depmap.org/portal/data_page/?tab=allData).

Whole exome sequencing (WES) and phenotypic data from UKBB^133^, available through https://ams.ukbiobank.ac.uk, were accessed via application 41250 and processed on the DNAnexus platform (https://ukbiobank.dnanexus.com/landing). MGBBB^134^ WES and phenotypic data were accessed via https://biobankportal.partners.org/ (PI: Khurana). Access to MGBBB data was restricted to only affiliated investigators. AMP-PD^135^ whole genome sequencing and phenotypic data (v2.5; release 2022) were accessed through the AMP-PD Knowledge Platform (https://www.amp-pd.org).

NERINE’s source code is available on github (https://github.com/snz20/NERINE) and zenodo (doi: 10.5281/zenodo.19209293). RVTT was run adapting the code from https://github.com/snz20/RVTT (zenodo doi: 10.5281/zenodo.10627549). CMC-Fisher, Fisher’s combined test, and SKAT-O were run on R (v4.3.2) using stats (v4.3.2), poolr (v1.2.0), and SKAT (v2.2.5) packages. MAGeCK-iNC analysis was performed using the MAGeCK^136^ (v0.5.9.2) package on python (v2.7) and GO gene set enrichment analysis was performed using GSEApy (v1.1.3) on python (v3.12.4). Comprehensive information on study-related resources is listed in **Supplementary Table T24**.

## Supporting information

Supplementary Figures with Legends

## Acknowledgements

We thank Drs. Matthew Stevens, Richard Sherwood, Benjamin Neale, and Isabel Lam for their valuable insights. S.N. is supported by the NIH grant R35GM127131, the Sudarsky Scholar Award (Brigham and Women’s Hospital, Movement Disorders Division), and the Australian Parkinson’s Mission. S.S. is supported by NIH grants U01HG012009, R35GM127131, and R01MH101244. V.K, X.W., L.S., and experiments in PD neuronal models are supported by the Aligning Science Across Parkinson’s Initiative (ASAP) award ASAP-000472 (PI: Studer, co-PI: Khurana). V.K. also acknowledges support from an APDA Center for Advanced Research grant, the NIH grant R01NS109209, the New York Stem Cell Foundation Robertson Investigator award (NYSCF-R-I49), as well as the generous support from Mrs. Nancy Black Simches and the Ocko family Parkinson’s Disease Innovation Award.

X.W. additionally acknowledges support from the NIH grant T32AG000222 (PI: Yankner).

We analyzed WES data from the UK Biobank (UKBB, application 41250, PI: Cassa; Mass General Brigham IRB 2020P002093; NIH R01HG010372, PI: Sunyaev), a global biomedical resource supported by the Wellcome Trust, UK Medical Research Council, the Department of Health, Scottish Government, the Northwest Regional Development Agency, British Heart Foundation, and Cancer Research UK. We also used WES and phenotypic data from ∼50k participants from the Mass General Brigham Biobank (MGBBB), a biorepository of consented patient samples at Mass General Brigham (parent organization of Massachusetts General Hospital and Brigham and Women’s Hospital), and WGS data from AMP-PD, a public-private partnership managed by FNIH and funded by Celgene, GSK, the Michael J. Fox Foundation, NINDS, Pfizer, and Verily. AMP-PD investigators did not review this work. We thank all the participants, clinical investigators, and research teams who contributed to UKBB, MGBBB, and AMP-PD.

## Author contributions

S.N., V.K., and S.S. conceived of the project, interpreted the results; S.N., X.W., V.K., and S.S. wrote the manuscript; S.N. and S.S. developed the novel methodology NERINE; S.N. pre-processed and analyzed the study cohorts; A.M. assisted in the quality control and the pre-processing of the sequencing data; S.N., S.S., N.S. and R.G. interpreted the results in lipid and cardiac phenotypes; C.C. helped analyze the UKBB cohorts; X.W. and V.K. designed the wet laboratory experiments to validate the NERINE’s findings in PD in neuronal models; X.W., R.S., and E.E. generated the CiS neurons and performed the experiments; X.W. and S.N. analyzed the experimental data and interpreted the results with V.K.; D.R., A.H., J.A., and L.S. designed the genome-wide CRISPR/Cas9 experiments in mid-brain DA neurons, identified the essentiality genes, and curated the GO biological process modules; K.L. designed and performed the experiments in the mouse PFF model, analyzed the data, and interpreted the results with V.K., X.W., and S.N.. All the authors read and helped edit the manuscript.

