## Supplementary Figures with Legends for "NERINE reveals rare variant associations in gene networks across phenotypes and implicates an *SNCA-PRL-LRRK2* subnetwork in Parkinson’s disease"

### Figure S1

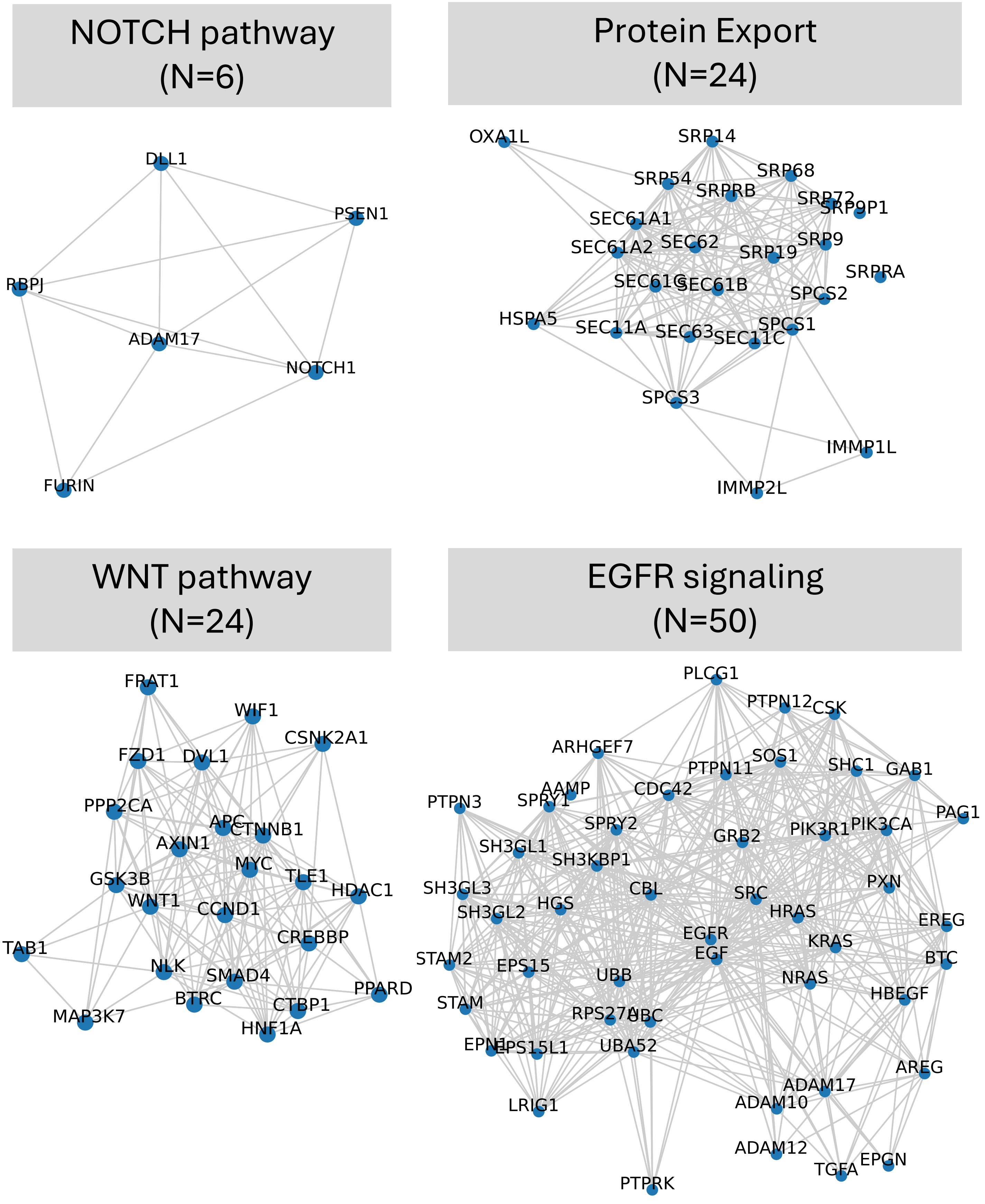
NOTCH pathway
(N=6)
DLL1
PSEN1
RBPJ
ADAM17
NOTCH1
FURIN
Protein Export
(N=24)
OXA1L
SRP14
SRP54
SRP68
SRPRB
SRP72
SRP9P1
SEC61A1
SEC61A2
SEC62
SRP19
SRP9
SRPRA
SEC61G
SEC61B
SPCS2
HSPA5
SEC11A
SEC63
SEC11C
SPCS1
SPCS3
IMMP1L
IMMP2L
WNT pathway
(N=24)
FRAT1
WIF1
CSNK2A1
FZD1
DVL1
PPP2CA
APC
CTNNB1
AXIN1
MYC
TLE1
GSK3B
HDAC1
WNT1
CCND1
CREBBP
TAB1
NLK
SMAD4
PPARD
BTRC
CTBP1
MAP3K7
HNF1A
EGFR signaling
(N=50)
PLCG1
PTPN12
CSK
ARHGEF7
SOS1
PTPN11
SHC1
GAB1
AAMP
CDC42
PTPN3
SPRY1
PAG1
SPRY2
PIK3R1
PIK3CA
GRB2
SH3GL1
SH3KBP1
PXN
SH3GL3
CBL
SRC
HGS
SH3GL2
HRAS
EREG
EGFR
KRAS
EGF
STAM2
BTC
EPS15
UBB
NRAS
HBEGF
STAM
RPS27A
UBC
EPN1
EPS15L1
UBA52
AREG
ADAM17
ADAM10
LRIG1
ADAM12
EPGN
TGFA
PTPRK

**Supplementary Figure S1. Well-studied canonical pathway networks used for testing NERINE's performance in simulations.** Four well-studied pathways of different sizes: the NOTCH pathway (m=6), the WNT pathway (m=24), the protein export pathway (m=24), and the EGFR signaling pathway (m=50) were used for simulations. Canonical pathway gene lists were extracted from MSigDB v7.3, and high-confidence physical and genetic interactions from protein-protein interaction (PPI) databases were used as network edges between pathway genes (see Methods).

### Figure S2

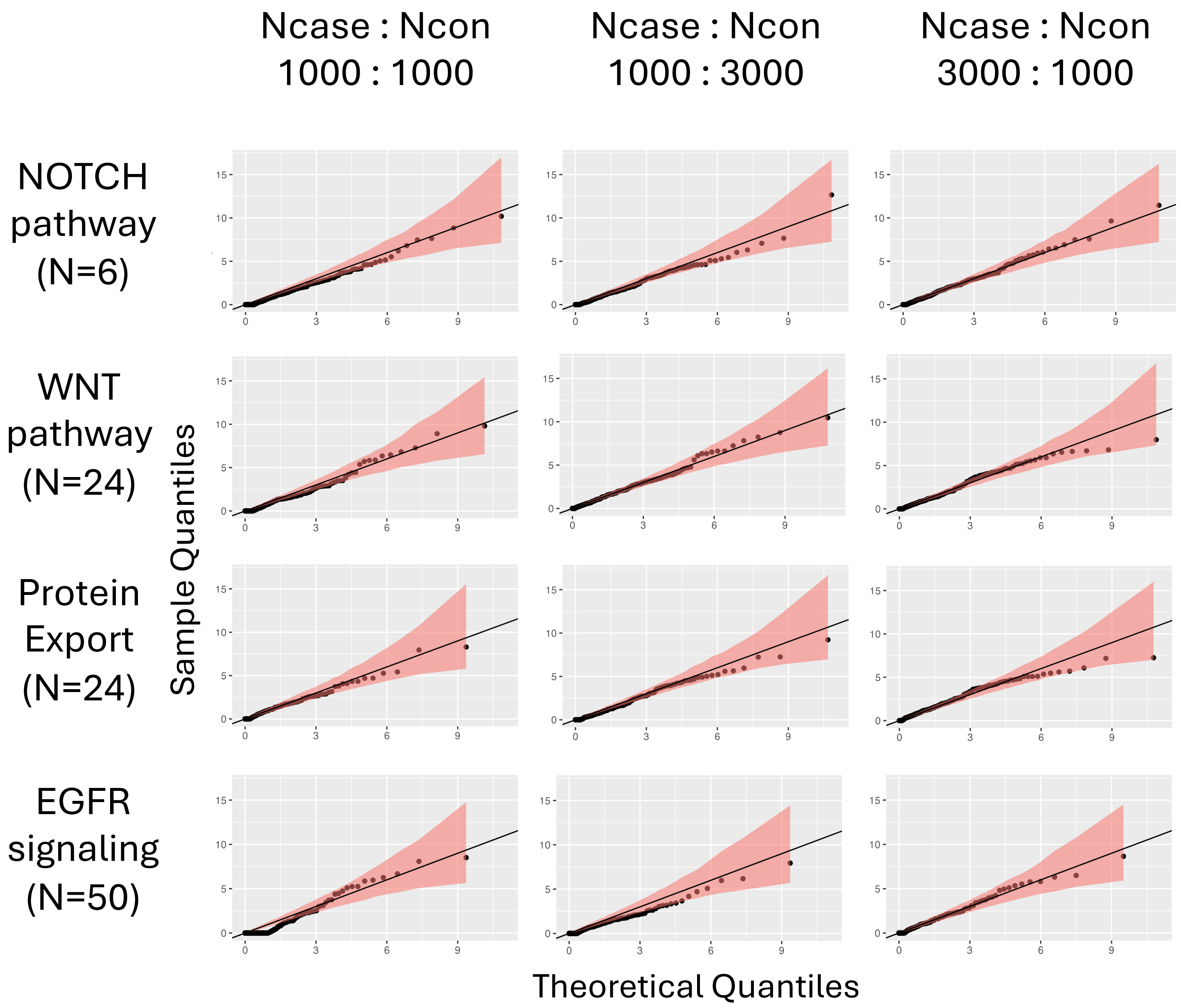

Ncase : Ncon
1000 : 1000
Ncase : Ncon
1000 : 3000
Ncase : Ncon
3000 : 1000
NOTCH pathway (N=6)
WNT pathway (N=24)
Protein Export (N=24)
EGFR signaling (N=50)
Sample Quantiles
Theoretical Quantiles

**Supplementary Figure S2. NERINE's performance in simulations at null with well-studied database pathway gene networks.** NERINE's test statistic asymptotically follows the theoretical distribution of a mixture of the delta function and a chi-square distribution with a degree of freedom of one. Simulations were performed with different network architectures for well-studied pathways of different sizes: the NOTCH pathway, the WNT pathway, the protein export pathway, and the EGFR signaling pathway. Pathway gene lists were extracted from MSigDB v7.3, and high-confidence physical and genetic interactions from protein-protein interaction (PPI) databases were used as network edges between pathway genes (see Methods). The allele counts in cases and controls were generated from independent binomial distributions. Simulations were performed in cohorts with different case-control skews. For each scenario, 1,000 iterations were performed to create the QQ plots. Confidence bands in the QQ plots represent 95% bootstrap confidence intervals around NERINE's test statistic.

### Figure S3

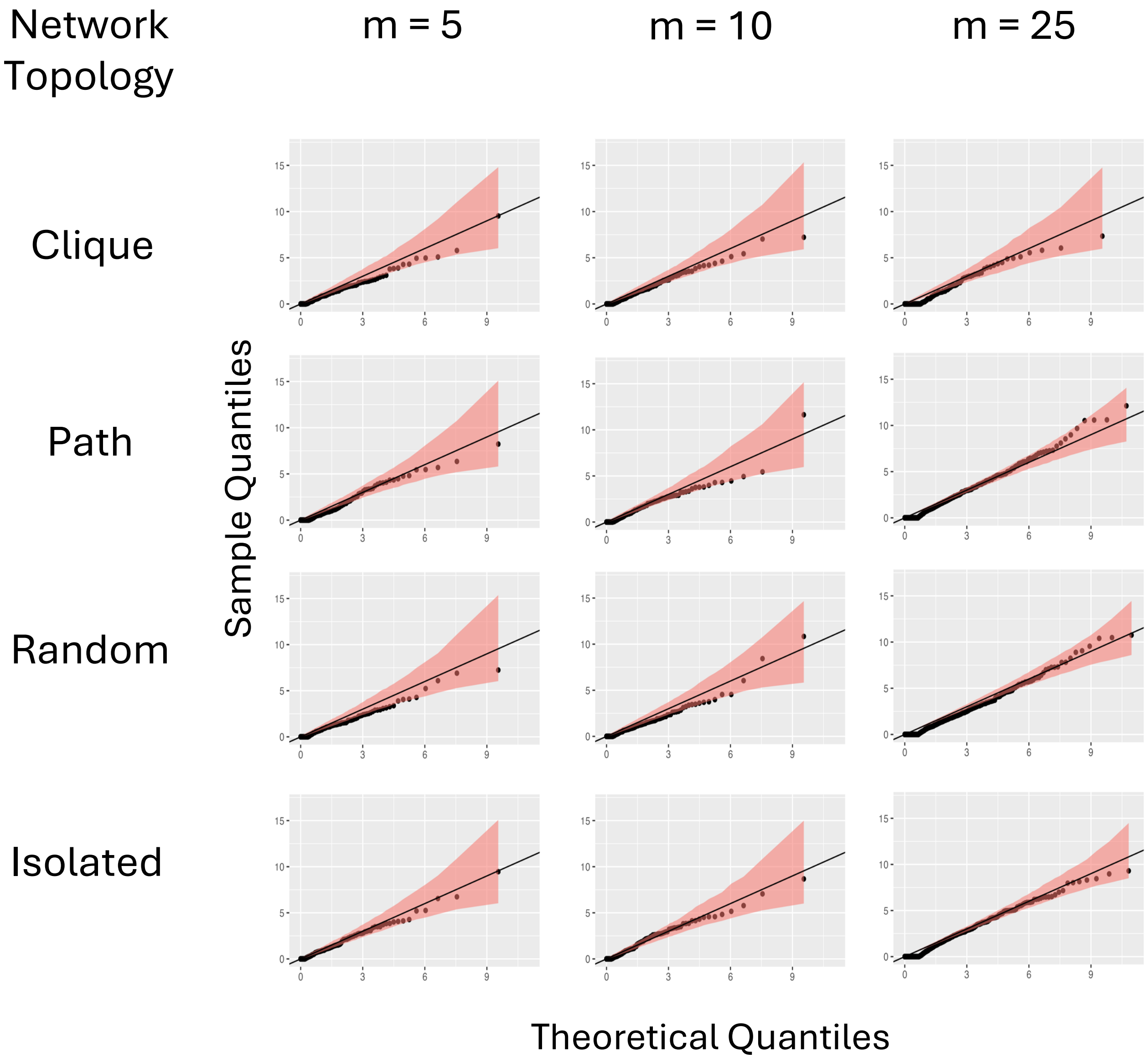
Network Topology
m = 5
m = 10
m = 25
Clique
Path
Random
Isolated
Sample Quantiles
Theoretical Quantiles

**Supplementary Figure S3. NERINE's performance in simulations at null with simulated network topologies.** Simulations were performed using networks of different sizes (i.e, 5, 10, and 25 genes) and different topological architectures (i.e., clique, path, random, and isolated nodes) for equal-sized case- and control-groups. For each scenario, 1,000 iterations were performed to generate the QQ plots. The allele counts in cases and controls were generated from independent binomial distributions. Confidence bands in the QQ plots represent 95% bootstrap confidence intervals around NERINE's test statistic. In each scenario, NERINE's test statistic asymptotically follows the theoretical distribution of a mixture of the delta function and a chi-square distribution with a degree of freedom of one.

### Figure S4

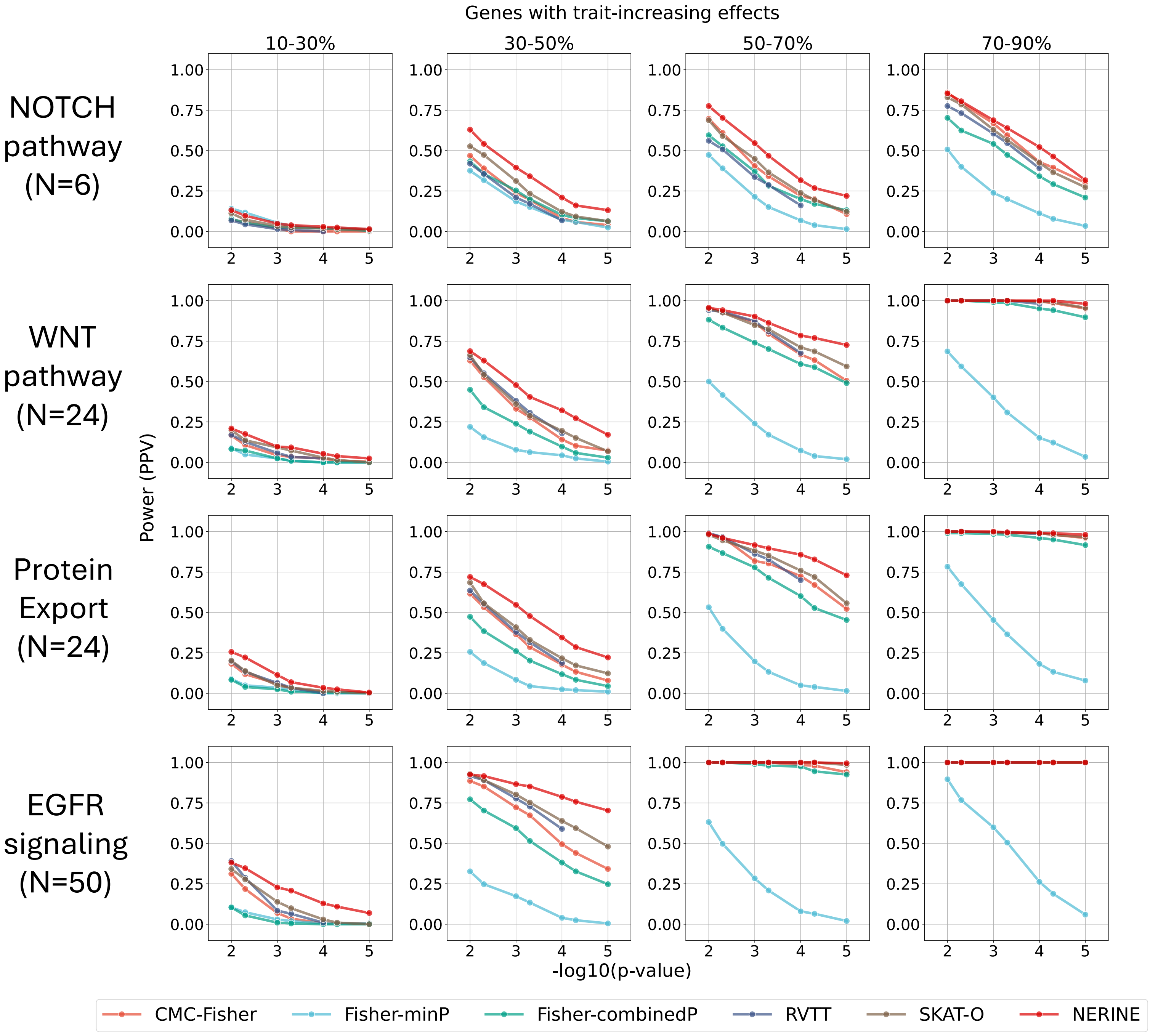

Genes with trait-increasing effects
10-30%
30-50%
50-70%
70-90%
NOTCH pathway (N=6)
WNT pathway (N=24)
Protein Export (N=24)
EGFR signaling (N=50)
Power (PPV)
-log10(p-value)
CMC-Fisher
Fisher-minP
Fisher-combinedP
RVTT
SKAT-O
NERINE

**Supplementary Figure S4. NERINE outperforms existing rare variant association tests in power simulations when relevant genes have only trait-increasing effects.** The empirical power of the methods was measured for a simulated binary disease trait in a cohort of 2,000 cases and 2,000 controls with network effect in four well-studied database pathways. NERINE was tested in only the scenarios where different proportions of genes in the network had only trait-increasing effects. From left to right, the power plots show those scenarios, mimicking situations from having a very noisy network to a highly relevant one. Power was calculated as the positive predictive value at different p-value cutoffs: 1e-2, 5e-3, 1e-3, 5e-4, 1e-4, 5e-5, and 1e-5. In all scenarios, NERINE outperforms existing rare variant tests. Note that RVTT p-values were calculated from 10,000 permutations. Hence, we don't report RVTT's power for the cutoffs below 1e-4.

### Figure S5

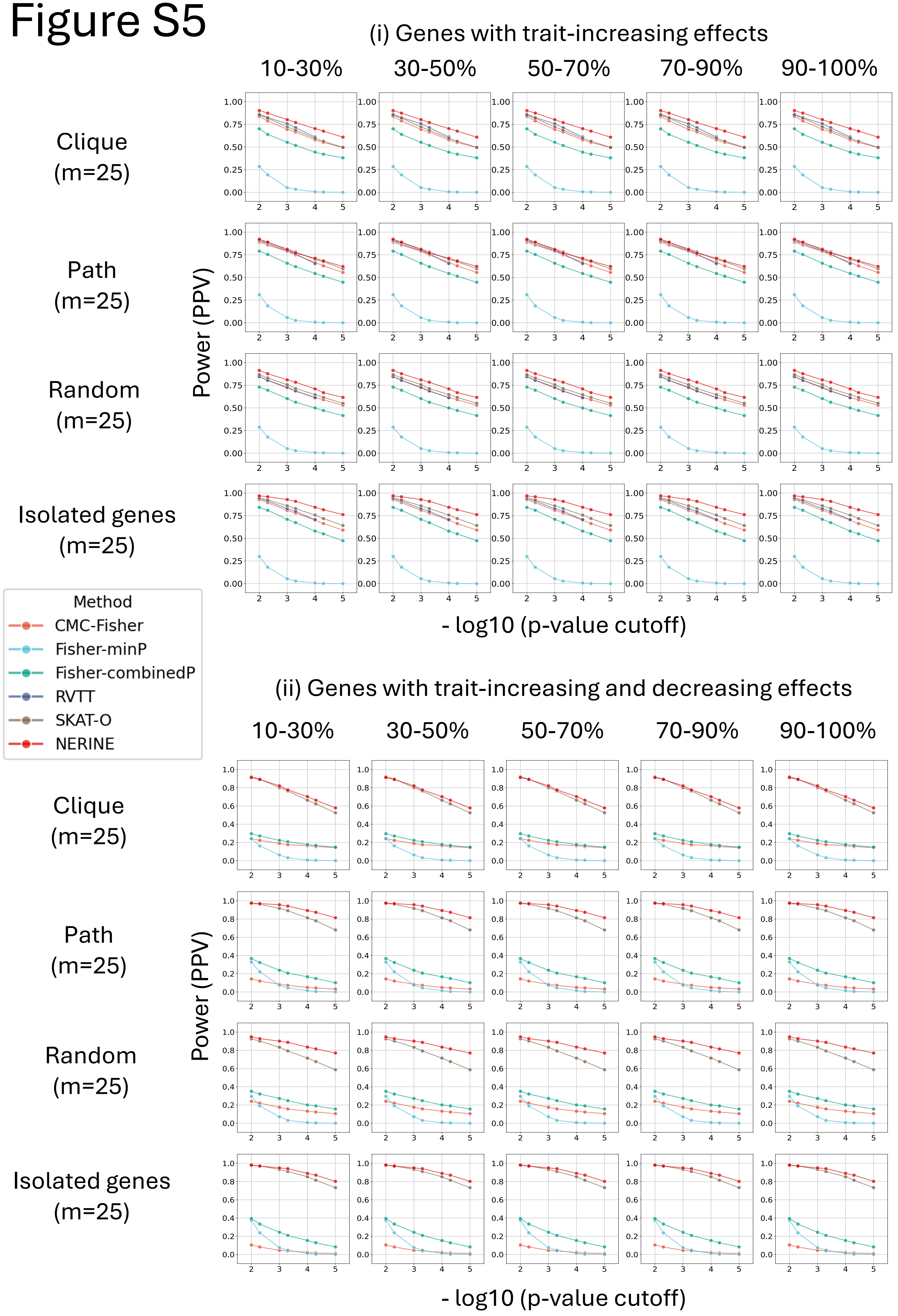
(i) Genes with trait-increasing effects
10-30%
30-50%
50-70%
70-90%
90-100%
Clique (m=25)
Path (m=25)
Random (m=25)
Isolated genes (m=25)
Power (PPV)
- log10 (p-value cutoff)
Method
CMC-Fisher
Fisher-minP
Fisher-combinedP
RVTT
SKAT-O
NERINE
(ii) Genes with trait-increasing and decreasing effects
10-30%
30-50%
50-70%
70-90%
90-100%
Clique (m=25)
Path (m=25)
Random (m=25)
Isolated genes (m=25)
Power (PPV)
- log10 (p-value cutoff)

**Supplementary Figure S5. NERINE outperforms existing rare variant association tests in power simulations with simulated network topologies.** We measured the empirical power of the methods for a simulated binary disease trait in a cohort of 1,000 cases and 1,000 controls with network effect for networks with different topological architectures (i.e., clique, path, random, and isolated nodes) in two scenarios: (i) genes in the network having only trait-increasing effects, and (ii) genes in the network having both trait-increasing and trait-decreasing effects. From left to right, the power plots show settings in which different proportions of genes within the network affect the trait, mimicking the scenarios from having a very noisy network to a highly relevant one. Power was calculated as the positive predictive value at different p-value cutoffs: 1e-2, 5e-3, 1e-3, 5e-4, 1e-4, 5e-5, and 1e-5. In all scenarios, NERINE outperforms existing rare variant tests. Note that RVTT p-values were calculated from 10,000 permutations. Hence, we don't report RVTT's power for the cutoffs below 1e-4. Also, RVTT is a test for monotonic trends in rare variant occurrences within a pathway. Hence, it was excluded from the comparison in scenario (ii).

### Figure S6

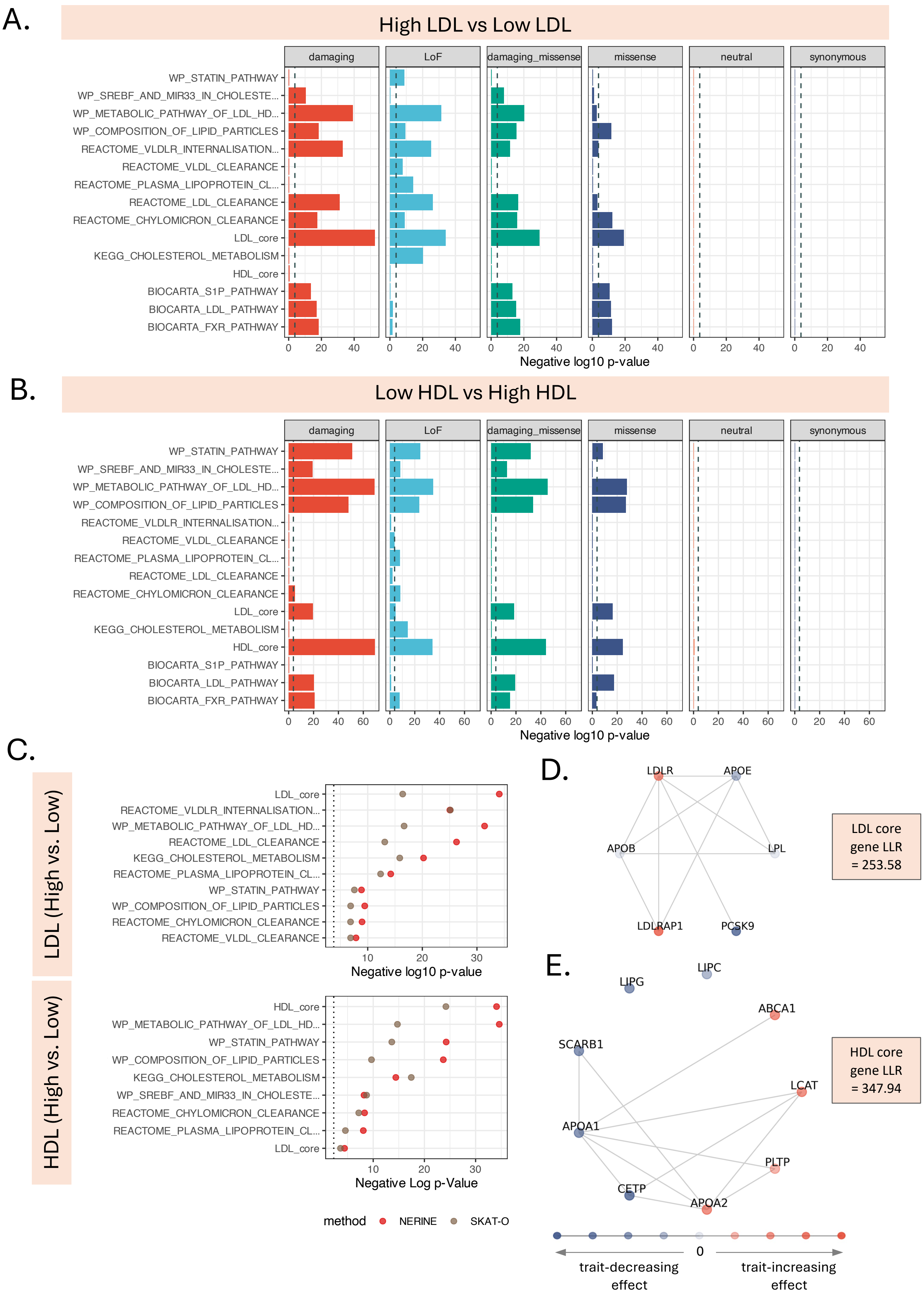

A.
High LDL vs Low LDL
damaging
LoF
damaging_missense
missense
neutral
synonymous
WP_STATIN_PATHWAY
WP_SREBF_AND_MIR33_IN_CHOLESTE...
WP_METABOLIC_PATHWAY_OF_LDL_HD...
WP_COMPOSITION_OF_LIPID_PARTICLES
REACTOME_VLDLR_INTERNALISATION...
REACTOME_VLDL_CLEARANCE
REACTOME_PLASMA_LIPOPROTEIN_CL...
REACTOME_LDL_CLEARANCE
REACTOME_CHYLOMICRON_CLEARANCE
LDL_core
KEGG_CHOLESTEROL_METABOLISM
HDL_core
BIOCARTA_S1P_PATHWAY
BIOCARTA_LDL_PATHWAY
BIOCARTA_FXR_PATHWAY
Negative log10 p-value
B.
Low HDL vs High HDL
Negative log10 p-value
C.
LDL (High vs. Low)
HDL (High vs. Low)
Negative log10 p-Value
Negative Log p-Value
method
NERINE
SKAT-O
D.
LDLR
APOE
APOB
LPL
LDLRAP1
PCSK9
LDL core gene LLR = 253.58
E.
LIPG
LIPC
ABCA1
SCARB1
LCAT
APOA1
PLTP
CETP
APOA2
HDL core gene LLR = 347.94
0
trait-decreasing effect
trait-increasing effect

**Supplementary Figure S6. Performance of NERINE while comparing high LDL-C vs. low LDL-C individuals and low HDL-C vs. high HDL-C individuals in the UK Biobank. A.** While comparing individuals with high LDL cholesterol with the ones with low LDL cholesterol in UKBB, NERINE identifies significant cumulative effect of rare (MAF < 0.001) variants in LoF (i.e., frameshifts, insertions, deletions, and splice), damaging missense (i.e., missenses predicted to be damaging by in-silico tools), damaging (i.e., damaging missense and LoF), and missense categories in key lipid-related pathways. No significant burden of neutral missense and synonymous variants was observed. The tests were performed across our canonical pathway database of 306 pathways. Pathway gene lists were extracted from MSigDB v7.3, and high-confidence physical and genetic interactions from protein-protein interaction (PPI) databases were used as network edges between pathway genes (see Methods). The dashed grey line represents the Bonferroni-corrected p-value threshold of 0.05. The core module of LDL genes, which contains *LDLR* and *PCSK9*, was identified as the most significant hit, which serves as a positive control. **B.** While comparing individuals with low HDL cholesterol with the ones with high HDL cholesterol in UKBB, NERINE identifies a significant cumulative effect of rare (MAF < 0.001) variants in LoF, damaging missense, damaging, and missense categories in key lipid-related pathways. Notably, the module of core HDL-related genes containing *ABCA1*, *CETP*, *LIPC*, and *LIPG* was the most significant hit, serving as a positive control. No significant burden of neutral missense and synonymous variants was observed. The tests were performed across the same canonical pathway database. The dashed grey line represents the Bonferroni-corrected p-value threshold of 0.05. **C.** Comparison of SKAT-O p-values against NERINE p-values for rare LoF variant burden across the significant pathways in the LDL (high vs low) and HDL (low vs high) phenotypes. For most of the pathways in both phenotypes, NERINE provides a lower p-value than SKAT-O. SKAT-O was applied at the pathway level, aggregating the allele counts from member genes. **D.** NERINE's estimates of gene effects in the most significant pathways with rare damaging variant burden in the LDL (high vs low) and HDL (low vs high) phenotypes. Trait-increasing effects are represented by shades of orange, and trait-decreasing effects are represented by shades of purple, as shown on the scale. A darker color represents a more pronounced effect. For example, *PCSK9* and *APOB* show trait-decreasing effects, and *LDLR* shows a trait-increasing effect on LDL cholesterol (high vs low) phenotype. Similarly, *ABCA1* and *LCAT* show trait-increasing effects, while *LIPC* and *LIPG* show trait-decreasing effects for the HDL (low vs high) phenotypes. These findings agree with known lipid biology. Note that NERINE's predicted gene effects represent the "most likely scenario" with the observed allele counts per gene and the gene-gene network topology under the estimated network effect. NERINE does not provide p-values on per-gene predictions.

### Figure S7

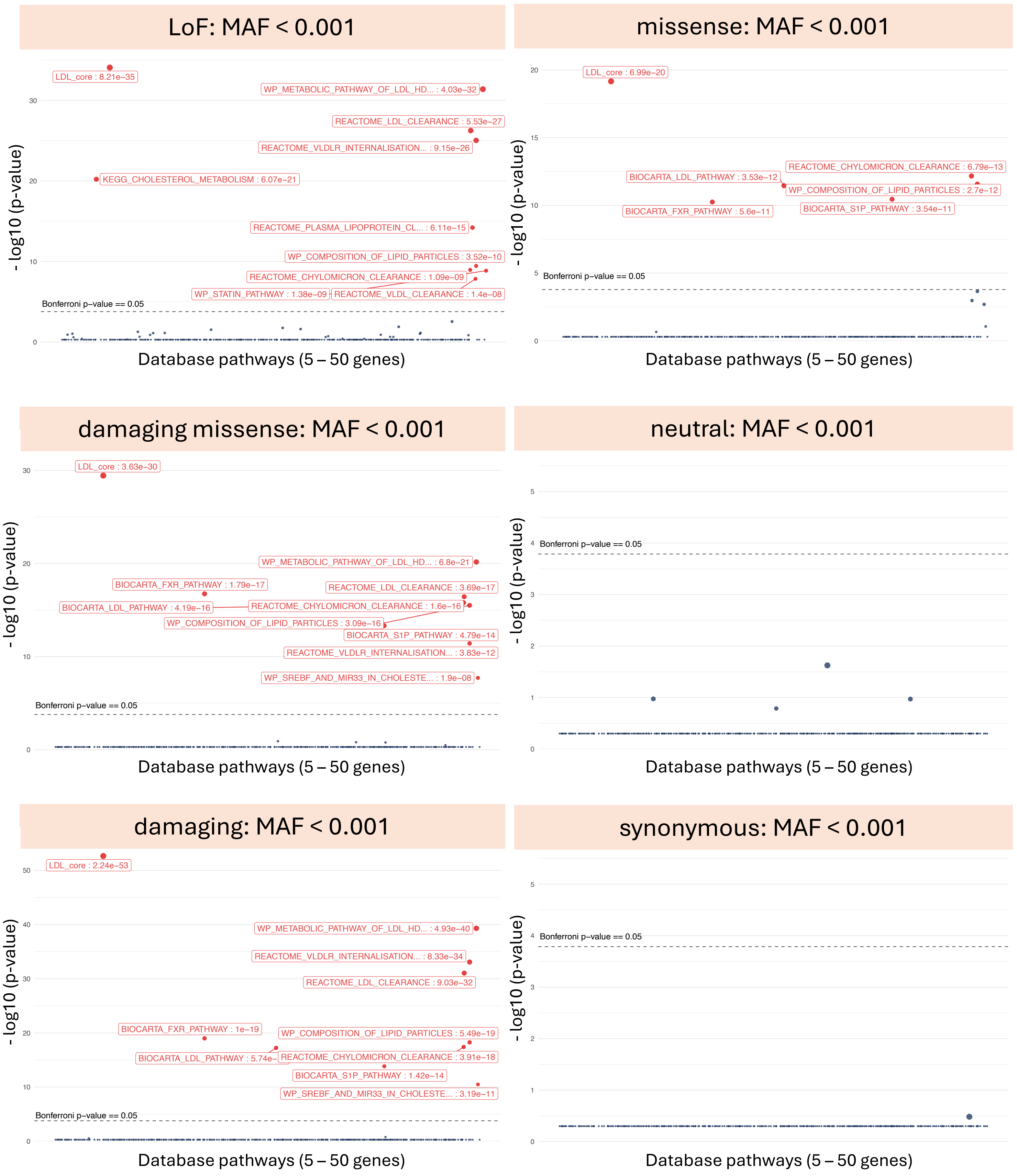

LoF: MAF < 0.001
LDL_core : 8.21e−35
WP_METABOLIC_PATHWAY_OF_LDL_HD... : 4.03e−32
REACTOME_LDL_CLEARANCE : 5.53e−27
REACTOME_VLDLR_INTERNALISATION... : 9.15e−26
KEGG_CHOLESTEROL_METABOLISM : 6.07e−21
REACTOME_PLASMA_LIPOPROTEIN_CL... : 6.11e−15
WP_COMPOSITION_OF_LIPID_PARTICLES : 3.52e−10
REACTOME_CHYLOMICRON_CLEARANCE : 1.09e−09
WP_STATIN_PATHWAY : 1.38e−09
REACTOME_VLDL_CLEARANCE : 1.4e−08
Bonferroni p−value == 0.05
- log10 (p-value)
Database pathways (5 – 50 genes)
missense: MAF < 0.001
LDL_core : 6.99e−20
REACTOME_CHYLOMICRON_CLEARANCE : 6.79e−13
BIOCARTA_LDL_PATHWAY : 3.53e−12
WP_COMPOSITION_OF_LIPID_PARTICLES : 2.7e−12
BIOCARTA_FXR_PATHWAY : 5.6e−11
BIOCARTA_S1P_PATHWAY : 3.54e−11
Bonferroni p−value == 0.05
- log10 (p-value)
Database pathways (5 – 50 genes)
damaging missense: MAF < 0.001
LDL_core : 3.63e−30
WP_METABOLIC_PATHWAY_OF_LDL_HD... : 6.8e−21
BIOCARTA_FXR_PATHWAY : 1.79e−17
REACTOME_LDL_CLEARANCE : 3.69e−17
BIOCARTA_LDL_PATHWAY : 4.19e−16
REACTOME_CHYLOMICRON_CLEARANCE : 1.6e−16
WP_COMPOSITION_OF_LIPID_PARTICLES : 3.09e−16
BIOCARTA_S1P_PATHWAY : 4.79e−14
REACTOME_VLDLR_INTERNALISATION... : 3.83e−12
WP_SREBF_AND_MIR33_IN_CHOLESTE... : 1.9e−08
Bonferroni p−value == 0.05
- log10 (p-value)
Database pathways (5 – 50 genes)
neutral: MAF < 0.001
Bonferroni p−value == 0.05
- log10 (p-value)
Database pathways (5 – 50 genes)
damaging: MAF < 0.001
LDL_core : 2.24e−53
WP_METABOLIC_PATHWAY_OF_LDL_HD... : 4.93e−40
REACTOME_VLDLR_INTERNALISATION... : 8.33e−34
REACTOME_LDL_CLEARANCE : 9.03e−32
BIOCARTA_FXR_PATHWAY : 1e−19
WP_COMPOSITION_OF_LIPID_PARTICLES : 5.49e−19
BIOCARTA_LDL_PATHWAY : 5.74e−
REACTOME_CHYLOMICRON_CLEARANCE : 3.91e−18
BIOCARTA_S1P_PATHWAY : 1.42e−14
WP_SREBF_AND_MIR33_IN_CHOLESTE... : 3.19e−11
Bonferroni p−value == 0.05
- log10 (p-value)
Database pathways (5 – 50 genes)
synonymous: MAF < 0.001
Bonferroni p−value == 0.05
- log10 (p-value)
Database pathways (5 – 50 genes)

**Supplementary Figure S7. Database pathway gene networks with significant rare variant burden identified by NERINE for LDL-C phenotype in UKBB.** Pathway Manhattan plots showing NERINE's results in six functional categories of variants: LoF (i.e., frameshifts, insertions, deletions, and splice variants), damaging missense (i.e., missenses predicted to be damaging by in-silico tools), damaging (i.e., LoF and damaging missenses), missense, neutral (i.e., missenses predicted to be benign by in-silico tools), and synonymous. Applying NERINE across a database of 306 canonical pathways to compare individuals with high LDL cholesterol with the ones with low LDL cholesterol in UKBB, we identified a significant cumulative effect of rare (MAF < 0.001) variants in LoF, damaging missense, damaging, and missense categories in key lipid-related pathways, such as the core module of LDL-related genes. No significant burden of neutral missense or synonymous variants was observed. The horizontal dashed line represents the Bonferroni-corrected p-value cutoff of 0.05.

### Figure S8

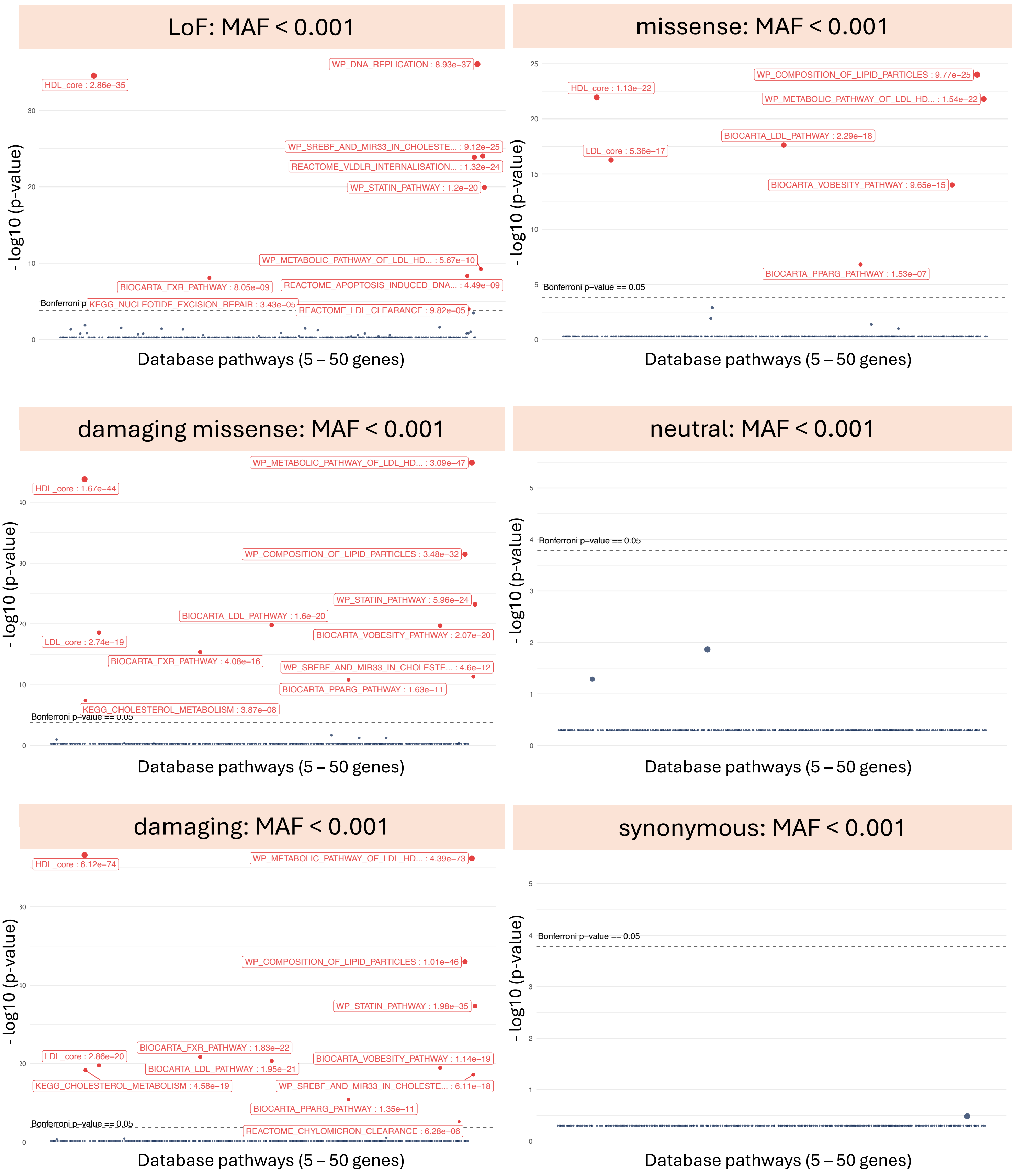

LoF: MAF < 0.001
WP_DNA_REPLICATION : 8.93e−37
HDL_core : 2.86e−35
WP_SREBF_AND_MIR33_IN_CHOLESTE... : 9.12e−25
REACTOME_VLDLR_INTERNALISATION... : 1.32e−24
WP_STATIN_PATHWAY : 1.2e−20
WP_METABOLIC_PATHWAY_OF_LDL_HD... : 5.67e−10
BIOCARTA_FXR_PATHWAY : 8.05e−09
REACTOME_APOPTOSIS_INDUCED_DNA... : 4.49e−09
KEGG_NUCLEOTIDE_EXCISION_REPAIR : 3.43e−05
REACTOME_LDL_CLEARANCE : 9.82e−05
Bonferroni p
- log10 (p-value)
Database pathways (5 – 50 genes)
missense: MAF < 0.001
WP_COMPOSITION_OF_LIPID_PARTICLES : 9.77e−25
HDL_core : 1.13e−22
WP_METABOLIC_PATHWAY_OF_LDL_HD... : 1.54e−22
BIOCARTA_LDL_PATHWAY : 2.29e−18
LDL_core : 5.36e−17
BIOCARTA_VOBESITY_PATHWAY : 9.65e−15
BIOCARTA_PPARG_PATHWAY : 1.53e−07
Bonferroni p−value == 0.05
- log10 (p-value)
Database pathways (5 – 50 genes)
damaging missense: MAF < 0.001
WP_METABOLIC_PATHWAY_OF_LDL_HD... : 3.09e−47
HDL_core : 1.67e−44
WP_COMPOSITION_OF_LIPID_PARTICLES : 3.48e−32
WP_STATIN_PATHWAY : 5.96e−24
BIOCARTA_LDL_PATHWAY : 1.6e−20
BIOCARTA_VOBESITY_PATHWAY : 2.07e−20
LDL_core : 2.74e−19
BIOCARTA_FXR_PATHWAY : 4.08e−16
WP_SREBF_AND_MIR33_IN_CHOLESTE... : 4.6e−12
BIOCARTA_PPARG_PATHWAY : 1.63e−11
KEGG_CHOLESTEROL_METABOLISM : 3.87e−08
Bonferroni p−value == 0.05
- log10 (p-value)
Database pathways (5 – 50 genes)
neutral: MAF < 0.001
Bonferroni p−value == 0.05
- log10 (p-value)
Database pathways (5 – 50 genes)
damaging: MAF < 0.001
HDL_core : 6.12e−74
WP_METABOLIC_PATHWAY_OF_LDL_HD... : 4.39e−73
WP_COMPOSITION_OF_LIPID_PARTICLES : 1.01e−46
WP_STATIN_PATHWAY : 1.98e−35
BIOCARTA_FXR_PATHWAY : 1.83e−22
LDL_core : 2.86e−20
BIOCARTA_LDL_PATHWAY : 1.95e−21
BIOCARTA_VOBESITY_PATHWAY : 1.14e−19
KEGG_CHOLESTEROL_METABOLISM : 4.58e−19
WP_SREBF_AND_MIR33_IN_CHOLESTE... : 6.11e−18
BIOCARTA_PPARG_PATHWAY : 1.35e−11
Bonferroni p−value == 0.05
REACTOME_CHYLOMICRON_CLEARANCE : 6.28e−06
- log10 (p-value)
Database pathways (5 – 50 genes)
synonymous: MAF < 0.001
Bonferroni p−value == 0.05
- log10 (p-value)
Database pathways (5 – 50 genes)

**Supplementary Figure S8. Database pathway gene networks with significant rare variant burden identified by NERINE for HDL-C phenotype in UKBB.** Pathway Manhattan plots showing NERINE's results in six functional categories of variants: LoF (i.e., frameshifts, insertions, deletions, and splice variants), damaging missense (i.e., missenses predicted to be damaging by in-silico tools), damaging (i.e., LoF and damaging missenses), missense, neutral (i.e., missenses predicted to be bening by in-silico tools), and synonymous. Applying NERINE across a database of 306 canonical pathways to compare individuals with low HDL cholesterol with the ones with high HDL cholesterol in UKBB, we identified a significant cumulative effect of rare (MAF < 0.001) variants in LoF, damaging missense, damaging, and missense categories in key lipid-related pathways, such as the core module of HDL-related genes. No significant burden of neutral missense or synonymous variants was observed. The horizontal dashed line represents the Bonferroni-corrected p-value cutoff of 0.05.

### Figure S9

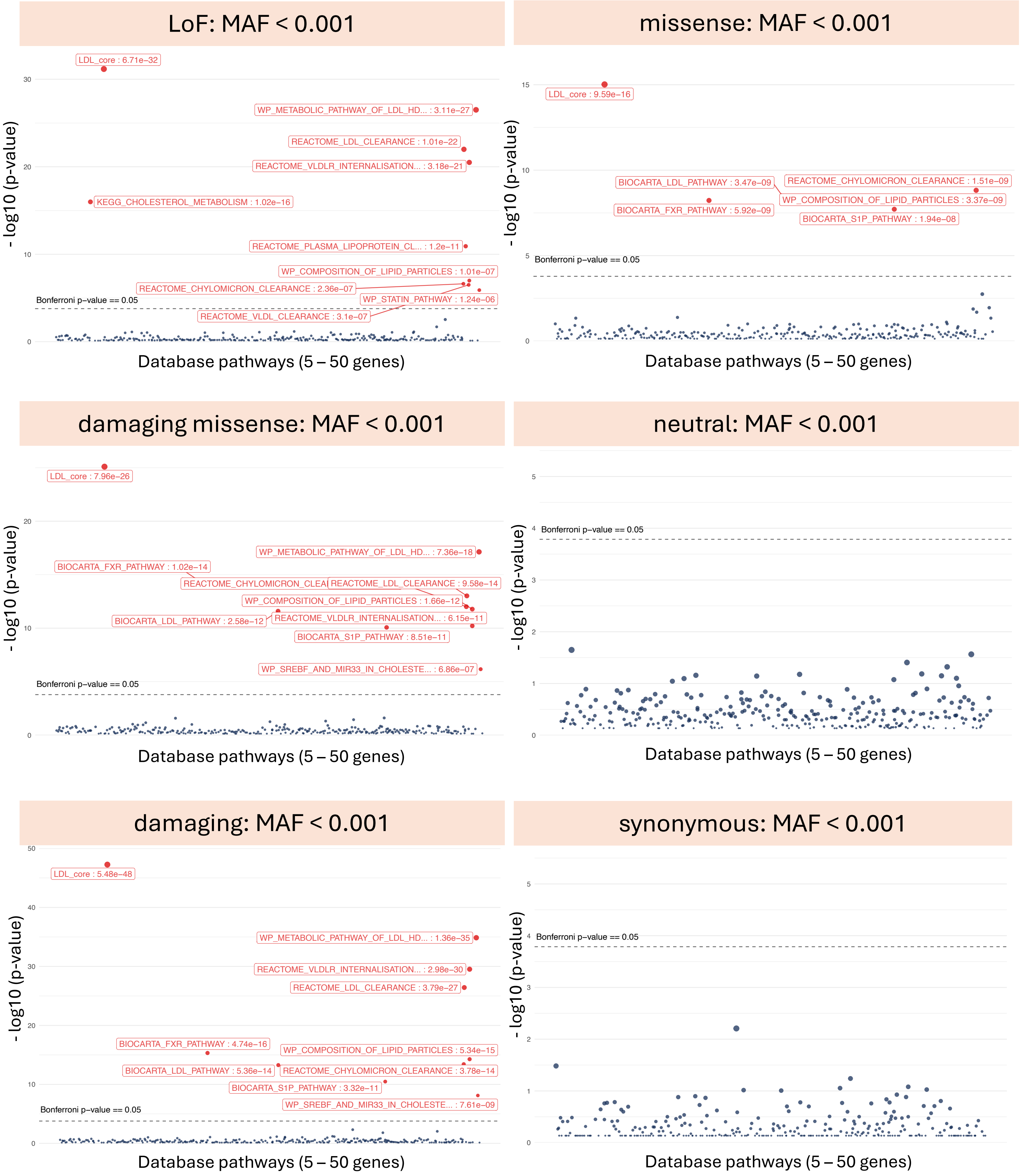
LoF: MAF < 0.001
LDL_core : 6.71e−32
WP_METABOLIC_PATHWAY_OF_LDL_HD... : 3.11e−27
REACTOME_LDL_CLEARANCE : 1.01e−22
REACTOME_VLDLR_INTERNALISATION... : 3.18e−21
KEGG_CHOLESTEROL_METABOLISM : 1.02e−16
REACTOME_PLASMA_LIPOPROTEIN_CL... : 1.2e−11
WP_COMPOSITION_OF_LIPID_PARTICLES : 1.01e−07
REACTOME_CHYLOMICRON_CLEARANCE : 2.36e−07
WP_STATIN_PATHWAY : 1.24e−06
REACTOME_VLDL_CLEARANCE : 3.1e−07
Bonferroni p−value == 0.05
- log10 (p-value)
Database pathways (5 – 50 genes)
missense: MAF < 0.001
LDL_core : 9.59e−16
BIOCARTA_LDL_PATHWAY : 3.47e−09
REACTOME_CHYLOMICRON_CLEARANCE : 1.51e−09
BIOCARTA_FXR_PATHWAY : 5.92e−09
WP_COMPOSITION_OF_LIPID_PARTICLES : 3.37e−09
BIOCARTA_S1P_PATHWAY : 1.94e−08
Bonferroni p−value == 0.05
- log10 (p-value)
Database pathways (5 – 50 genes)
damaging missense: MAF < 0.001
LDL_core : 7.96e−26
WP_METABOLIC_PATHWAY_OF_LDL_HD... : 7.36e−18
BIOCARTA_FXR_PATHWAY : 1.02e−14
REACTOME_CHYLOMICRON_CLEA
REACTOME_LDL_CLEARANCE : 9.58e−14
WP_COMPOSITION_OF_LIPID_PARTICLES : 1.66e−12
BIOCARTA_LDL_PATHWAY : 2.58e−12
REACTOME_VLDLR_INTERNALISATION... : 6.15e−11
BIOCARTA_S1P_PATHWAY : 8.51e−11
WP_SREBF_AND_MIR33_IN_CHOLESTE... : 6.86e−07
Bonferroni p−value == 0.05
- log10 (p-value)
Database pathways (5 – 50 genes)
neutral: MAF < 0.001
Bonferroni p−value == 0.05
- log10 (p-value)
Database pathways (5 – 50 genes)
damaging: MAF < 0.001
LDL_core : 5.48e−48
WP_METABOLIC_PATHWAY_OF_LDL_HD... : 1.36e−35
REACTOME_VLDLR_INTERNALISATION... : 2.98e−30
REACTOME_LDL_CLEARANCE : 3.79e−27
BIOCARTA_FXR_PATHWAY : 4.74e−16
WP_COMPOSITION_OF_LIPID_PARTICLES : 5.34e−15
BIOCARTA_LDL_PATHWAY : 5.36e−14
REACTOME_CHYLOMICRON_CLEARANCE : 3.78e−14
BIOCARTA_S1P_PATHWAY : 3.32e−11
WP_SREBF_AND_MIR33_IN_CHOLESTE... : 7.61e−09
Bonferroni p−value == 0.05
- log10 (p-value)
Database pathways (5 – 50 genes)
synonymous: MAF < 0.001
Bonferroni p−value == 0.05
- log10 (p-value)
Database pathways (5 – 50 genes)

**Supplementary Figure S9. Stratified analysis of rare variant burden in database pathway modules across multiple ancestries in UKBB for the high LDL-C vs low LDL-C phenotype with NERINE.** Pathway Manhattan plots showing Fisher's combined p-values across different ancestries in six functional categories of variants: LoF (i.e., frameshifts, insertions, deletions, and splice variants), damaging missense (i.e., missenses predicted to be damaging by in-silico tools), damaging (i.e., LoF and damaging missenses), missense, neutral (i.e., missenses predicted to be benign by in-silico tools), and synonymous. Applying NERINE across a database of 306 canonical pathways to compare individuals with high LDL cholesterol with the ones with low LDL cholesterol from five major ancestry groups: EUR, AFR, AMR, SAS, and EAS in UKBB, we identified a significant cumulative effect of rare (MAF < 0.001) variants in LoF, damaging missense, damaging, and missense categories in key lipid-related pathways, such as the core module of LDL-related genes. No significant burden of neutral missense or synonymous variants was observed. The horizontal dashed line represents the Bonferroni-corrected combined p-value cutoff of 0.05.

### Figure S10

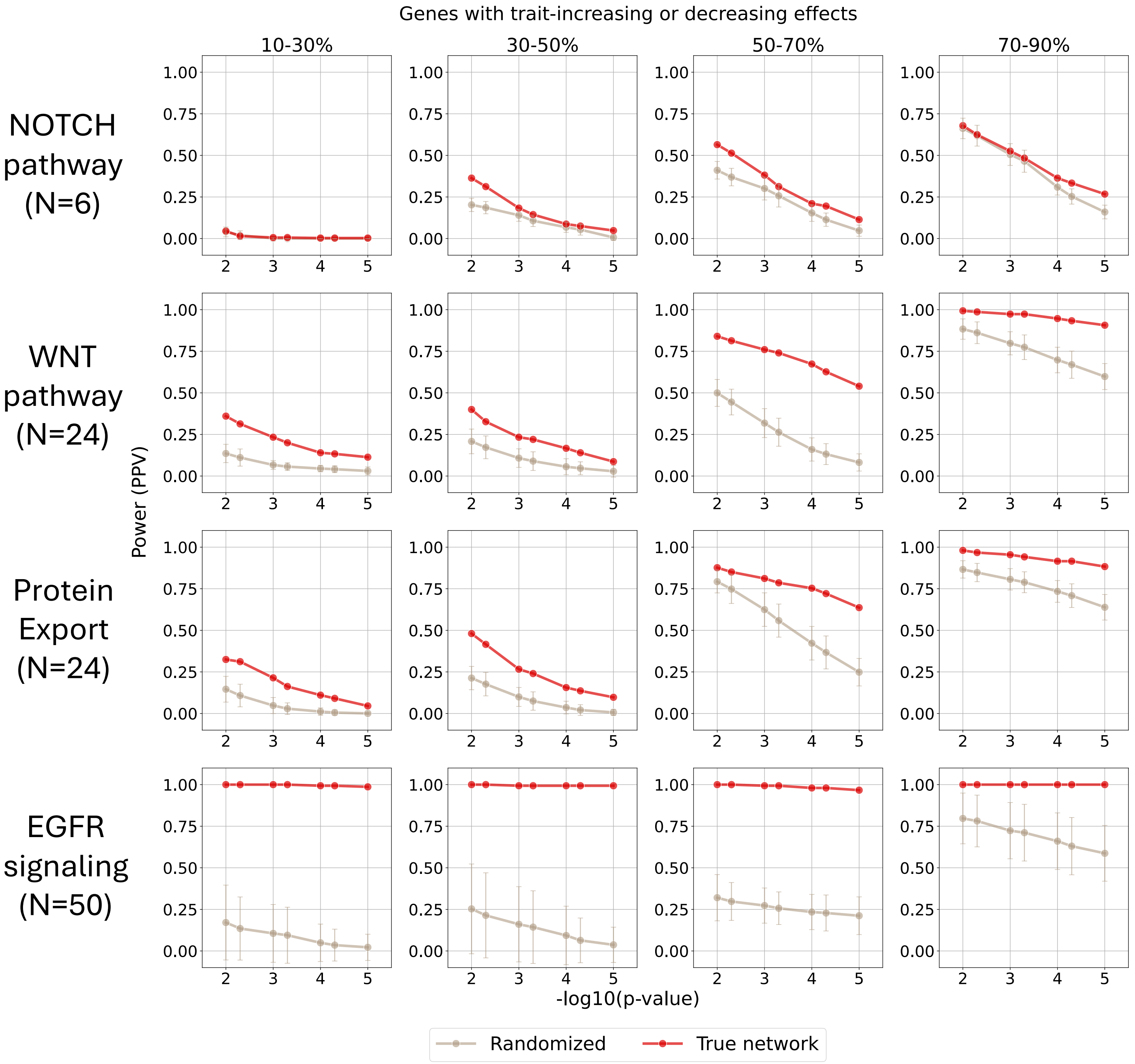

Genes with trait-increasing or decreasing effects
10-30%
30-50%
50-70%
70-90%
NOTCH pathway (N=6)
WNT pathway (N=24)
Protein Export (N=24)
EGFR signaling (N=50)
Power (PPV)
-log10(p-value)
Randomized
True network

**Supplementary Figure S10. NERINE effectively utilizes the true network topology to achieve greater statistical power in simulations with well-studied pathways.** Simulations were performed with different network architectures for well-studied pathways of different sizes: the NOTCH pathway (m=6), the WNT pathway (m=24), the protein export pathway (m=24), and the EGFR signaling pathway (m=50). Pathway gene lists were extracted from MSigDB v7.3, and high-confidence physical and genetic interactions from protein-protein interaction (PPI) databases were used as true network edges between pathway genes (see Methods). For each network gene set, 100 random networks were generated by randomly assigning edges between the genes. The allele counts in cases and controls were generated from independent binomial distributions. The empirical power of the methods was measured for a simulated binary disease trait in a cohort of 2,000 cases and 2,000 controls with different proportions of genes in the network having both trait-increasing and trait-decreasing effects simulated with a network effect of $\theta = 0.1$. From left to right, the power plots show networks with different noise profiles, mimicking situations from having a very noisy network to a highly relevant one. We performed 250 per noise profile per network. Power was calculated as the positive predictive value (PPV) at different p-value cutoffs: 1e-2, 5e-3, 1e-3, 5e-4, 1e-4, 5e-5, and 1e-5. Error bars represent the standard deviation from the mean PPV across different iterations.

### Figure S11

A.

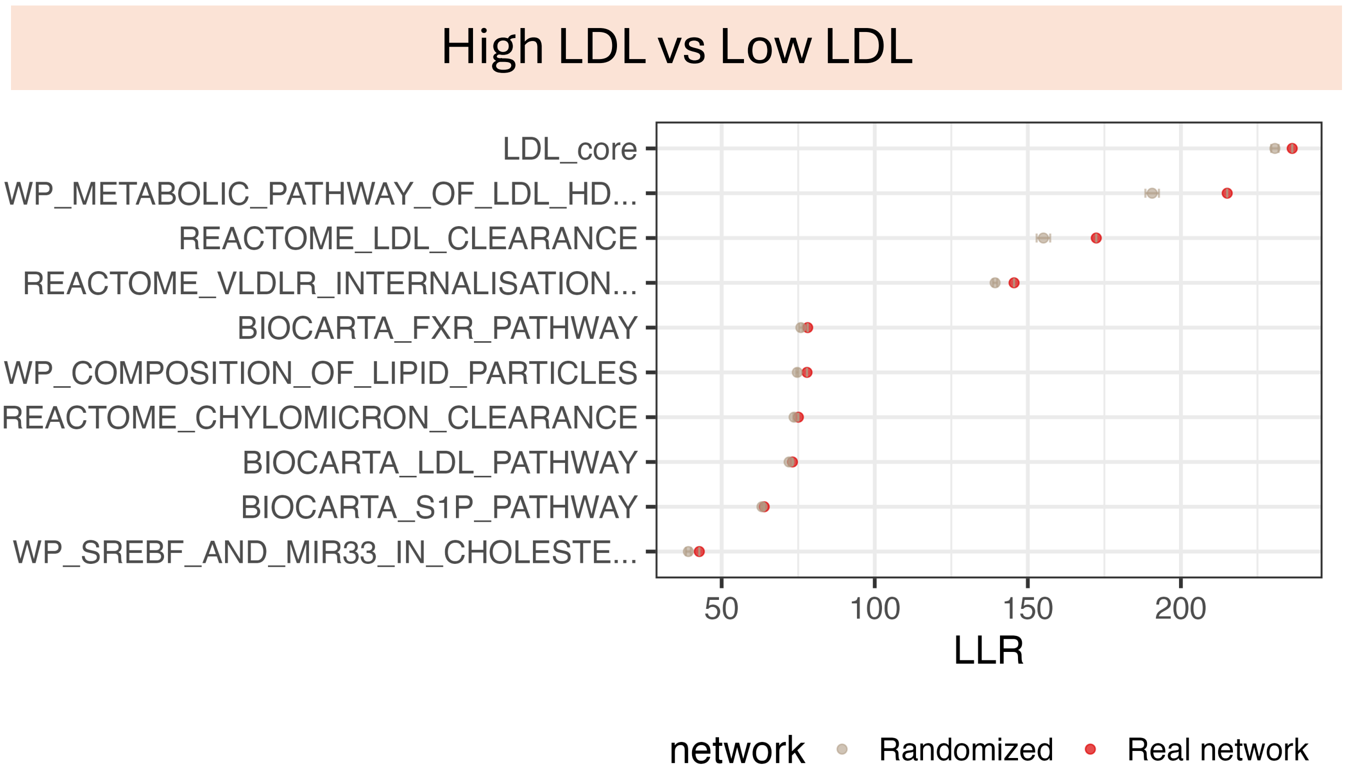


B.

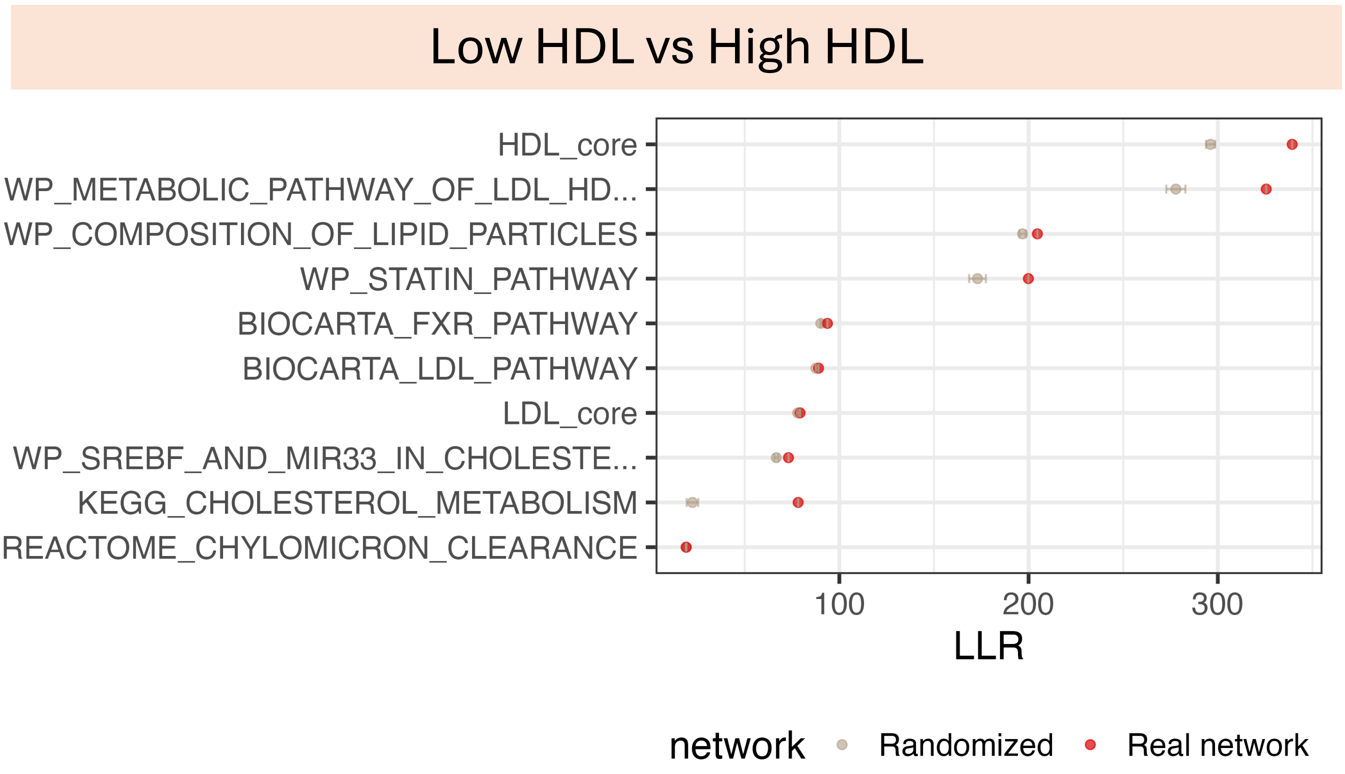

**Supplementary Figure S11. NERINE's performance on real vs. randomized network topologies in binarized LDL-C and HDL-C phenotypes in the UK biobank.** We tested real vs randomized networks in two comparisons in the UKBB cohort **A)** high LDL-C vs low LDL-C, and **B)** low HDL-C vs high HDL-C. We selected the database-wide significant pathways in each case and extracted database edges for the member genes to form "real" network topologies (Methods). For each pathway, 100 randomized networks were created by introducing random edges among the member genes. NERINE was applied on both real and randomized network topologies for these phenotypes. For each network in both phenotypes, NERINE achieves a higher log-likelihood ratio (LLR) and lower p-value with real topology than with random edges. Here, the error bars indicate standard error around LLR over 100 random networks.

### Figure S12

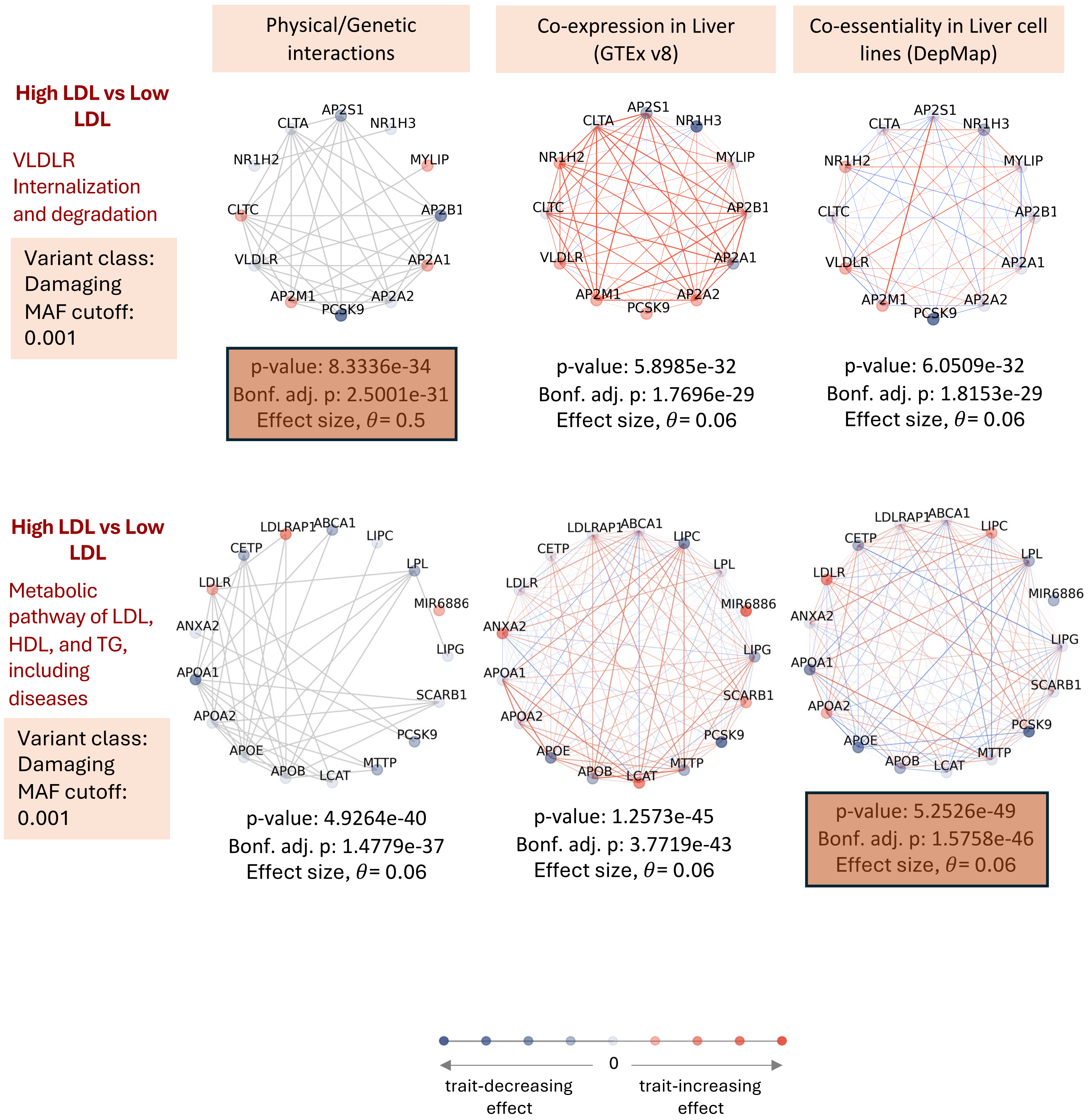

Physical/Genetic interactions
Co-expression in Liver (GTEx v8)
Co-essentiality in Liver cell lines (DepMap)
High LDL vs Low LDL
VLDLR Internalization and degradation
Variant class: Damaging
MAF cutoff: 0.001
AP2S1
CLTA
NR1H3
NR1H2
MYLIP
CLTC
AP2B1
VLDLR
AP2A1
AP2M1
AP2A2
PCSK9
p-value: 8.3336e-34
Bonf. adj. p: 2.5001e-31
Effect size, θ= 0.5
p-value: 5.8985e-32
Bonf. adj. p: 1.7696e-29
Effect size, θ= 0.06
p-value: 6.0509e-32
Bonf. adj. p: 1.8153e-29
Effect size, θ= 0.06
High LDL vs Low LDL
Metabolic pathway of LDL, HDL, and TG, including diseases
Variant class: Damaging
MAF cutoff: 0.001
LDLRAP1
ABCA1
LIPC
CETP
LPL
LDLR
MIR6886
ANXA2
LIPG
APOA1
SCARB1
APOA2
PCSK9
APOE
MTTP
APOB
LCAT
p-value: 4.9264e-40
Bonf. adj. p: 1.4779e-37
Effect size, θ= 0.06
p-value: 1.2573e-45
Bonf. adj. p: 3.7719e-43
Effect size, θ= 0.06
p-value: 5.2526e-49
Bonf. adj. p: 1.5758e-46
Effect size, θ= 0.06
0
trait-decreasing effect
trait-increasing effect

**Supplementary Figure S12. Example scenarios where NERINE selects physical and genetic interactions and gene-gene coessentiality relationships as the most informative source of topology.** To describe the edge relationship of selected genes, we used three data sources: high-confidence physical or genetic interactions from protein-protein interaction (PPI) databases, co-expression in liver tissue from GTEx v8, coessentiality from DepMap (2023Q2). The top row shows the enrichment of rare damaging variants in the VLDLR Internalization and degradation pathway in the hyperlipidemia phenotype (high LDL vs low LDL) in UKBB. In this case, NERINE achieved the most significant p-value with high-confidence physical and genetic interactions from PPI databases. The bottom row shows the enrichment of rare damaging variants in the metabolic pathway of LDL, HDL, and TG, including diseases from the Wikipathways database in hyperlipidemia in UKBB. The coessentiality of genes across liver cell lines in DepMap best describes the relationship of these genes, enabling NERINE to achieve the most significant p-value.

### Figure S13

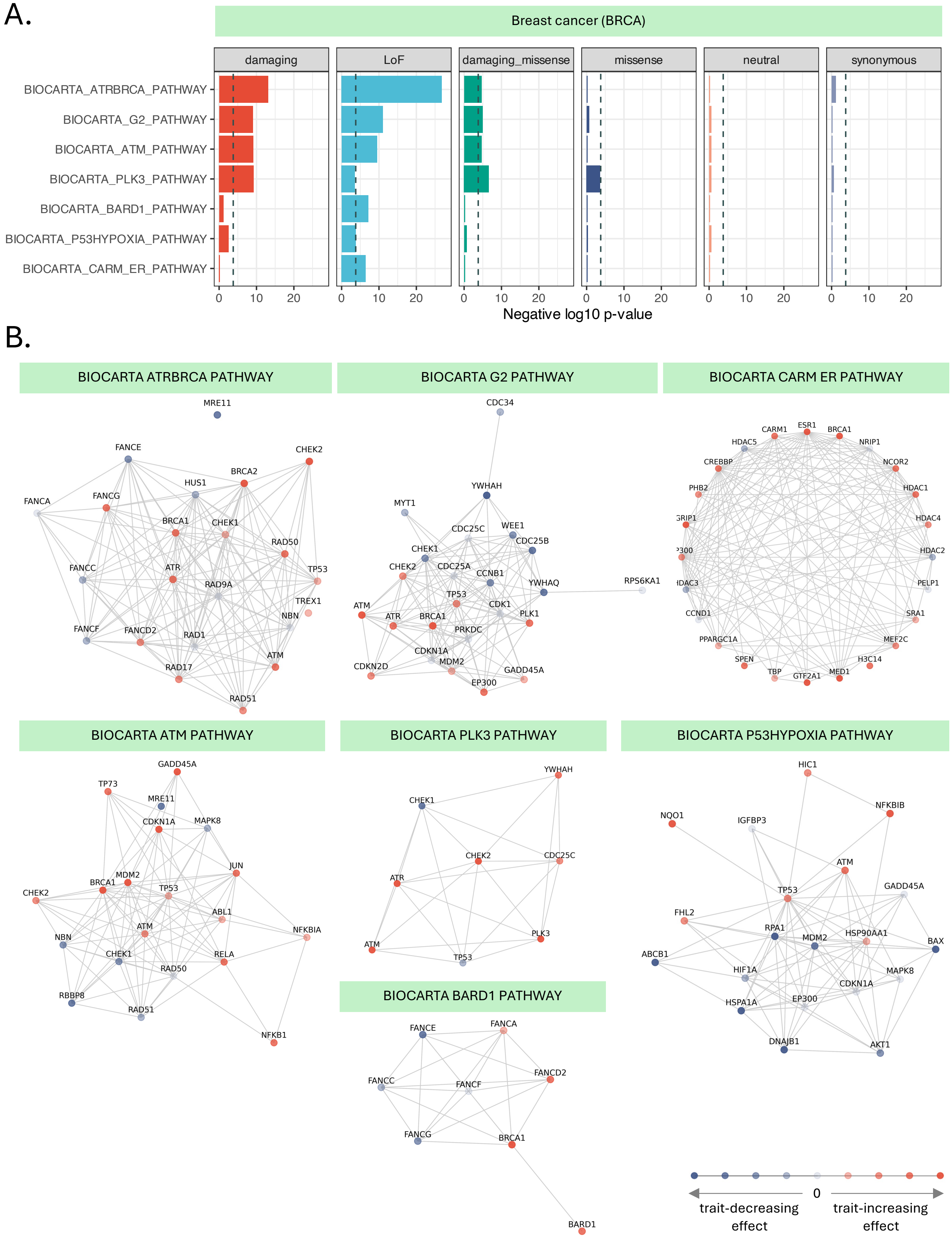

A.
Breast cancer (BRCA)
damaging
LoF
damaging_missense
missense
neutral
synonymous
BIOCARTA_ATRBRCA_PATHWAY
BIOCARTA_G2_PATHWAY
BIOCARTA_ATM_PATHWAY
BIOCARTA_PLK3_PATHWAY
BIOCARTA_BARD1_PATHWAY
BIOCARTA_P53HYPOXIA_PATHWAY
BIOCARTA_CARM_ER_PATHWAY
0 10 20
Negative log10 p-value
B.
BIOCARTA ATRBRCA PATHWAY
MRE11
FANCE
CHEK2
BRCA2
HUS1
FANCA
FANCG
BRCA1
CHEK1
RAD50
FANCC
ATR
TP53
RAD9A
TREX1
NBN
FANCF
FANCD2
RAD1
ATM
RAD17
RAD51
BIOCARTA G2 PATHWAY
CDC34
YWHAH
MYT1
CDC25C
WEE1
CDC25B
CHEK1
CHEK2
CDC25A
CCNB1
YWHAQ
RPS6KA1
TP53
ATM
CDK1
ATR
BRCA1
PLK1
PRKDC
CDKN1A
MDM2
CDKN2D
GADD45A
EP300
BIOCARTA CARM ER PATHWAY
CARM1
ESR1
BRCA1
HDAC5
NRIP1
CREBBP
NCOR2
PHB2
HDAC1
HDAC4
HDAC2
PELP1
CCND1
SRA1
PPARGC1A
MEF2C
SPEN
H3C14
TBP
GTF2A1
MED1
BIOCARTA ATM PATHWAY
GADD45A
TP73
MRE11
CDKN1A
MAPK8
JUN
MDM2
BRCA1
CHEK2
TP53
ABL1
ATM
NBN
NFKBIA
CHEK1
RELA
RAD50
RBBP8
RAD51
NFKB1
BIOCARTA PLK3 PATHWAY
YWHAH
CHEK1
CHEK2
CDC25C
ATR
PLK3
ATM
TP53
BIOCARTA BARD1 PATHWAY
FANCE
FANCA
FANCD2
FANCC
FANCF
FANCG
BRCA1
BARD1
BIOCARTA P53HYPOXIA PATHWAY
HIC1
NFKBIB
NQO1
IGFBP3
ATM
GADD45A
TP53
FHL2
RPA1
MDM2
HSP90AA1
BAX
ABCB1
HIF1A
MAPK8
CDKN1A
HSPA1A
EP300
DNAJB1
AKT1
0
trait-decreasing effect
trait-increasing effect

**Supplementary Figure S13. NERINE identifies significant rare variant burden in seven pathway gene modules in breast cancer (BRCA) in the UK and MGB biobanks. A.** Bonferroni-significant findings for the BRCA phenotype across a database of 306 pathways in six functional categories of rare variants—(i) LoF, (ii) damaging missense, (iii) damaging, (iv) missense, (v) neutral missense, and (vi) synonymous are shown. No inflation was observed in neutral missense or synonymous variant categories. For each variant category, pathways were tested individually in UKBB and MGBBB cohorts, and p-values were meta-analyzed using Fisher's combined test. Figure shows negative log-transformed Fisher's combined p-values for significant pathways. The dashed grey line represents the Bonferroni threshold of 0.05. **B.** Network topologies for the pathways with significant rare LoF variant-burden in BRCA are shown; nodes are colored according to individual gene effects, averaged across biobanks. Trait-increasing effects are represented by shades of orange, and trait-decreasing effects are represented by different shades of purple, as shown on the scale. A darker color means a more pronounced effect.

### Figure S14

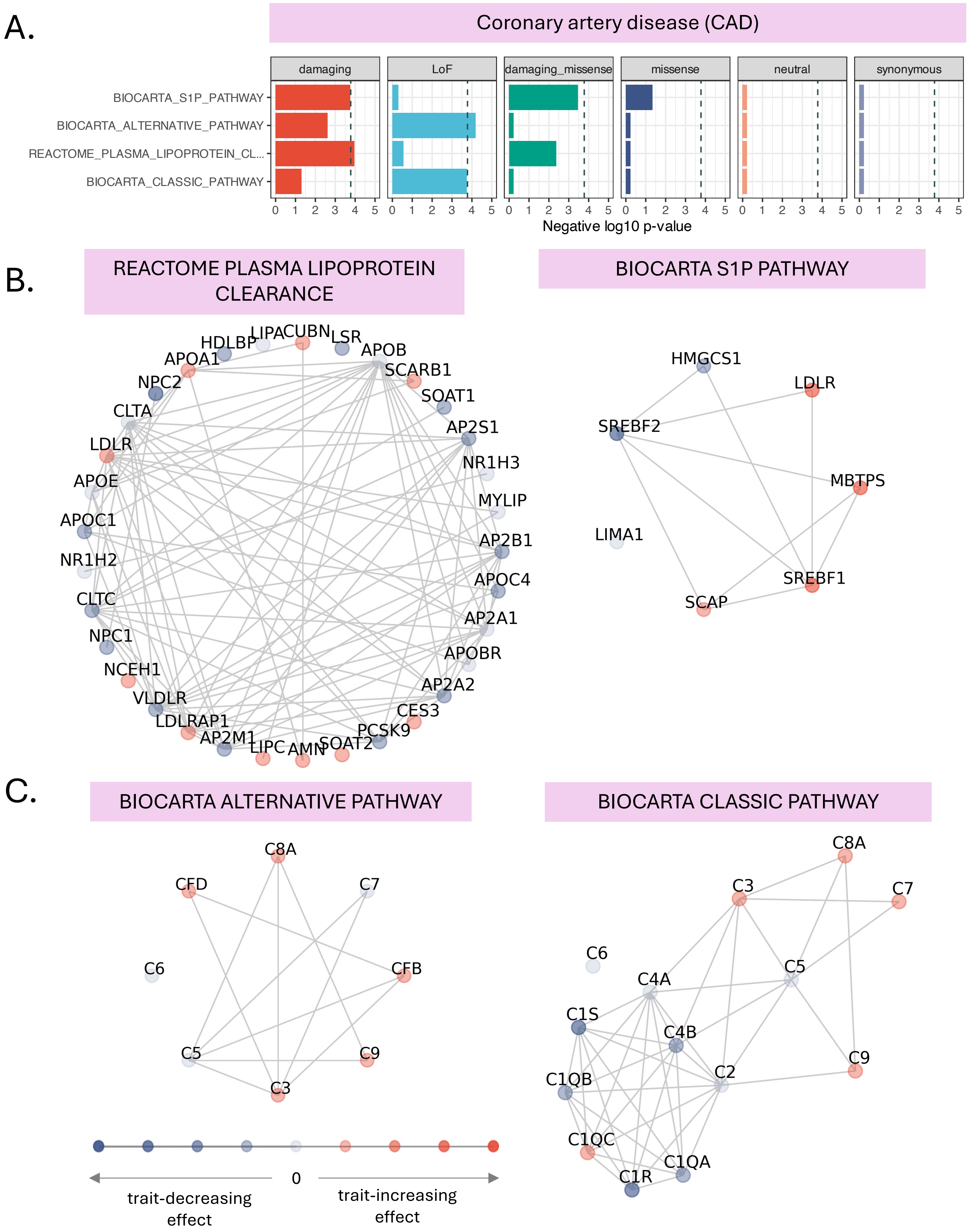

A.
Coronary artery disease (CAD)
damaging
LoF
damaging_missense
missense
neutral
synonymous
BIOCARTA_S1P_PATHWAY
BIOCARTA_ALTERNATIVE_PATHWAY
REACTOME_PLASMA_LIPOPROTEIN_CL...
BIOCARTA_CLASSIC_PATHWAY
0 1 2 3 4 5
Negative log10 p-value
B.
REACTOME PLASMA LIPOPROTEIN CLEARANCE
HDLBP
LIPA
CUBN
LSR
APOB
APOA1
SCARB1
NPC2
SOAT1
CLTA
AP2S1
LDLR
NR1H3
APOE
MYLIP
APOC1
AP2B1
NR1H2
APOC4
CLTC
AP2A1
NPC1
APOBR
NCEH1
AP2A2
VLDLR
CES3
LDLRAP1
PCSK9
AP2M1
LIPC
AMN
SOAT2
BIOCARTA S1P PATHWAY
HMGCS1
LDLR
SREBF2
MBTPS
LIMA1
SREBF1
SCAP
C.
BIOCARTA ALTERNATIVE PATHWAY
C8A
CFD
C7
C6
CFB
C5
C9
C3
0
trait-decreasing effect
trait-increasing effect
BIOCARTA CLASSIC PATHWAY
C8A
C3
C7
C6
C5
C4A
C1S
C4B
C9
C2
C1QB
C1QC
C1QA
C1R

**Supplementary Figure S14. NERINE identifies significant rare variant burden in four pathway gene modules in coronary artery disease (CAD) in the UK and MGB biobanks. A.** Bonferroni-significant findings for the CAD phenotype across a database of 306 pathways in six functional categories of rare variants—(i) LoF, (ii) damaging missense, (iii) damaging, (iv) missense, (v) neutral missense, and (vi) synonymous are shown. No inflation was observed in neutral missense or synonymous variant categories. Pathways were tested individually in UKBB and MGBBB cohorts for each variant category, and p-values were meta-analyzed using Fisher's combined test. The figure shows negative log-transformed Fisher's combined p-values for significant pathways. The dashed grey line represents the Bonferroni threshold of 0.05. **B.** Network topologies for the pathways with significant rare damaging variant-burden in CAD are shown with individual gene effects. **C.** Network topologies for the pathways with significant rare LoF variant burden in CAD are shown with individual gene effects. Trait-increasing effects are represented by shades of orange, and trait-decreasing effects are represented by shades of blue, as shown on the scale. A darker color means a more pronounced effect.

### Figure S15

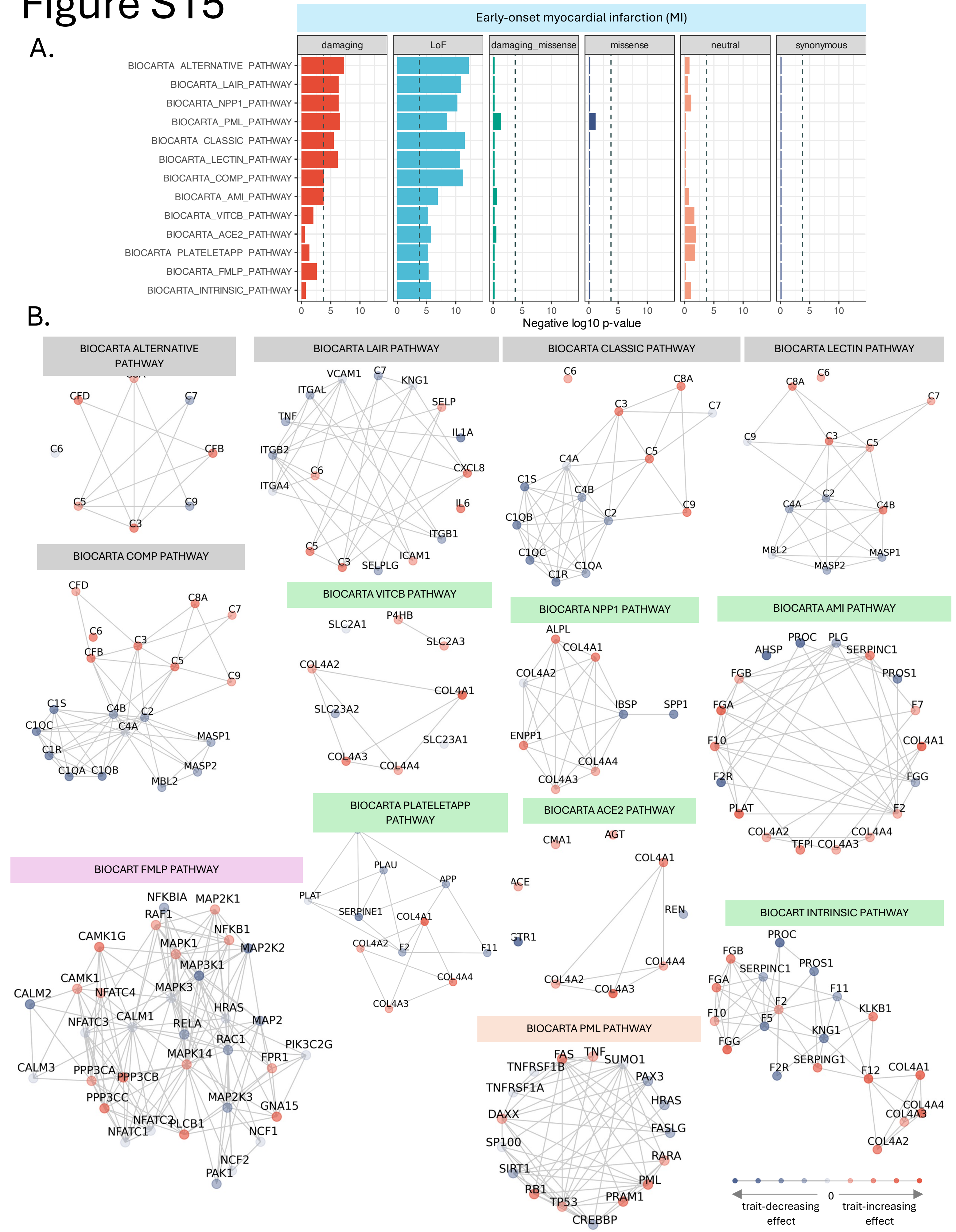
A.
Early-onset myocardial infarction (MI)
damaging
LoF
damaging_missense
missense
neutral
synonymous
BIOCARTA_ALTERNATIVE_PATHWAY
BIOCARTA_LAIR_PATHWAY
BIOCARTA_NPP1_PATHWAY
BIOCARTA_PML_PATHWAY
BIOCARTA_CLASSIC_PATHWAY
BIOCARTA_LECTIN_PATHWAY
BIOCARTA_COMP_PATHWAY
BIOCARTA_AMI_PATHWAY
BIOCARTA_VITCB_PATHWAY
BIOCARTA_ACE2_PATHWAY
BIOCARTA_PLATELETAPP_PATHWAY
BIOCARTA_FMLP_PATHWAY
BIOCARTA_INTRINSIC_PATHWAY
0 5 10
Negative log10 p-value
B.
BIOCARTA ALTERNATIVE PATHWAY
BIOCARTA LAIR PATHWAY
BIOCARTA CLASSIC PATHWAY
BIOCARTA LECTIN PATHWAY
BIOCARTA COMP PATHWAY
BIOCARTA VITCB PATHWAY
BIOCARTA NPP1 PATHWAY
BIOCARTA AMI PATHWAY
BIOCARTA PLATELETAPP PATHWAY
BIOCARTA ACE2 PATHWAY
BIOCART FMLP PATHWAY
BIOCART INTRINSIC PATHWAY
BIOCARTA PML PATHWAY
0
trait-decreasing effect
trait-increasing effect

**Supplementary Figure S15. NERINE identifies significant rare variant burden in thirteen pathway gene modules in early-onset myocardial infarction (MI) in the UK and MGB biobanks. A.** Bonferroni-significant findings for the CAD phenotype across a database of 306 pathways in six functional categories of rare variants—(i) LoF, (ii) damaging missense, (iii) damaging, (iv) missense, (v) neutral missense, and (vi) synonymous are shown. No inflation was observed in neutral missense or synonymous variant categories. For each variant category, pathways were tested individually in UKBB and MGBBB cohorts, and p-values were meta-analyzed using Fisher's combined test. The figure shows negative log-transformed Fisher's combined p-values for significant pathways. The dashed line represents the Bonferroni threshold of 0.05. **B.** Network topologies for the pathways with significant rare LoF variant burden in MI are shown with individual gene effects. Trait-increasing effects are represented by shades of orange, and trait-decreasing effects are represented by different shades of purple as shown on the scale. A darker color means a more pronounced effect. Pathways are color-coded by the broad groups they belong to. Here, light grey: inflammatory response, light green: extracellular matrix proteins and coagulation, light orange: regulation of transcriptional activity, and light purple: MAPK signaling cascade. We recognize several caveats: the findings around complement system pathways showed disparity among the UKBB and MGBBB. Also, the finding around the MAPK signaling pathway might be due to either true biology or the CHIP (Clonal Hematopoiesis of Indeterminate Potential) effect because the original bio-samples were primarily from blood in MGBBB.

### Figure S16

A.

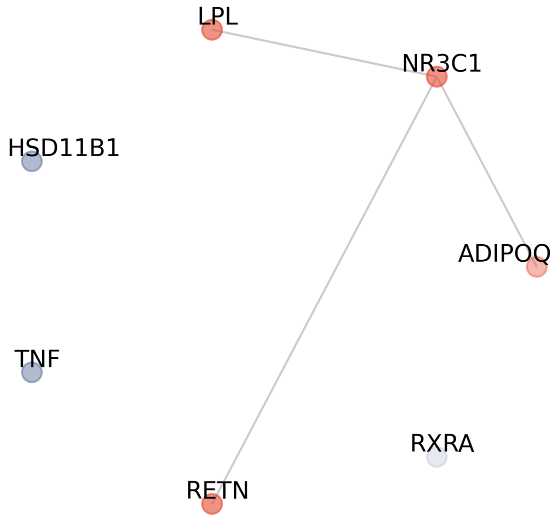


**Adipogenesis without PPARG**
Fisher's combined p = **1.9964e-6**

B.

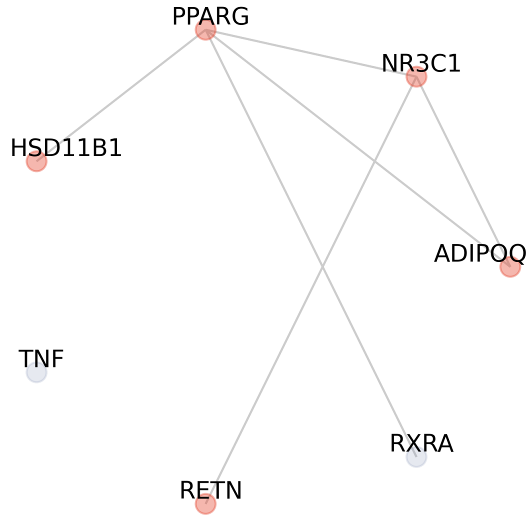


**Adipogenesis without LPL**
Fisher's combined p = **1.3008e-02**

C.

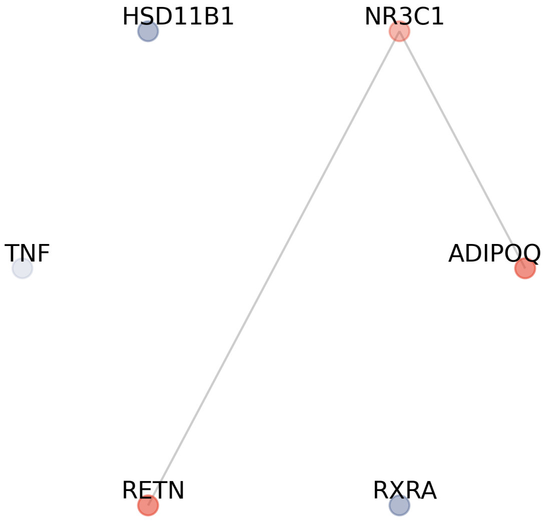


**Adipogenesis without LPL and PPARG**
Fisher's combined p = **1.5571e-02**

**Supplementary Figure S16. Sensitivity analysis showing NERINE's performance on the *adipogenesis* (BIOCARTA VOBESITY PATHWAY) network for the T2D phenotype in UKBB after removing the effects of *LPL and PPARG*. A.** NERINE identifies a significant rare damaging variant burden even after removing *PPARG* and its connections from the network. **B.** After removing *LPL* and its connections, NERINE identified a nominally significant burden across the rest of the network. **C.** Removing both *LPL and PPARG* from the network, NERINE still identifies a nominally significant burden of rare damaging variant burden. Neither *LPL* nor *PPARG* by themselves showed significant genome-wide rare variant burden in single-gene analyses of the UKBB (Genebass[33] and SAIGE-GENE+[19]).

### Figure S17

A.

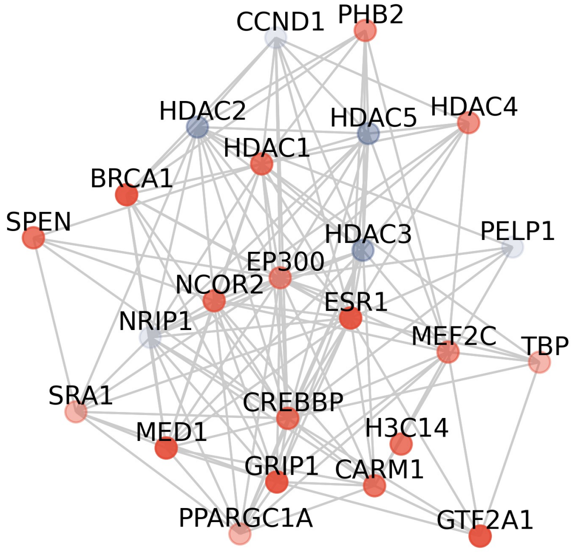


**Estrogen receptor pathway**
NERINE p-value: **5.78e-6**
variant category: LoF

B.

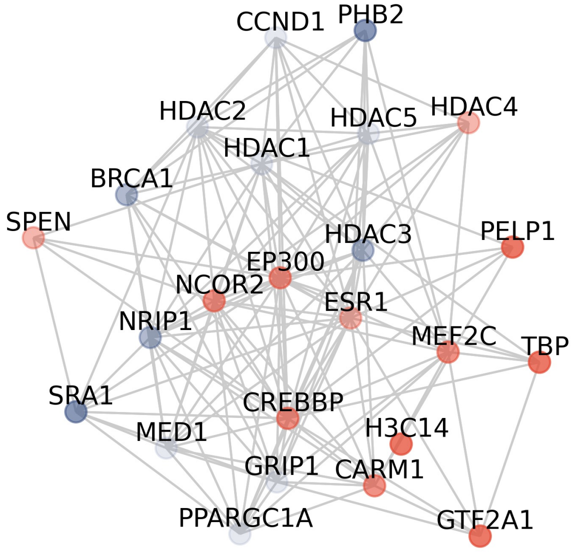


**Estrogen receptor pathway without *BRCA1* mutations**
NERINE p-value: **3.43e-2**
variant category: LoF

C.

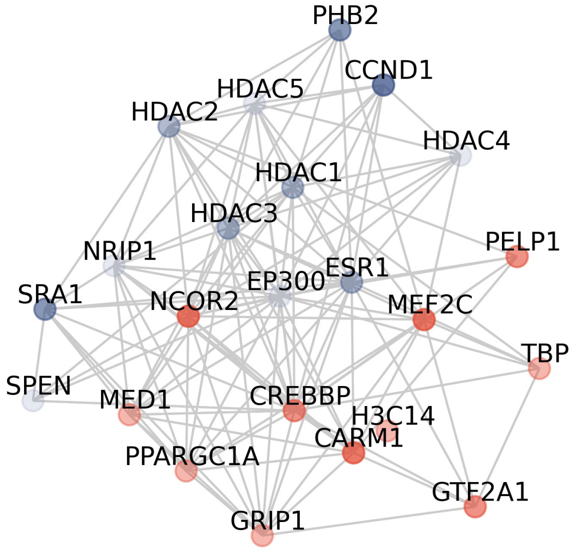


**Estrogen receptor pathway without *BRCA1* and its connections**
NERINE p-value: 6.61e-2
variant category: LoF

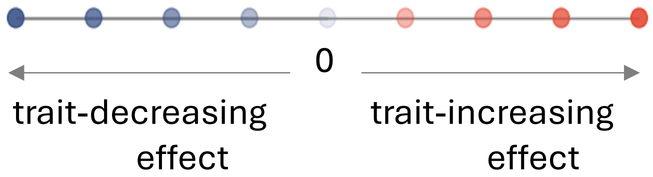

**Supplementary Figure S17. Sensitivity analysis shows NERINE's performance on the *regulation of the estrogen receptor* network for the BRCA phenotype in UKBB after removing the effect of *BRCA1*. A.** NERINE identifies a database-wide significant rare LoF variant burden in the original analysis of the *regulation of the estrogen receptor* network. **B.** After removing the observed mutation counts in *BRCA1* but keeping the gene and its connections in the network, NERINE identified a nominally significant burden. **C.** NERINE was run after removing the *BRCA1* gene and its edges from the network, and a suggestive rare LoF variant burden was still identified.

### Figure S18

A.

#### Running NERINE on PD GWAS gene modules

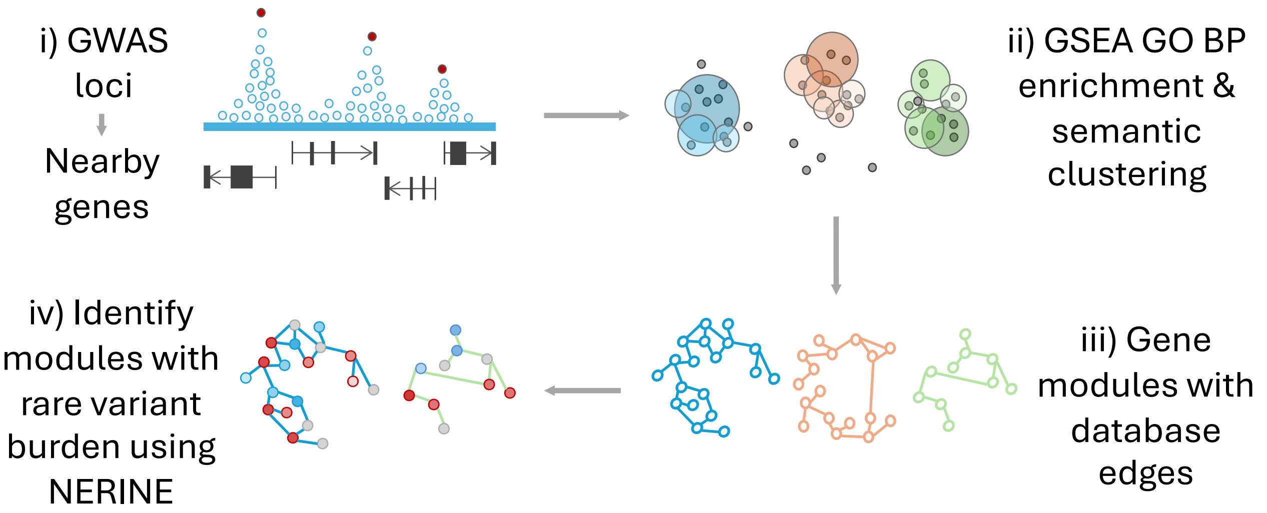


B.

#### GWAS gene module with significant rare variant burden

**peptidyl-threonine modification**

| Category | Avg. $\hat{\theta}$ | Fisher combo-p | Screen-wide Bonf. p |
|---|---|---|---|
| LoF | **0.9** | **7.23E-03** | **0.0434** |
| Neutral | 0 | 5.97E-01 | 1 |
| Synonymous | 0 | 5.97E-01 | 1 |

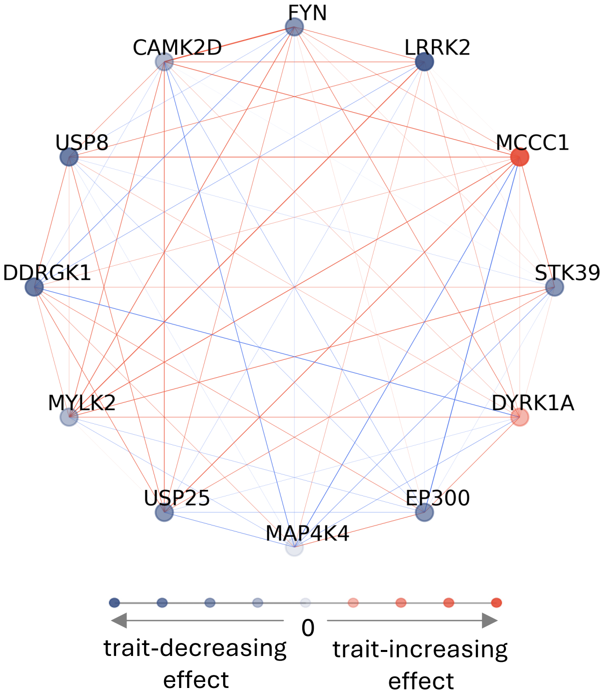


| Gene symbol | Known rare variant hit | Role relevant to PD |
|---|---|---|
| CAMK2D | X | Synaptic plasticity |
| DDRGK1 | X | Protein homeostasis, inflammation, stress response |
| DYRK1A | X | Tau phosphorylation, neurodevelopment |
| EP300 | X | Histone acetylation, neuronal survival |
| FYN | X | Tau phosphorylation, synaptic function |
| LRRK2 | ✓ | Key risk gene for PD, kinase activity, neuroinflammation |
| MAP4K4 | X | Neuroinflammation, stress response, apoptosis |
| MCCC1 | X | Mitochondrial metabolism |
| MYLK2 | X | Muscle and neuronal cytoskeletal stability |
| STK39 | X | Oxidative stress response, neuroinflammation |
| USP25 | X | Inflammation, ubiquitin signaling, proteostasis |
| USP8 | X | α-synuclein degradation, endosomal trafficking, LRRK2 regulation |

**Supplementary Figure S18. NERINE identifies significant rare LoF variant burden in a *peptidyl-threonine modification* network enriched in PD GWAS genes. A.** Schematic diagram showing the process of network hypotheses generation for interrogating gene modules enriched in PD GWAS genes using NERINE. Among GWAS genes, we identified six GO biological process (BP) modules which were tested with NERINE on both AMP-PD and UKBB sporadic PD vs. control cohorts. **B.** GWAS genes enriched in the GO biological process module related to *Peptidyl-threonine modification* show significant rare LoF variant (i.e., frameshifts, insertions, deletions, and splice variants) burden in sporadic PD cases compared to controls in two independent datasets: AMP-PD and UK Biobank (UKBB). Co-essentiality in CNS cell-types provided the most informative edge relationships for this module. Screen-wide significance was determined by applying Bonferroni correction over Fisher combined p-values. The absence of enrichment of neutral missense and synonymous variants in the networks served as an internal control. In each scenario, trait-increasing effects are represented by shades of orange and trait-decreasing effects are represented by shades of purple, as shown on the scale. A darker color means a more pronounced effect in each direction. For the coessentiality network in **B**, color of the edge indicates the sign of the correlation (red: positive; blue: negative) and the thickness of the edge corresponds to the strength of the correlation. We extracted information about member genes based on germline genetics from the GWAS catalog (https://www.ebi.ac.uk/gwas/) and the Genebass[33], SAIGE-GENE+[19], and PD-specific rare variant studies[26,27]. Annotations indicating known roles of genes relevant to PD and neurodegeneration were derived from SynGO (https://www.syngoportal.org/).

### Figure S19

A.

#### DA neuron essentiality network

***HMGB1-USP10* regulation of autophagy module**

**Category:** damaging missense
Bonf. p: **1.86E-02**

**Category:** damaging
Bonf. p: **3.11E-02**

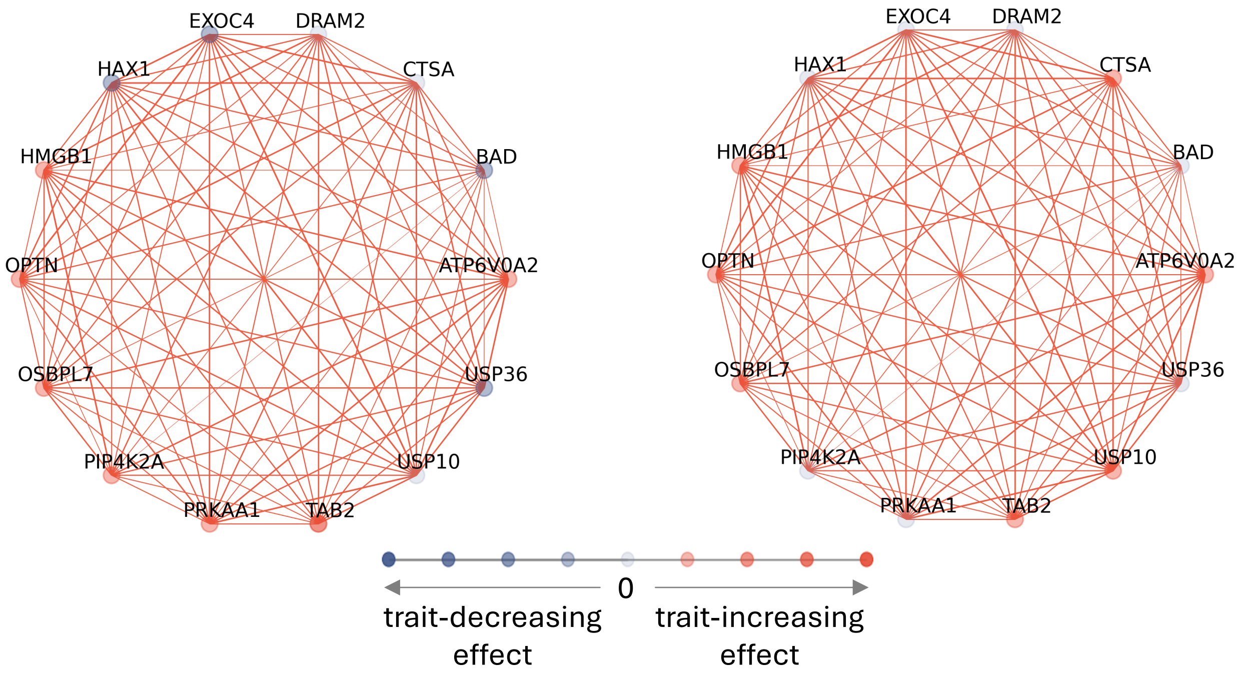


B.

| Gene symbol | GWAS hit | Known rare variant hit | Role relevant to PD |
|---|---|---|---|
| ATP6V0A2 | X | X | Lysosomal acidification, autophagy |
| BAD | X | X | Apoptosis regulation, neuronal survival |
| CTSA | X | X | Lysosomal function, proteostasis |
| DRAM2 | X | X | Autophagy regulation, oxidative stress response |
| EXOC4 | X | X | Synaptic vesicle trafficking |
| HAX1 | X | X | Mitochondrial integrity, cell survival |
| HMGB1 | X | X | Neuroinflammation, immune response |
| OPTN | X | X | Mitophagy, clearance of damaged mitochondria |
| OSBPL7 | X | X | Lipid metabolism, neuronal membrane homeostasis |
| PIP4K2A | X | X | Membrane signaling, autophagy regulation |
| PRKAA1 | X | X | Energy metabolism, neuronal stress response |
| TAB2 | X | X | NF-κB signaling, neuroinflammation |
| USP10 | X | X | Ubiquitin signaling, protein degradation |
| USP36 | X | X | Nucleolar protein homeostasis |

**Supplementary Figure S19. Additional gene-level information on NERINE's screen-wide significant findings in DA neuron essentiality screen. A.** In the DA neuron essentiality screen, NERINE identified screen-wide significant burdens of rare damaging missense (left) and damaging (right) variants in the GO biological process module related to the *regulation of autophagy*. Coexpression in the midbrain *substantia nigra* region provided the most informative topology for this module. Screen-wide significance of networks was determined by applying Bonferroni correction over the Fisher combined p-values using the number of gene modules tested in the screen ($t_{eff}$ = 10). Here, trait-increasing effects are represented by shades of orange and trait-decreasing effects are represented by shades of purple, as shown on the scale. A darker color means a more pronounced effect in each direction. For the coexpression network in **A**, the color of the edge indicates the sign of the correlation (red: positive; blue: negative) and the thickness of the edge corresponds to the correlation strength. **B.** Additional information on DA essentiality genes in the *regulation of autophagy* module from previous studies. We extracted information about member genes based on germline genetics from the GWAS catalog (https://www.ebi.ac.uk/gwas/) and the Genebass[33], SAIGE-GENE+[19], and PD-specific rare variant studies[26,27]. Annotations indicating known roles of genes relevant to PD and neurodegeneration were derived from SynGO (https://www.syngoportal.org/).

### Figure S20

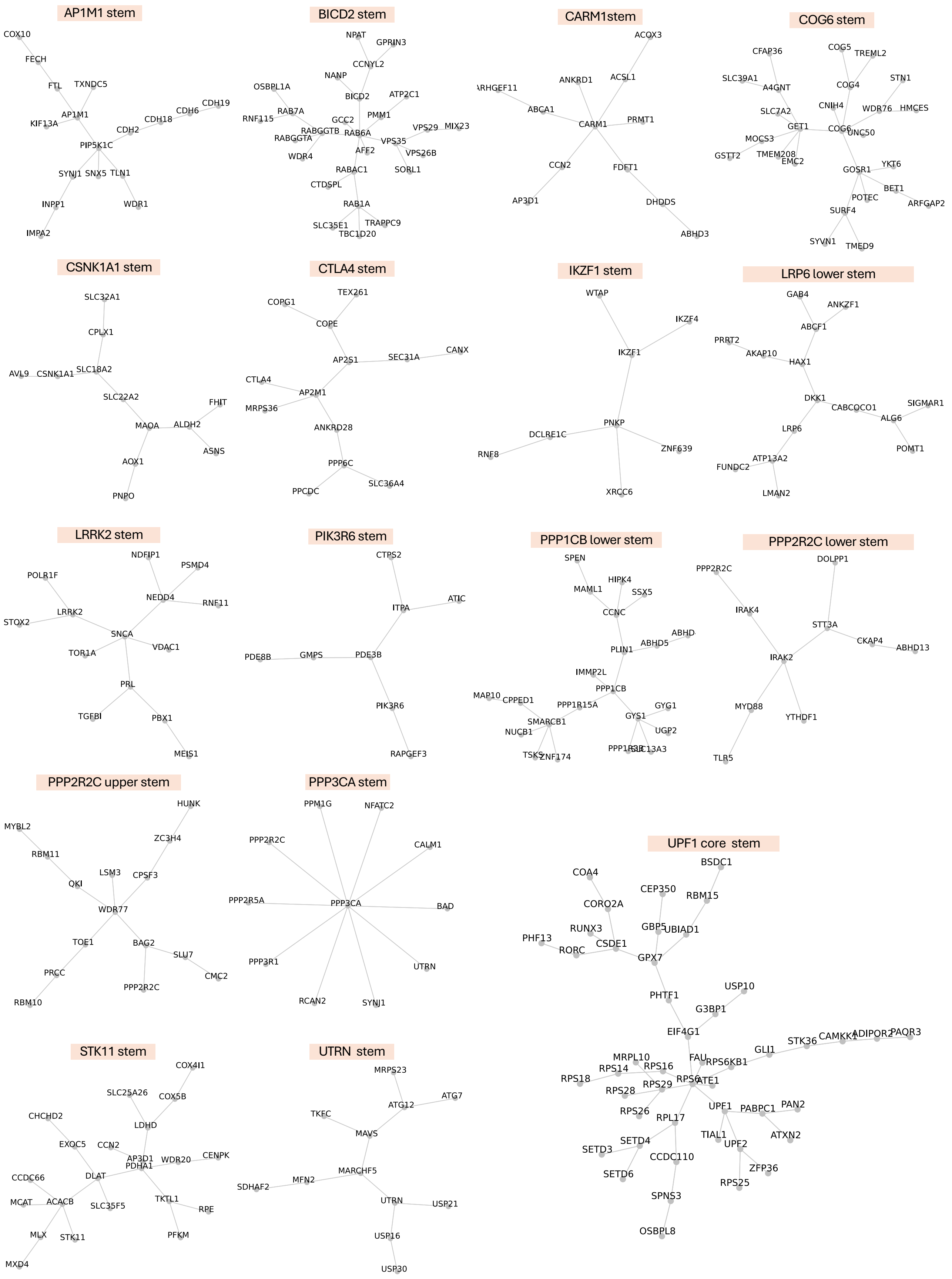

AP1M1 stem
COX10
FECH
FTL
TXNDC5
AP1M1
KIF13A
CDH19
CDH6
CDH18
CDH2
PIP5K1C
SYNJ1
SNX5
TLN1
INPP1
WDR1
IMPA2
BICD2 stem
NPAT
GPRIN3
CCNYL2
NANP
OSBPL1A
BICD2
ATP2C1
RAB7A
RNF115
GCC2
PMM1
RABGGTB
RAB6A
VPS29
MIX23
RABGGTA
VPS35
WDR4
AFF2
VPS26B
RABAC1
SORL1
CTDSPL
RAB1A
SLC35E1
TRAPPC9
TBC1D20
CARM1stem
ACOX3
ANKRD1
ACSL1
ARHGEF11
ABCA1
CARM1
PRMT1
CCN2
FDFT1
AP3D1
DHDDS
ABHD3
COG6 stem
CFAP36
COG5
TREML2
SLC39A1
A4GNT
COG4
STN1
CNIH4
WDR76
HMCES
SLC7A2
GET1
COG6
UNC50
MOCS3
GSTT2
TMEM208
EMC2
YKT6
GOSR1
BET1
POTEC
ARFGAP2
SURF4
SYVN1
TMED9
CSNK1A1 stem
SLC32A1
CPLX1
AVL9
CSNK1A1
SLC18A2
SLC22A2
FHIT
MAOA
ALDH2
ASNS
AOX1
PNPO
CTLA4 stem
COPG1
TEX261
COPE
AP2S1
SEC31A
CANX
CTLA4
AP2M1
MRPS36
ANKRD28
PPP6C
PPCDC
SLC36A4
IKZF1 stem
WTAP
IKZF4
IKZF1
PNKP
DCLRE1C
RNF8
ZNF639
XRCC6
LRP6 lower stem
GAB4
ANKZF1
ABCF1
PRRT2
AKAP10
HAX1
DKK1
CABCOCO1
SIGMAR1
ALG6
LRP6
POMT1
ATP13A2
FUNDC2
LMAN2
LRRK2 stem
NDFIP1
PSMD4
POLR1F
NEDD4
RNF11
LRRK2
STOX2
SNCA
TOR1A
VDAC1
PRL
TGFBI
PBX1
MEIS1
PIK3R6 stem
CTPS2
ITPA
ATIC
PDE8B
GMPS
PDE3B
PIK3R6
RAPGEF3
PPP1CB lower stem
SPEN
HIPK4
MAML1
SSX5
CCNC
ABHD
ABHD5
PLIN1
IMMP2L
PPP1CB
MAP10
CPPED1
PPP1R15A
GYG1
SMARCB1
GYS1
NUCB1
UGP2
TSKS
ZNF174
PPP1R3B
SLC13A3
PPP2R2C lower stem
DOLPP1
PPP2R2C
IRAK4
STT3A
CKAP4
ABHD13
IRAK2
MYD88
YTHDF1
TLR5
PPP2R2C upper stem
HUNK
MYBL2
ZC3H4
RBM11
LSM3
CPSF3
QKI
WDR77
TOE1
BAG2
SLU7
PRCC
CMC2
PPP2R2C
RBM10
PPP3CA stem
PPM1G
NFATC2
PPP2R2C
CALM1
PPP2R5A
PPP3CA
BAD
PPP3R1
UTRN
RCAN2
SYNJ1
UPF1 core stem
BSDC1
COA4
CEP350
RBM15
CORO2A
RUNX3
GBP5
UBIAD1
PHF13
CSDE1
RORC
GPX7
PHTF1
USP10
G3BP1
EIF4G1
STK36
CAMKK1
ADIPOR2
PAQR3
GLI1
MRPL10
FAU
RPS6KB1
RPS14
RPS16
RPS18
RPS6
ATE1
RPS28
RPS29
UPF1
PABPC1
PAN2
RPS26
RPL17
TIAL1
ATXN2
SETD4
UPF2
SETD3
CCDC110
ZFP36
SETD6
RPS25
SPNS3
OSBPL8
STK11 stem
COX4I1
SLC25A26
COX5B
CHCHD2
LDHD
EXOC5
CCN2
AP3D1
WDR20
CENPK
PDHA1
CCDC66
DLAT
MCAT
ACACB
SLC35F5
TKTL1
RPE
MLX
STK11
PFKM
MXD4
UTRN stem
MRPS23
ATG12
ATG7
TKFC
MAVS
MARCHF5
SDHAF2
MFN2
UTRN
USP21
USP16
USP30

**Supplementary Figure S20. TransposeNet topological architectures of 17 stems of the network of αS-modifier genes.**

### Figure S21

A.

#### αS proteotoxicity network

***LRRK2-SNCA* vesicle trafficking & protein homeostasis stem**

**Category:** damaging missense; Bonf. p: **2.17E-02**

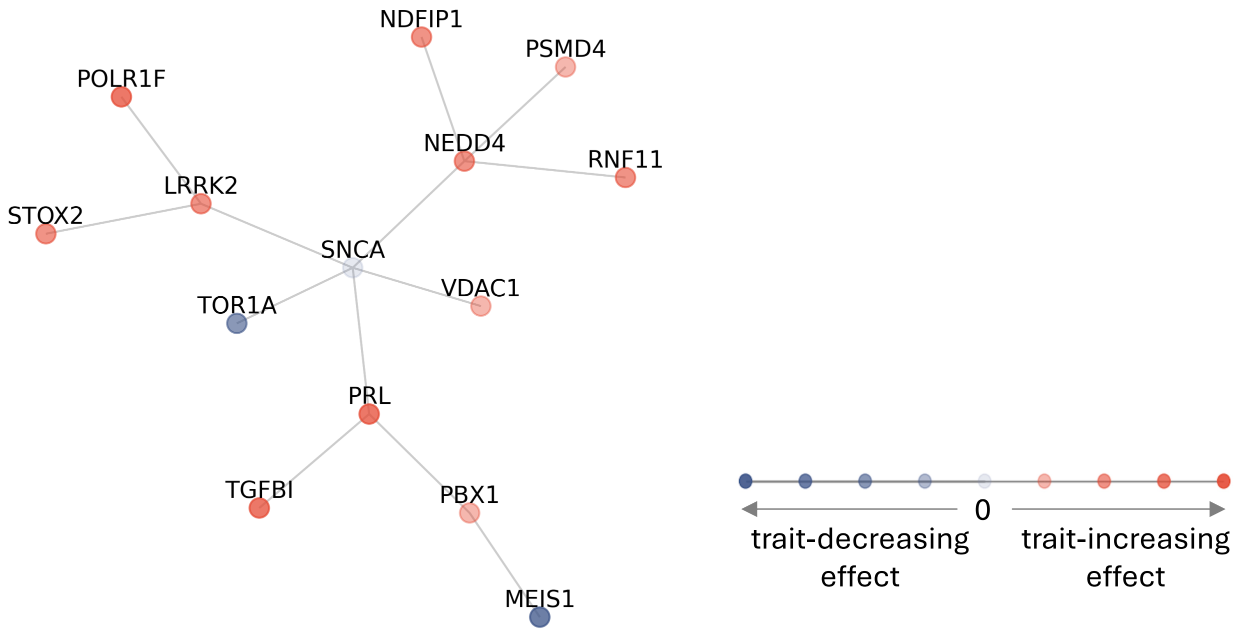


B.

| Gene symbol | GWAS hit | Known rare variant hit | Role relevant to PD |
|---|---|---|---|
| PSMD4 | X | X | Ubiquitin-proteasome system, protein degradation |
| NEDD4 | X | X | Ubiquitin ligase, α-synuclein degradation |
| LRRK2 | ✓ | ✓ | Key PD risk gene, kinase activity, neuroinflammation |
| SNCA | ✓ | * | Major component of Lewy bodies, α-synuclein aggregation |
| MEIS1 | X | X | Dopaminergic neuron development |
| RNF11 | X | X | Ubiquitin signaling, neuronal survival |
| POLR1F | X | X | Ribosomal RNA synthesis, neuroprotection |
| STOX2 | X | X | Neurodevelopment, oxidative stress response |
| NDFIP1 | X | X | protein degradation regulation, α-synuclein clearance |
| PRL | X | X | Neuroprotective effects, dopamine regulation |
| TOR1A | X | X | ER-associated protein folding, dystonia |
| PBX1 | X | X | Neural differentiation, dopaminergic neuron maintenance |
| TGFBI | X | X | Neuroinflammation, extracellular matrix regulation |
| VDAC1 | X | X | Mitochondrial function, apoptosis regulation |

**Supplementary Figure S21. Additional gene-level information on NERINE's screen-wide significant finding in the *α-synuclein* proteotoxicity screen. A.** The *LRRK2- and SNCA*-containing *vesicle trafficking and protein homeostasis*-related subnetwork of αS-modifier genes showed a screen-wide significant burden of rare damaging missense variants in AMP-PD and UKBB datasets. Screen-wide significance of networks was determined by applying Bonferroni correction over the Fisher combined p-values using the number of gene modules tested in the screen ($t_{eff}$ = 17). Here, trait-increasing effects are represented by shades of orange and trait-decreasing effects are represented by shades of purple, as shown on the scale. A darker color means a more pronounced effect in each direction. The edges in the TransposeNet module in **A** represent binary relationships and are therefore colored in grey. **B.** Additional information on αS-modifier genes in the *LRRK2-SNCA* containing *vesicle trafficking and protein homeostasis* module. Information on the member genes from previous human genetics studies is shown in the table. Only *LRRK2* had converging signals from common and rare variants, as well as from Mendelian genetics. Although *SNCA* was not a hit in population-based rare variant association tests, linkage studies identified rare Mendelian variants at this locus (as indicated by * in the table). We extracted information about member genes based on germline genetics from the GWAS catalog (https://www.ebi.ac.uk/gwas/) and the Genebass[33], SAIGE-GENE+[19], and PD-specific rare variant studies[26,27]. Annotations indicating known roles of genes relevant to PD and neurodegeneration were derived from SynGO (https://www.syngoportal.org/).

### Figure S22

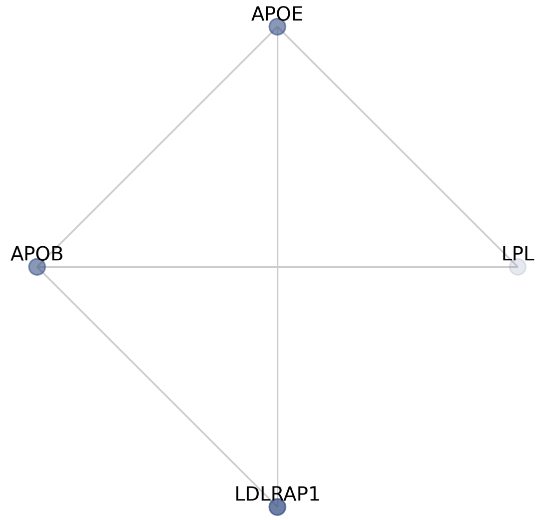


LDL core
Original p-value (with LDLR & PCSK9): 8.2102e-35
p-value (without LDLR & PCSK9): 4.1412e-9
Gene LLR (without LDLR & PCSK9): 38.95

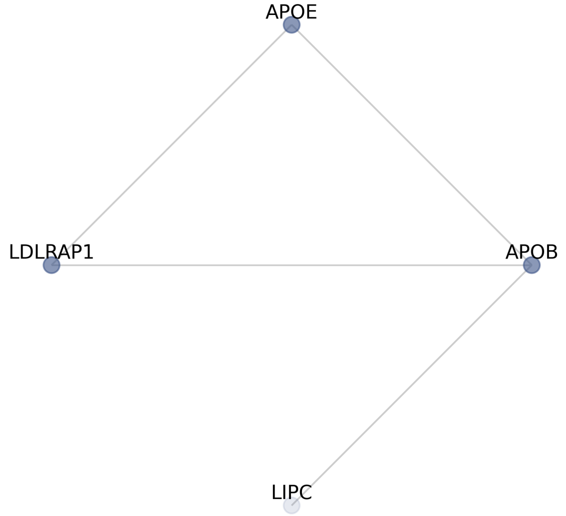


REACTOME CHYLOMICRON CLEARANCE
Original p-value (with LDLR & PCSK9): 1.0927e-9
p-value (without LDLR & PCSK9): 7.1256e-9
Gene LLR (without LDLR & PCSK9): 36.09

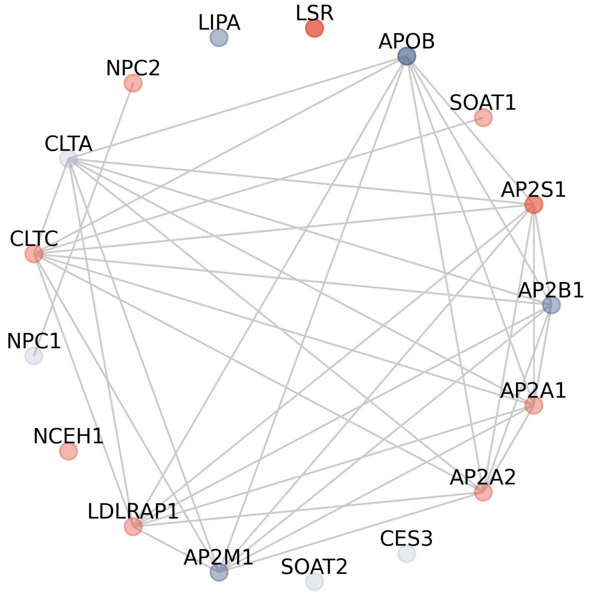


REACTOME LDL CLEARANCE
Original p-value (with LDLR & PCSK9): 5.5273e-27
p-value (without LDLR & PCSK9): 1.9755e-7
Gene LLR (without LDLR & PCSK9): 44.14

**Supplementary Figure S22. Sensitivity analysis showing lipid-related networks with significant LoF variant burden identified by NERINE for the high LDL-C vs low LDL-C phenotype in UKBB after removing *LDLR* and *PCSK9*.** Sensitivity analysis was performed by eliminating *LDLR* and *PCSK9* from the networks with significant LoF variant burden for the "high LDL-C vs low LDL-C" phenotype in UKBB. Despite removing *LDLR* and *PCSK9*, two large-effect genes, the module of core LDL-related genes, along with LDL clearance and chylomicron clearance pathways, remained significant after Bonferroni correction.

### Figure S23

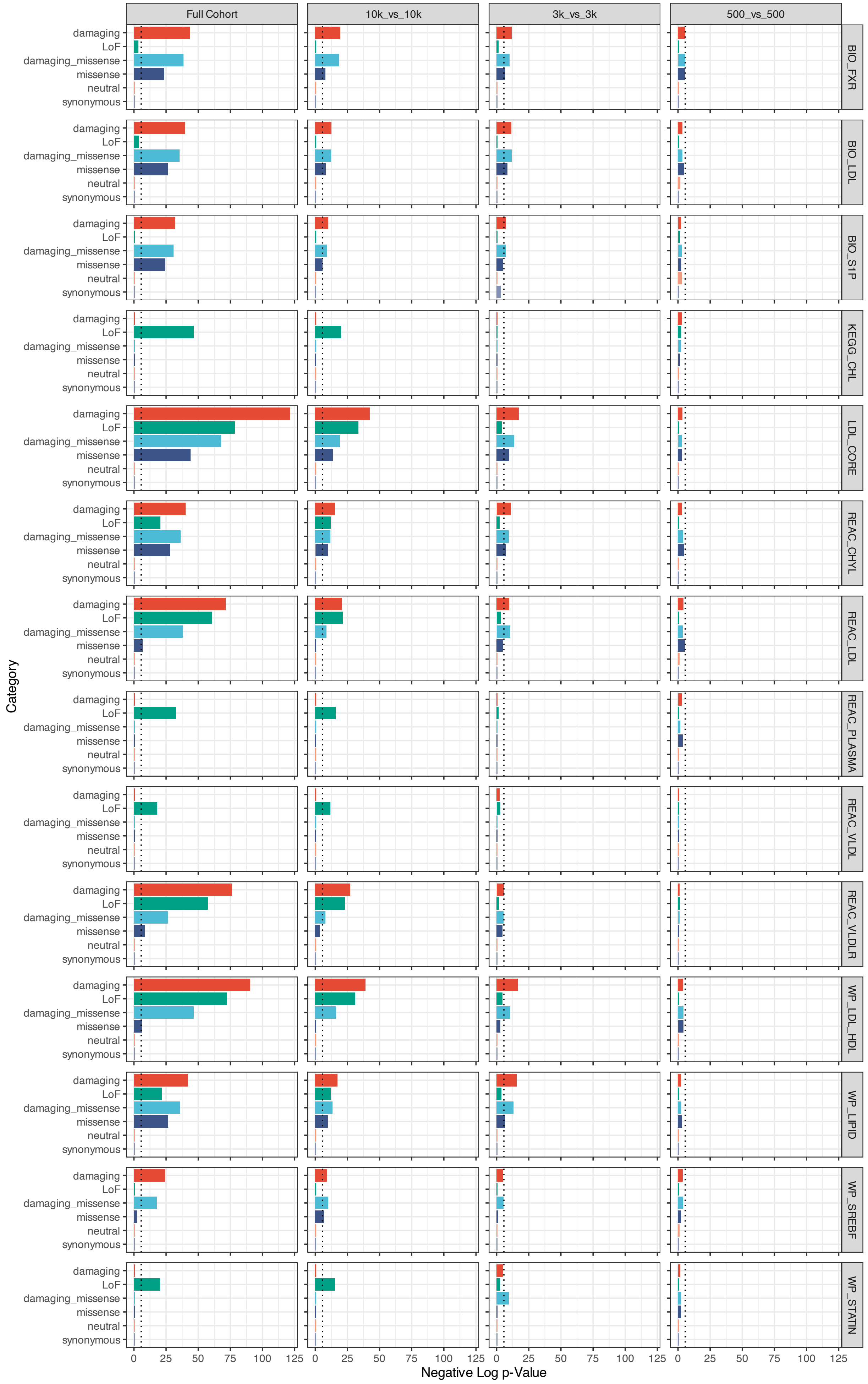

Full Cohort
10k_vs_10k
3k_vs_3k
500_vs_500
damaging
LoF
damaging_missense
missense
neutral
synonymous
BIO_FXR
BIO_LDL
BIO_S1P
KEGG_CHL
LDL_CORE
REAC_CHYL
REAC_LDL
REAC_PLASMA
REAC_VLDL
REAC_VLDLR
WP_LDL_HDL
WP_LIPID
WP_SREBF
WP_STATIN
Category
0 25 50 75 100 125
Negative Log p-Value

**Supplementary Figure S23. Performance of NERINE in downsampled cohorts for the high LDL-C vs low LDL-C phenotype in UKBB.** The high LDL-direct vs. low LDL-direct cohort was downsampled at different case-control ratios (1/3, 1/10, and 1/60). For each downsampled cohort, NERINE was competitively applied across the pathway database. We recovered a significant rare damaging variant burden in most of the lipid-related pathways that were significant in the analysis of the original cohort. For a cohort with as few as 500 cases and 500 controls, most top pathways showed nominal significance in the functional categories (LoF, damaging, and damaging missense) without inflation in the neutral missense and synonymous categories. The dashed line represents database-wide Bonferroni-corrected p-value cuto4 of 0.05. Here, BIO_FXR = BIOCARTA FXR PATHWAY, BIO_LDL = BIOCARTA LDL PATHWAY, BIO_S1P = BIOCARTA S1P PATHWAY, HDL_CORE = HDL core, KEGG_CHL = KEGG CHOLESTEROL METABOLISM, LDL_CORE = LDL core, REAC_CHYL = REACTOME CHYLOMICRON CLEARANCE, REAC_LDL = REACTOME LDL CLEARANCE, REAC_PLASMA = REACTOME PLASMA LIPOPROTEIN CLEARANCE, REAC_VLDL = REACTOME VLDL CLEARANCE, REAC_VLDLR = REACTOME VLDLR INTERNALISATION AND DEGRADATION, WP_LIPID = WP COMPOSITION OF LIPID PARTICLES, WP_LDL_HDL = WP METABOLIC PATHWAY OF_LDL HDL AND TG INCLUDING DISEASES, WP_SREBF = WP SREBF AND MIR33 IN CHOLESTEROL AND LIPID HOMEOSTASIS, and WP_STATIN = WP STATIN PATHWAY.

### Figure S24

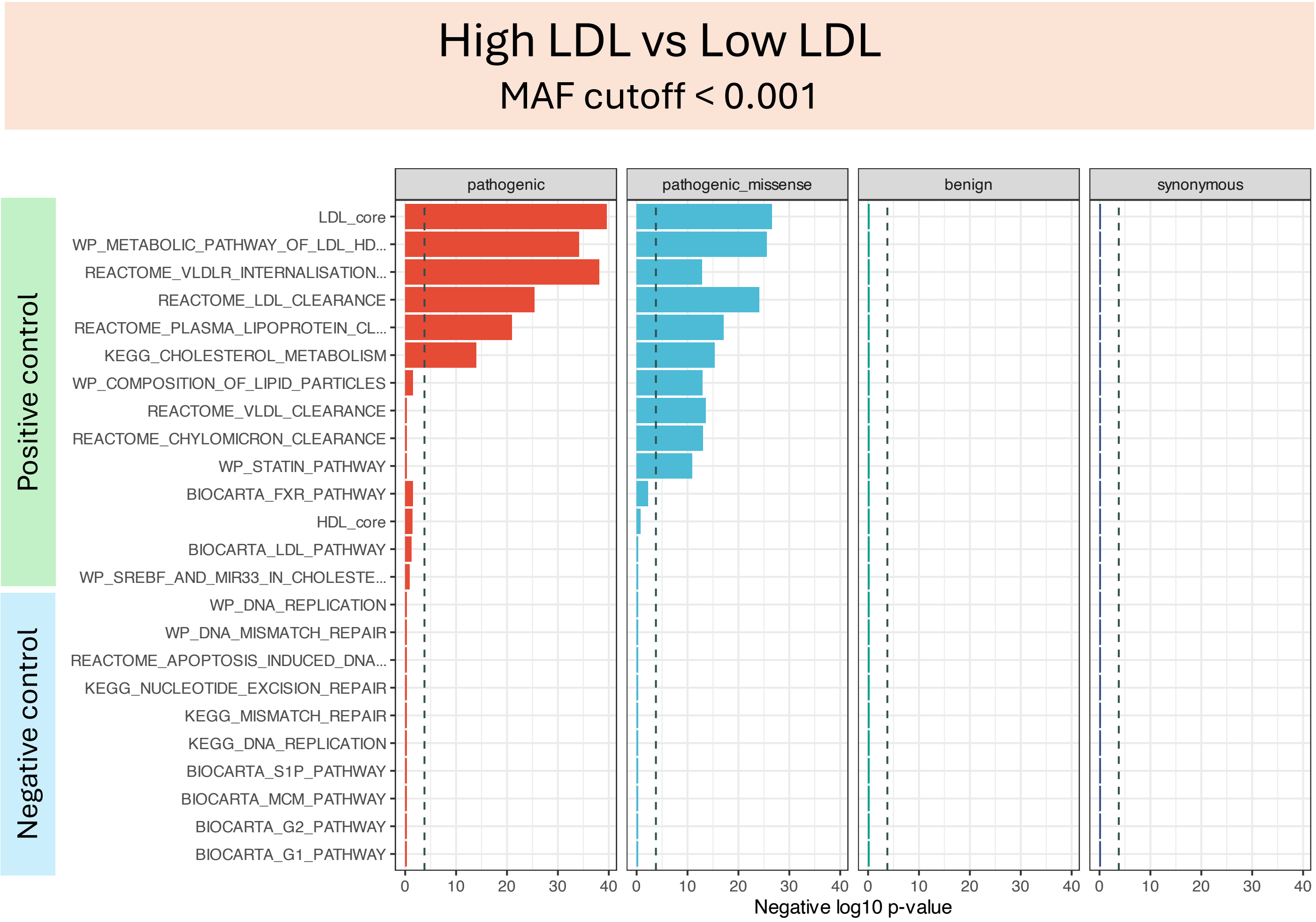
High LDL vs Low LDL
MAF cutoff < 0.001
Positive control
Negative control
pathogenic
pathogenic_missense
benign
synonymous
LDL_core
WP_METABOLIC_PATHWAY_OF_LDL_HD...
REACTOME_VLDLR_INTERNALISATION...
REACTOME_LDL_CLEARANCE
REACTOME_PLASMA_LIPOPROTEIN_CL...
KEGG_CHOLESTEROL_METABOLISM
WP_COMPOSITION_OF_LIPID_PARTICLES
REACTOME_VLDL_CLEARANCE
REACTOME_CHYLOMICRON_CLEARANCE
WP_STATIN_PATHWAY
BIOCARTA_FXR_PATHWAY
HDL_core
BIOCARTA_LDL_PATHWAY
WP_SREBF_AND_MIR33_IN_CHOLESTE...
WP_DNA_REPLICATION
WP_DNA_MISMATCH_REPAIR
REACTOME_APOPTOSIS_INDUCED_DNA...
KEGG_NUCLEOTIDE_EXCISION_REPAIR
KEGG_MISMATCH_REPAIR
KEGG_DNA_REPLICATION
BIOCARTA_S1P_PATHWAY
BIOCARTA_MCM_PATHWAY
BIOCARTA_G2_PATHWAY
BIOCARTA_G1_PATHWAY
0 10 20 30 40
Negative log10 p-value

**Supplementary Figure S24. NERINE identifies significant rare variant network burden in functional categories defined by AlphaMissense and REVEL in high LDL-C vs. low LDL-C individuals in the UK Biobank. A.** While comparing individuals with high LDL cholesterol with the ones with low LDL cholesterol in UKBB, NERINE identifies significant burden of rare (MAF < 0.001) variants in pathogenic missense (i.e., missense variants with either AlphaMissense score > 0.564 or REVEL score >= 0.664, and pathogenic (i.e., frameshifts, insertions, deletions, splice region variants, and pathogenic missenses), categories in key lipid-related pathways. The results are very similar to the analysis performed with functional variants identified by in-silico predictors, PolyPhen2 and SIFT. No significant burden of benign missense (i.e., missense variants with either AlphaMissense score < 0.34 or REVEL score < 0.5) and synonymous variants was observed. None of the cell-cycle and DNA damage repair pathways showed an enrichment in the pathogenic categories which served as a negative control. The tests were performed across our canonical pathway database. Pathway gene lists were extracted from MSigDB v7.3, and high-confidence physical and genetic interactions from protein-protein interaction (PPI) databases were used as network edges between pathway genes (see Methods).
