## Supplementary Notes for "NERINE reveals rare variant associations in gene networks across phenotypes and implicates an *SNCA-PRL-LRRK2* subnetwork in Parkinson’s disease"

**Application of NERINE to GWAS gene modules in Parkinson's Disease**

To demonstrate how NERINE can be applied to molecular networks assembled from ontological gene sets to highlight specific genes and gene modules that are amenable to more focused experimentation, we applied NERINE to six GO biological process modules significantly enriched in PD GWAS-associated genes[1-3] (**Supplementary Figure S18A**, **Supplementary Table T12**). Network modules were generated by grouping semantically similar GO biological process terms enriched in significant GWAS loci and then extracting the edge relationships of genes in each group from—(i) physical and genetic interaction databases, (ii) co-expression in substantia nigra of mid-brain (GTEx v8), and (iii) co-essentiality in CNS cell lines (DepMap 2023Q2). We tested rare variants from UKBB and AMP-PD sporadic cohorts in six categories—(i) loss-of-function (LoF), (ii) damaging missense, (iii) damaging (i.e., damaging missense and LoF), (iv) missense, (v) neutral, and (vi) synonymous. Here, LoF variants refer to frameshifts, in-frame insertions and deletions, stop-gained, stop-lost, start-lost, splice acceptors, splice donors, and splice-region variants.

NERINE identified a significant burden of rare LoF variants in the module related to *peptidyl-threonine modification* (Avg. $\hat{\theta}$ = 0.9, Bonf. p-value = 4.34e-2; **Supplementary Figure 18B**, **Supplementary Tables T13 and T14**), consistent with known kinase-phosphatase dysregulation in PD[4,5]. Co-essentiality in cells from the central nervous system (CNS) provided the most informative edge geometry for this module. Within this module, NERINE suggested trait-increasing LoF burden in *MCCC1* and *DYRK1A*, concordant with their effect on mitochondrial dysfunction[6] and DA neuron degeneration[7-9], and trait-decreasing LoF effects in *FYN* and *USP8*, concordant with preclinical evidence that their inhibition protected DA neurons[7-9] and reduced α-synuclein accumulation[10,11]. These findings highlighted candidate genes for further functional investigation. For *LRRK2*, NERINE suggested a trait-decreasing effect of LoF variants, a pattern directionally consistent with preclinical studies pointing to potential benefits of LRRK2 inhibition in PD[12,13]. Notably, however, prior genetic studies of PD found the evidence for haploinsufficiency inconclusive[14], underscoring the need for further experimental investigation.

We also explored an alternative strategy for constructing gene modules from GWAS signals. We utilized the results from the recent multi-ancestry PD GWAS[2], which applied MAGMA[15] gene set analysis and gene ontology (GO) enrichment to identify 21 significant conditionally independent GO biological process (BP) terms (**Supplementary Table T23**). For each module, NERINE was competitively applied with network topologies created from PPI, co-expression, and co-essentiality databases to test the six categories of rare variants as described above. This analysis yielded only nominally significant burdens using co-expression networks: damaging and damaging missense variants in the *regulation of* *neuronal action potential*, damaging variants in *microglial cell proliferation* and *macrophage proliferation*, and LoF variants in the *response to mitochondrial depolarization* module. None of the modules were significant after Bonferroni correction. A summary of these results is provided in **Supplementary Table T23**.

**Conditional probability of allele counts in genes in a network**

Let, the observed allele counts in gene $i$ in cases and controls be represented by two independent Poisson random variables, $X_i$ and $Y_i$, respectively, with rate parameters $\lambda_{case}^{i}$ and $\lambda_{control}^{i}$. The total allele counts for gene $i$ across the cohort is therefore distributed as $X_i + Y_i \sim Poisson(\lambda_{case}^{i}+\lambda_{control}^{i})$. The conditional probability of observing $X_i = k$ alleles in cases, given the total allele count $X_i + Y_i = n$, follows a Binomial distribution with success probability ($p$) being proportional to the ratio, $\frac{\lambda_{case}^{i}}{\lambda_{case}^{i}+\lambda_{control}^{i}}$. We assume $p$ to be a function of the gene effect $\alpha_i$ in our model.

$$
\begin{aligned}
P(X_i = k | X_i + Y_i = n) &= \frac{P(X_i = k) \times P(Y_i = n - k)}{P(X_i + Y_i = n)} \\
&= \frac{e^{-\lambda_{case}^{i}}(\lambda_{case}^{i})^{k}}{k!} \times \frac{e^{-\lambda_{control}^{i}}(\lambda_{control}^{i})^{(n-k)}}{(n-k)!} \\
&\times \frac{n!}{e^{-(\lambda_{case}^{i}+\lambda_{control}^{i})}\left(\lambda_{case}^{i} + \lambda_{control}^{i}\right)^{n}} \\
&= \binom{n}{k}\left(\frac{\lambda_{case}^{i}}{\lambda_{case}^{i} + \lambda_{control}^{i}}\right)^{k}\left(\frac{\lambda_{control}^{i}}{\lambda_{case}^{i} + \lambda_{control}^{i}}\right)^{n-k} \\
&= Binom\left(k, n, \frac{\lambda_{case}^{i}}{\lambda_{case}^{i} + \lambda_{control}^{i}}\right) \approx Binom(n, p = \phi(\alpha_i))
\end{aligned}
$$

A more appropriate modeling strategy would treat $X_i$ and $Y_i$ as independent Binomial random variables. However, under that formulation, the sum $X_i + Y_i$, no longer follows a Binomial distribution[16], and the conditional probability $P(X_i | X_i + Y_i)$ does not have a closed-form expression. Consequently, evaluating NERINE's likelihood function would require computationally intensive moment-based approximations[17], substantially increasing resource requirements and runtime. For this reason, the Poisson-based formulation described above is adopted in NERINE, providing a practical and computationally efficient approximation while preserving essential probabilistic structure.

**Custom Beta transformation for gene effects**

NERINE models the effects of genes within a network, denoted by vector $\vec{\alpha}$, as drawn from a multivariate skew-normal distribution $\vec{\alpha} \sim MSN(0, \theta \cdot \Sigma, \nu)$, where $\nu = f(N_{case}, N_{control})$. The univariate marginals follow skew-normal distributions on $(-\infty, \infty)$. When $\nu = 0$, the model reduces to a multivariate normal distribution with univariate normal marginals. Since we use each $\alpha_i$ as a proxy for the success probability parameter of a binomial distribution approximating the conditional probability of case allele counts for gene $i$ in the cohort given the total allele counts $(P(X_i = k|X_i + Y_i = n) \sim Binom(k, n, p = \phi(\alpha_i))$, we require a transformation that maps $\vec{\alpha}$ to the interval $[0,1]$.

This transformation must satisfy two conditions:

1. The mean of the transformed distribution equals $N_{case}/(N_{case} + N_{control})$.
2. The shape of the transformed distribution adapts with $\theta$: small $\theta$s (close to 0) concentrate density near the mean, while large $\theta$s shift density toward the extremes, allowing stronger gene effects with an increasingly high probability.

To achieve this, we apply a custom transformation from skew-normal to Beta distributions, rather than a probit transformation. The probit mapping, even under balanced cohorts, distorts local density around the mean, which is critical for lookup table-based likelihood calculations. Moreover, in imbalanced case-control settings, the standard probit approach is not applicable. Although skewed probit regression could theoretically achieve the mapping, it is computationally more complex involving moment-based approximations.

Let's denote the transformed gene effects by vector $\overrightarrow{\alpha'}$, where case-control imbalance is incorporated via the shape parameters of the Beta distribution. For simplicity, let's first start with a balanced case-control cohort; the univariate skew-normal marginals reduce to a scalar normals, $\alpha_i \sim Normal(\mu = 0, \sigma^2)$. In this case, we assume the transformed gene effects are denoted by $\alpha'_i \sim Beta(a, b)$. We find a mapping, $\phi: \alpha_{i_{(-\infty,\infty)}} \rightarrow \alpha'_{i_{[0,1]}}$ preserving the ordering of data points so that $(\alpha_i)_p < (\alpha_i)_q \Longrightarrow (\alpha'_i)_p < (\alpha'_i)_q$:

$$\alpha'_i = F^{-1}_{\alpha'_i}\left(\Phi\left(\frac{\alpha_i - \mu}{\sigma}\right)\right) = F^{-1}_{\alpha'_i}\left(\Phi\left(\frac{\alpha_i}{\sigma}\right)\right)$$

Here, $F_{\alpha'_i}$ is the cumulative distribution function (CDF) of $\alpha'_i$ and $\Phi$ is the standard normal CDF. In the balanced case, the Beta mean is 0.5, implying $a = b$. For $a > 1$, the distribution is bell-shaped; for $a < 1$, it is U-shaped, both symmetric around 0.5. These two cases correspond to:

- small $\theta$ (close to 0): raw gene effects, $\alpha_i$s, cluster near zero.
- large $\theta$: raw gene effects, $\alpha_i$s, shift toward extremes.

To parameterize this dependence, we set both $a$ and $b$ proportional to the reciprocal of $\theta$. Case-control imbalance is incorporated by setting $a \propto \frac{N_{case}}{N_{control}} b$. In NERINE's current implementation, we set:

$$a = \frac{N_{case}}{(N_{case}+N_{control})\times\theta}, \text{ and } b = \frac{N_{case}\times N_{control}}{N_{case}\times(N_{case}+N_{control})\times\theta}.$$

**Figure A1. NERINE's custom Beta transformation of gene-effect marginals (two-genes case).** Gene effects, $\vec{\alpha}$ are drawn from a $\mathrm{BSN}(0, \theta\cdot\Sigma, \nu)$, with $\theta = 0.1$, $\Sigma = \boldsymbol{I}$, and $\nu = 3\frac{N_{case}-N_{control}}{N_{case}+N_{control}}$. Both balanced and imbalanced cohorts with left and right skews are shown.

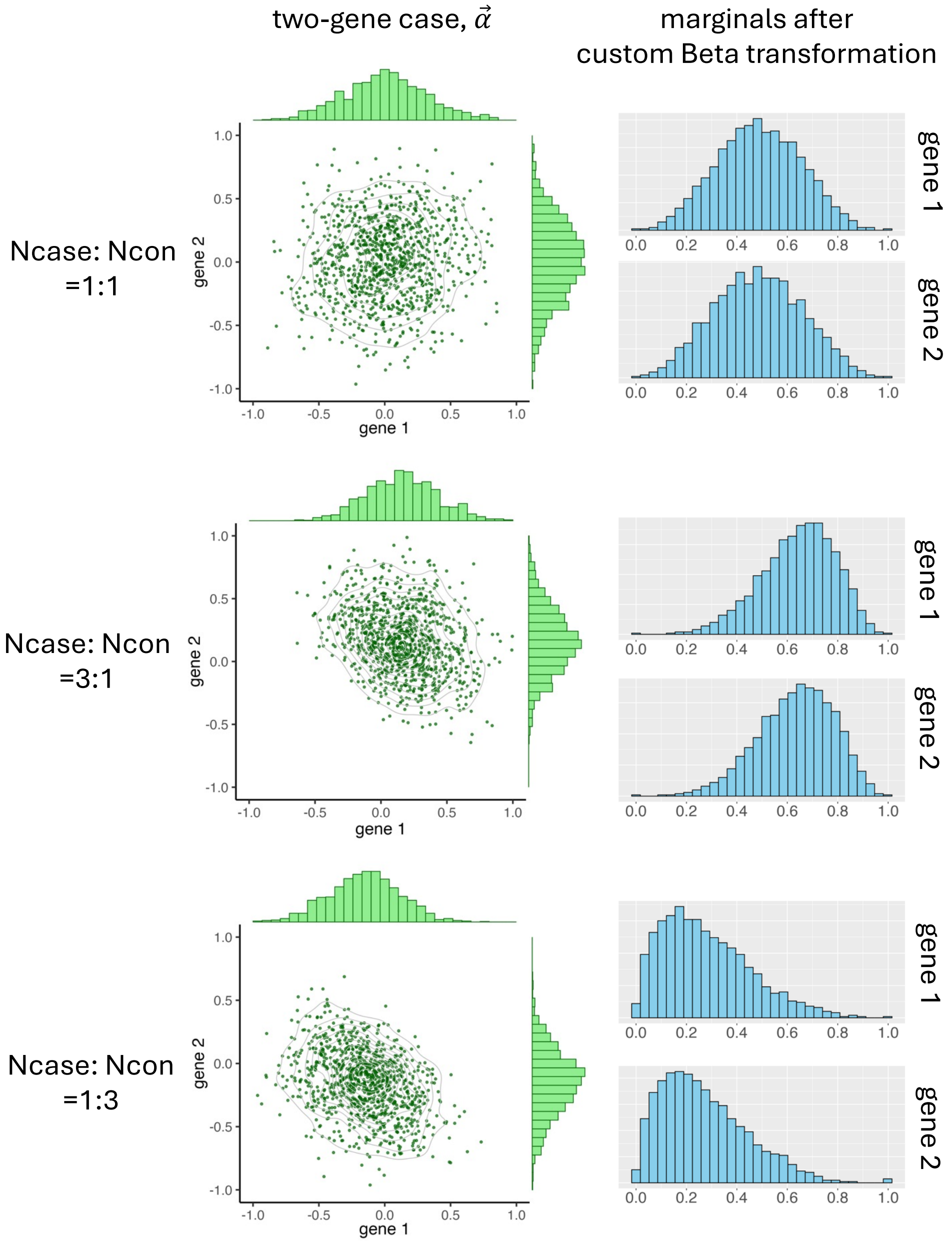

Figure A1 illustrates the custom transformation in a two-genes example, with gene effects modeled by a bivariate skew-normal distribution $\vec{\alpha} \sim BSN(0, \theta \cdot \Sigma, \nu)$. The mapping correctly shifts the mean gene effect to 0.5 in balanced cohorts, to 0.75 when cases outnumber controls 3:1, and to 0.25 when controls outnumber cases 3:1. This design also separates the treatment of skew from the lookup tables, allowing their efficient reuse in likelihood calculations.

Finally, Figure A2 compares our custom Beta transformation with the standard probit approach under a balanced two-gene cohort ($\theta = 0.1, \nu = 0, \Sigma = \boldsymbol{I}$). Our method preserves the bell-shaped structure and local density of the marginals, while the probit mapping distorts both, making it unsuitable for NERINE's marginal likelihood calculations.

**Figure A2. Comparing our custom beta transformation against standard probit transformation in two-genes case**. For gene effects $\vec{\alpha}$ drawn from $\mathrm{BVN}(0, \theta \cdot \Sigma)$, with $\theta = 0.1$ and $\Sigma = \boldsymbol{I}$, our transformation preserves the desired properties of shape and local density needed for NERINE's likelihood calculation. In contrast, the probit transformation (computed using `scipy.stats.norm.cdf` with a mean of 0 and scale of $\sqrt{0.1}$) distorts the local densities, and is therefore unsuitable for our purposes.

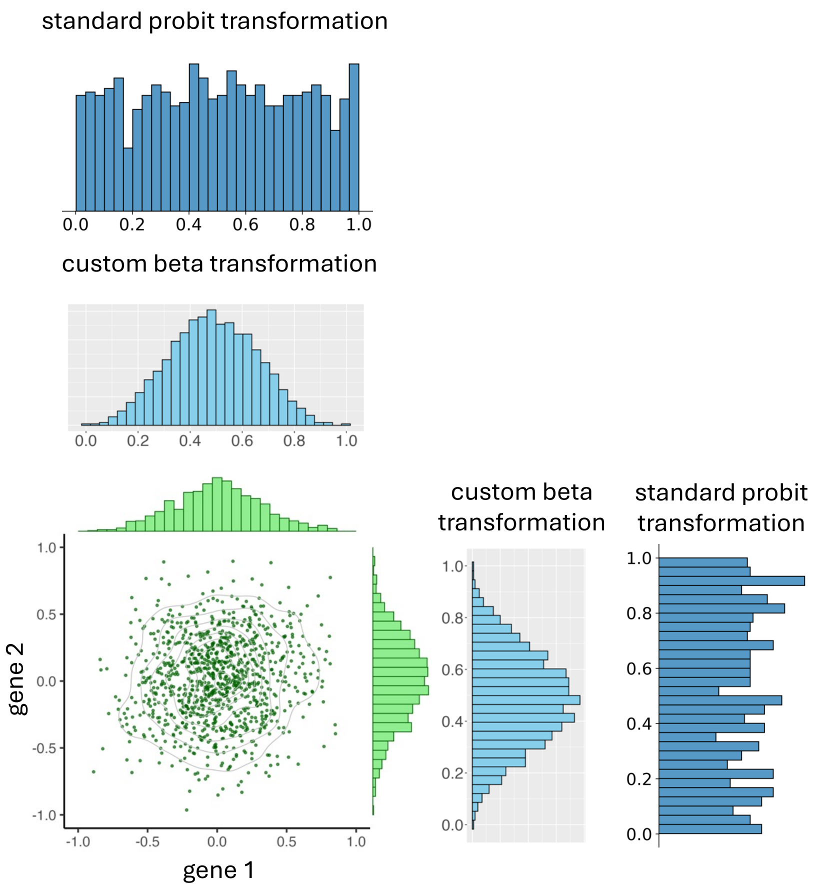

**Computing the likelihood of network effect using a lookup table with pruning**

NERINE infers the network-effect ($\theta$) on a dichotomous phenotype using the maximum likelihood estimation (MLE) framework, where the likelihood is given by,

$$L(\theta|\boldsymbol{X}, \mathbf{Y}, \vec{\alpha}, \Sigma, N_{case}, N_{control}) = \int \left( \prod_{i=1}^{m} P(X_i|X_i + Y_i, \alpha_i) \right) P(\vec{\alpha}|\theta; \Sigma, \nu) d\vec{\alpha}$$

, where $\nu = f(N_{case}, N_{control})$.

We approximate this integral as a weighted sum over *K*-dimensional quadrature points from the domain of integration:

$$L(\theta|\boldsymbol{X}, \mathbf{Y}, \vec{\alpha}, \Sigma, N_{case}, N_{control}) \approx \sum_{\vec{\alpha}} \left( \prod_{i=1}^{m} P(X_i|X_i + Y_i, \alpha_i) \right) \cdot P(\vec{\alpha}|\theta; \Sigma, \nu)$$

The weight of each *K*-variate quadrature point is given by the product of the corresponding univariate weights, effectively sampling the function over a *K*-dimensional grid. Each quadrature point involves computing two terms. For the first term, the conditional probability of case allele counts in each gene can be approximated using the probability density function of a standard Binomial distribution as described above. To calculate the probability of network-gene effects, $\vec{\alpha}$, for a given network topology ($\Sigma$) of $m$ genes and network effect ($\theta = \theta_z$), $P(\vec{\alpha}|\theta; \Sigma, \nu)$, we use a lookup table approach with pruning.

In our general framework, $\vec{\alpha} \sim MSN(0, \theta \cdot \Sigma, \nu)$. When $\nu = 0$, $\vec{\alpha}$ follows a multivariate normal (MVN) distribution with mean 0 and univariate normal marginals. For this distribution we can adopt the Gauss-Hermite (GH) quadrature[18] approach. Here, $\theta = \theta_z = 0$ implies each gene-effect $\alpha_i$ is zero. For $\theta = \theta_z > 0$, we first sample N=10,000 points from $\vec{\alpha} \sim MVN(0, \theta_z \cdot \Sigma)$ and apply variable transformation as described above to bound the marginals between 0 and 1. The transformed gene effects are denoted by $\overrightarrow{\alpha'}$. To keep the size of the lookup table tractable we impose a $1 \times K$ grid on each $\alpha_i'$. This implicitly achieves the effect of applying Cholesky decomposition on our sampled set of points. For the one-directional version of NERINE (i.e., genes can have only trait-increasing effects), we set $K = 4$. For the bi-directional version (i.e., genes can contribute to either increasing or decreasing the trait), we set $K = 9$. From the sampled points, we determine the weightings of the combinations of $\alpha_i'$s. We prune $\alpha_i'$ combinations with extremely low weight (i.e. weight < 1e-4). After the transformation, the approximate likelihood is given by,

$$L \approx \sum_{\overrightarrow{\alpha'}} \left( \prod_{i=1}^{m} P(X_i|X_i + Y_i, {\alpha'}_i) \right) \cdot P\left(\overrightarrow{\alpha'}|\theta; \Sigma, \nu\right)$$

We sample $\theta_z$s from a modified log-linear scale such that we have more $\theta_z$s with very small values close to zero (0) and sparsely distributed samples of $\theta_z$ as we move towards larger values. Since very large network effects (i.e., $\theta$ >1) are unlikely in practice, we restrict our search on the interval $[0, 1]$. Thus, the final size of the lookup table is set at $|\theta_z| \times K$, where $|\theta_z|$ represents the number of sampled $\theta_z$s. We can extend this multivariate quadrature setup for an MVN (balanced cohort) to compute the general MSN expectations almost for free by reweighting the MVN nodes[19] based on the skew parameter $\nu$. In our case, the reweighting is effectively achieved through the custom Beta transformation of the marginals as described above. This approximation scheme performs reasonably well in practice.

**Variant and sample quality control**

For each dataset, we retained high-quality biallelic variants passing GATK best practices filters and having maximum 10% missingness. For the UKBB dataset, variant-level pre-processing was performed on the DNAnexus platform. Only variants with AQ >= 50 were considered as high-quality. For MGBBB and AMP-PD datasets, variants having depth of coverage (DP) at least 10 and mapping quality (MQ) at least 90 were included. All the variant calls were based on the GRCh38 assembly. We annotated the variants with the gnomAD minor allele frequencies (genome AF: gnomADv3 and exome AF: lifted over gnomADv2 exome AFs) and in-silico predictions of deleteriousness of the missense variants by PolyPhen2 and SIFT from the dbNSFP (v4.3a) database using the using VEP (v109). Variants termed as synonymous, missense, splice donor, splice acceptor, splice region, stop-gained, stop-lost, start-lost, frameshift, in-frame insertion, and in-frame deletion, were included in the analysis. We used six masks to group variants into functional categories: (i) *Damaging missense*: missense variants predicted to be either "P" or "D" by PolyPhen2 or "deleterious" by SIFT, (ii) *LoF*: variants labelled as splice donors, splice acceptors, splice region variants, stop-gained, stop-lost, start-lost, frameshifts, in-frame insertions, and in-frame deletions; (iii) *Damaging*: LoFs and damaging missenses, (iv) *Missense*, (v) *Neutral*: missense variants predicted to be either "B" by PolyPhen2 or "tolerated" by SIFT, and (vi) *Synonymous*.

Relatedness of individuals was calculated using *King* (v2.3.2) on all variants with MAF > 0.01. All individuals marked as related by King were excluded from our analyses. We performed ancestry analysis of the individuals with the first five genetic principal components using the *somalier* (v0.2.16) tool. Our analyses primarily focused on individuals of European ancestry. Additional sample outliers were removed based on Ts/Tv, Het/Hom ratios, and per-haploid SNV counts. Outliers were defined as samples which are +/- 3 standard deviations away from the mean. We examined the distributions of ultra-rare variants such as singletons (bi-allelic SNPs for which the alternative allele is observed exactly once in the population), doubletons (bi-allelic SNPs for which the alternative allele is observed only twice in the population) and tripletons (bi-allelic SNPs for which the alternative allele is observed only thrice in the population) in all the retained samples to ensure that their distribution follows the binomial expectation. This test enables us to examine whether the distribution of ultra-rare alleles in the case-control cohorts is driven by the underlying structure of the data. We detected no differences when evaluating the distribution of doubletons and tripletons in the AMP-PD dataset. AMP-PD showed highly inflated counts of singletons in PD cases; therefore, we excluded singletons from our analysis of the AMP-PD cohort.

**Quantification code of immunostaining data**

Images were analyzed using ImageJ Macro Software. Source code is provided below. Program 1 was used for analyzing all *prolactin* immunostaining images of CiS neurons at DIV7. Program 2 was used for analyzing the images at DIV28.

Program 1:

```
//READ THE FOLLOWING BEFORE USE:
//This macro will analyze the images and output the number of cells, and the green intensity


run("Colors...", "foreground=white background=black selection=yellow");
run("Options...", "iterations=1 count=1 black");
run("Set Measurements...", "area mean standard min integrated area_fraction redirect=None decimal=5");

var width, height, pixelscale;
var cellroi, range=5, flag=0;;
var dapi_ch=1, cell_ch=1, green_ch=2;    //you can change the channels here
var T_upper_limit=220, exclude_percent=3, min_cell_size=1000;
var value;
var name;
var Intensity;
rawdir=getDirectory("User Choose Raw Data Folder");
resultdir=getDirectory("User Choose Result Data Folder");
list=getFileList(rawdir);

print("RawFolder: "+rawdir);
print("Neighbor Range:  "+range);
print("dapiCh     cellCh      punctaCh");
print(dapi_ch+"   "+cell_ch);

print("File name  cellROI#    Total Intensity   Total Intensity/Cell Number");
```

```
for(f=0;f<list.length;f++)
 {
  run("Bio-Formats Windowless Importer", "open=["+rawdir+list[f]+"]");

  //get single focused image
  focusimage();

  //width=getWidth(); height=getHeight();
  //get positive cell outline
  selectWindow("Log");
  saveAs("Text", resultdir+ "Summary" +name+ ".csv");

 }

selectWindow("Log"); run("Close");
beep();

//get depth focus image
function focusimage()
 {
   getDimensions(width, height, channels, slices, frames);
   getPixelSize(unit, pixelscale, pixelscale);
   run("Split Channels");

     selectImage("C"+1+"-"+list[f]);
     run("Z Project...", "projection=[Max Intensity]");
     saveAs("Tiff", resultdir+list[f]+"_focusch"+1+".tif"); rename("ch"+1);
      cell();
      selectImage("C"+1+"-"+list[f]); close();
      selectImage("C"+2+"-"+list[f]);
     run("Z Project...", "projection=[Max Intensity]");
     saveAs("Tiff", resultdir+list[f]+"_focusch"+2+".tif"); rename("ch"+2);
     Intensityquant();
   run("Close All");
 }

function cell()
 {
  // get dapi ROI
  selectImage("ch1");
```

```
  run("Duplicate...", "title=ch1_copy.tif");
  run("Enhance Contrast", "saturated=0.50");
  setOption("ScaleConversions", true);
  run("8-bit");
  //run("Subtract Background...", "rolling=50");
  //run("Auto Threshold", "method=MaxEntropy white");
  //run("Auto Threshold", "method=Li white");
  //run("Auto Threshold", "method=Default white");
  //run("Auto Threshold", "method=Intermodes white"); //for clumping
  run("Auto Threshold", "method=Huang2 white");
  //run("Threshold...");
  //setThreshold(51, 255);
  setOption("BlackBackground", true);
  run("Convert to Mask");
  //run("Fill Holes");
  run("Watershed");
  run("Analyze  Particles...",  "size=100-Infinity  circularity=0.1-2.00  clear
add");

  if (roiManager("Count") > 0){
  roiManager("Save", resultdir+list[f]+"_dapiROI.zip");
  cellroi=roiManager("Count");
  selectImage("ch1_copy.tif"); close();
  roiManager("Show None");
  }
  //get strong DAPI/dead cell ROI
  selectImage("ch1");
  run("Duplicate...", "title=[ch"+dapi_ch+" strong]");
  run("Enhance Contrast", "saturated=0.50");
  run("8-bit");
  run("Subtract Background...", "rolling=50");
  setThreshold(T_upper_limit, 255);
  setOption("BlackBackground", true);
  run("Convert to Mask");

  //get cell positive
  selectImage("ch"+dapi_ch+" strong");
  run("Clear Results");
  roiManager("Measure"); count=roiManager("Count");
  tmp=0;

  for(i=0;i<count;i++)
   {
```

```
    if( getResult("%Area",i) >= exclude_percent )
     {
      roiManager("Select", i-tmp);
      roiManager("Delete");
      tmp++;
     }
   }
  run("Clear Results");
  value=roiManager("count");

 if (roiManager("Count") > 0){
      roiManager("Save", resultdir+list[f]+"_cellROI.zip");
      cellroi=roiManager("Count");
      selectImage("ch"+dapi_ch+" strong"); close();
}
      roiManager("reset");
 }


function Intensityquant(){
      selectImage("ch2");
      setOption("ScaleConversions", true);
      run("8-bit");
      run("Threshold...");
      setThreshold(20, 250, "raw");
      run("Convert to Mask");
      run("Watershed");
      run("Analyze Particles...", "size=50-Infinity circularity=0-1.00 clear
add");
      if (roiManager("Count") > 0){
      roiManager("Save", resultdir+list[f]+"_intensityroi.zip");
            }
      roiManager("reset");
      open(resultdir+list[f]+"_focusch2.tif");
      if( File.exists(resultdir+list[f]+"_IntensityROI.zip")){
            roiManager("Open", resultdir+list[f]+"_Intensityroi.zip");
            roiManager("Measure");
      }
      roiManager("Measure");
      if(isOpen("Results")){
            selectWindow("Results");
            saveAs("Text", resultdir+list[f]+"_Intensity.csv");
            }
```

```
	Intensity=0;

	for(row=0; row<nResults; row++)
	{
	Intensity = Intensity + getResult("IntDen", row);
	roiManager("reset");
}
	name=list[f];
	print(list[f]+"  "+cellroi+"  "+Intensity+"     "+Intensity/cellroi);
	run("Clear Results"); run("Close All");

}
```

<u>Program 2:</u>

*Convert NDN2 Files to OmeTiff:*

```
//This macro opens High Content .nd2 file, and split it into individual .tiff files
//Output: Individual Z-stack images

rawdir=getDirectory("User Choose Individual Data Folder");

run("Bio-Formats Macro Extensions");
file = File.openDialog("Choose raw .nd2 file");
Ext.setId(file);
Ext.getSeriesCount(seriesCount);
name=File.getName(file);

for(j=1;j<seriesCount;j++)
{
		run("Bio-Formats  Importer",  "open=["+file+"]  color_mode=Default rois_import=[ROI manager] view=Hyperstack stack_order=XYCZT series_"+j);
		run("Bio-Formats                                       Exporter", "save=["+rawdir+"/"+name+"series_"+j+".ome.tif] compression=Uncompressed");
		close();
 }
-----------------------------------------------------------------------------------------------------------------------------------------------
```

*Find live cell count and green intensity:*

```
// raw data .nd2 three channel, zstack
// dapi channel : max projection collect ROI#
// green channel : ave projection  intensity for whole frame
```

```
run("Colors...", "foreground=white background=black selection=yellow");
run("Options...", "iterations=1 count=1 black");
run("Set Measurements...", "area mean min integrated redirect=None decimal=5");

var width, height, pixelscale;
var  cellroi,  punctaroi,  neighborroi,  range,  flag=0,  minsize,  maxsize,
nuclei_method, puncta_method, rollingball;
var dapi_ch=1, cell_ch=2, puncta_ch=3;
var min_cell_size, cir;
var width, height, pixelscale;
var g_Int, r_Int, puncta_size, puncta_Int, frame_size, cell_size;

rawdir=getDirectory("User Choose Raw Data Folder");
resultdir=getDirectory("User Choose Result Data Folder");
list=getFileList(rawdir);

parameter_input();

print("RawFolder: "+rawdir);

print(minsize+"   "+maxsize+" "+min_cell_size+" "+nuclei_method+"
      "+rollingball+"   "+cir);
print("dapiCh     cellCh");
print(dapi_ch+"   "+cell_ch); print("");
print("File FrameSize   cellROI#    CellSize    GreenInt    RedInt");
print("dapiCh     cellCh");
print(dapi_ch+"   "+cell_ch); print("");
print("File FrameSize   cellROI#    CellSize    GreenInt    RedInt");

selectWindow("Log");
saveAs("Text", resultdir+"Summary.xls");
selectWindow("Log"); run("Close");


for(f=0;f<list.length;f++)
 {
  run("Bio-Formats  Importer",  "open=["+rawdir+list[f]+"]  color_mode=Default
rois_import=[ROI manager] view=Hyperstack stack_order=XYCZT");
  getDimensions(width, height, channels, slices, frames);
  getPixelSize(unit, pixelscale, pixelscale);

  // create maximum projection of each channel
  run("Split Channels");
```

```
  selectWindow("C"+dapi_ch+"-"+list[f]); run("Z Project...", "projection=[Max Intensity]");        saveAs("Tiff",        resultdir+list[f]+"_ch1max.tif"); rename("ch1max"); run("Enhance Contrast", "saturated=0.35");
  selectWindow("C"+cell_ch+"-"+list[f]);         run("Z         Project...", "projection=[Average               Intensity]");               saveAs("Tiff", resultdir+list[f]+"_ch2ave.tif");  rename("ch2ave"); run("Enhance Contrast", "saturated=0.35");

  selectWindow("C"+dapi_ch+"-"+list[f]); close();
  selectWindow("C"+cell_ch+"-"+list[f]); close();

  //width=getWidth(); height=getHeight();
  //get positive cell outline
  cell();

  // measure green
  selectWindow("ch"+cell_ch+"ave"); run("Measure");
  frame_size = getResult("Area", 0); g_Int = getResult("IntDen", 0);

  // append measurement
  //print("File   FrameSize   cellRO#     Puncta#     PunctaSize  PunctaInt
      GreenInt");
  string = list[f]+"    "+frame_size+"    "+cellroi+" "+cell_size+"
      "+g_Int+"   "+r_Int;
  File.append(string, resultdir+"Summary.xls");

  run("Close All"); roiManager("reset"); run("Clear Results");
 }

roiManager("reset");
run("Clear Results");
print("MACRO FINISHED!!!");

function parameter_input()
 {
  Dialog.create("Parameter_input");
  Dialog.addNumber("Min Cell Size:", 200);
  Dialog.addChoice("Nuclei  Threshold  Type:",  newArray("Huang  dark",  "Otsu dark",  "Default  dark",  "Triangle  dark",  "Yen  dark",  "Sahnbhag  dark", "Intermodes dark", "IsoData dark", "Li dark", "MaxEntropy dark", "Mean dark", "MinError  dark",  "Minimum  dark",  "Moments  dark",  "Percentile  dark", "RenyEntropy dark" ));
  Dialog.addNumber("RollingBall Radius (pixel):", 50);
```

```
  Dialog.addNumber("Circularity:", 0.6);

  Dialog.show();

  min_cell_size = Dialog.getNumber();
  minsize = Dialog.getNumber();
  maxsize = Dialog.getNumber();
  nuclei_method = Dialog.getChoice();
  puncta_method = Dialog.getChoice();
  rollingball = Dialog.getNumber();
  cir = Dialog.getNumber();
 }


function cell()
 {
  // get dapi ROI
  selectImage("ch"+dapi_ch+"max");

  run("Duplicate...", "title=[ch"+dapi_ch+" copy]");
  run("Enhance Contrast", "saturated=0.35");
  run("8-bit");
  run("Gaussian Blur...", "sigma=1");
  run("Threshold...");
  setThreshold(59, 231, "raw");
  setOption("BlackBackground", true);
  run("Convert to Mask");
  run("Watershed");

  run("Analyze    Particles...",    "size="+min_cell_size+"-Infinity    pixel
circularity="+cir+"-1.00 exclude clear add");
  cellroi = roiManager("Count"); cell_size=0;
  if( cellroi > 0 )
   {
    roiManager("Save", resultdir+list[f]+"_dapiROI.zip");
    roiManager("Measure");
    for(i=0; i< cellroi; i++)
     {
      cell_size = cell_size + getResult("Area",i);
     }
   }
  selectImage("ch"+dapi_ch+" copy"); close();
 roiManager("Show None"); roiManager("reset");
```

```
  run("Clear Results");
 }
```
